# Deciphering the intra-tissue specific Collagen PTMs site-specific heterogeneity in Human Adrenal extracellular-matrix

**DOI:** 10.1101/2025.10.27.684899

**Authors:** Ashutosh Joshi, Ayush Nigam, Jean Lucas Kremer, Claudimara Ferini Pacicco Lotfi, Bhaskar Mondal, Trayambak Basak

**Affiliations:** School of Biosciences and Bioengineering, Indian Institute of Technology (IIT)- Mandi, Himachal Pradesh 175075, India; Institute of Biomedical Sciences, Department of Anatomy, University of São Paulo, São Paulo, SP 05508-000, Brazil; School of Chemical Sciences, Indian Institute of Technology (IIT)- Mandi, Himachal Pradesh 175075, India; Centre for Human-Computer Interface (CHCI), Indian Institute of Technology (IIT)- Mandi, Himachal Pradesh 175075, India

**Keywords:** Human adrenal, Collagen, Mass spectrometry, Data Analysis, PTM Quantitation

## Abstract

The human adrenal is one of the pivotal glands of the endocrine system. Recently, the extracellular matrix (ECM) of the adrenal capsule and cortex was explored by dividing them into two fractions: outer (OF) and inner (IF). A significant variation in the levels of ECM proteins, including collagens, was documented. During the biosynthesis of collagen, it undergoes a plethora of PTMs exhibiting crucial roles such as cell-matrix interaction, adhesion, crosslinking, stability, etc. However, the site-specific identification and characterization of collagen PTMs remained challenging and is unknown for the human adrenal gland. We applied our in-house developed proteomics pipeline to identify several PTMS in 25 collagen chains from human adrenal-ECM. In the entire collagenome, we identified a total of 963 4-hydroxyproline (4-HyP), 201 3-hydroxyproline (3-HyP), 105 hydroxylysine (HyK), 17 galactosyl-hydroxylysine (G-HyK), and 35 glucosyl galactosyl-hydroxylysine (GG-HyK) sites. Although the site-specificity of collagen PTMs (3-HyP, HyK, G/GG-HyK) across fractions is conserved, the occupancies were different in a site-specific manner. Classically, a fully 3-hydroxylated site (P^1164^) of COL1A1 associated with osteogenesis imperfecta was found to be approximately fully hydroxylated (∼99%) across fractions. Similarly, we also looked at the microheterogeneity of lysine modifications on one lysine residue (K^862^) of COL1A1. We observed that the hydroxylation level was higher in OF, while glycosylation levels were higher in IF. This may suggest a change in the crosslinking of collagen I across both fractions. Further, our analysis revealed much higher site-specific O-glycosylation in basement membrane collagen-IV, potentially facilitating the secretion of steroids from the adrenal gland. For the first time, we have annotated the collagen PTMs, developed a COL1A1 PTM map, and quantitated site-specific PTMs in the human adrenal gland. Taken together, this work revealed that intra-tissue-specific site-specific PTM collagen heterogeneity and lay the foundation for understanding their role in region-specific functions.

## Introduction

The cortex of the human adrenal gland synthesizes and secretes essential hormones such as cortisol, androgens, and aldosterone ^1^. The gland comprises a capsule, a cortex, and a medulla. The cortex has three zones that are zona glomerulosa (ZG), zona fasciculata (ZF), and zona reticularis (ZR) ^2^. Alteration in the extracellular matrix (ECM) composition correlates with tissue structure, remodeling, and steroidogenesis. Type IV collagen, an ECM protein, facilitates the expression of hydroxy-delta-5-steroid dehydrogenase (HSD3B2) and the production of cortisol ^3^. Therefore, the intra-tissue-specific extracellular matrix (ECM) composition of the human adrenal cortex must be determined. Recently, the human (adult) adrenal cortex was profiled to enrich the current knowledge base for the matrisome. Kremer *et al.* investigated the matrisome in the outer fraction (OF) and the inner fraction, revealing collagens as the most abundant adrenal matrisome (IF) ^2^ constituent. The OF comprises capsule and ZG cells, and the ZF and ZR cells comprise the IF ^2^. However, the site-specific identification and changes in the collagen post-translational modifications (PTMs) across the OF and IF of the human adrenal ECM have remained unknown.

Collagen, a prevalent protein, is one of the main components of the ECM ^4^. During ECM remodeling, not only does the collagenome vary, but the PTMs of the collagen chains also vary ^5^. Understanding the distribution of collagen chains and their PTMs in the ECM of tissue is highly significant for pathological changes in the ECM. Type I collagen is more abundant in bone or skin than other types ^6^. Type I collagen has one triple helical and two non-collagenous domains at the N- and C-terminals ^7^. The triple helix has three polyproline type II (PPII) strands held together via hydrogen bonding ^4,8^. Each single strand has a repeat of the tripeptide motif in the form of -Xaa-Yaa-Gly- ^4,8^.

Abundantly, proline (Pro) and a post-translationally modified proline i.e., 4-hydroxyproline (4-HyP) are present at Xaa and Yaa positions, respectively ^9–11^. Rarely does another PTM, i.e., 3-HyP, occupy the Xaa position in the presence of 4-HyP in the Yaa position ^12,13^. In humans, collagenome -Pro-4-HyP-Gly- forms the most abundant tripeptide motif ^14^. The hydroxylation of proline is one of the key modifications in collagen. 4-hydroxylation and 3-hydroxylation of proline are catalyzed by prolyl 4-hydroxylases (P4Hs) and prolyl 3-hydroxylases (P3Hs), respectively ^15^. The hydroxylation and glycosylation of lysine are other key PTMs in collagens. Lysine moieties are present mainly on Yaa in the -Xaa-Lys-Gly- motif. The enzyme lysyl hydroxylases (LHs) catalyze lysyl hydroxylation to give hydroxylysine (HyK) as a product ^15,16^. The hydroxylysine can get further modified via glycosylation catalyzed by COLGALT 1 or 2 and LH3 ^16,17^. During glycosylation, one sugar (galactose) or two sugars (glucose and galactose) moieties are added to HyK to form galactosyl-hydroxylysine (G-HyK) or glucosyl galactosyl-hydroxylysine (GG-HyK), respectively ^18–22^. Modifications such as 4-HyP are essential for the stabilization of the collagen structure ^4,8^. Moreover, the alterations of lysine are helpful in the formation of stabilized collagen cross-link products ^23–25^.

Liquid chromatography-tandem mass spectrometry (LC-MS/MS) has emerged as a powerful technique to analyze the PTMs of collagen ^26,27^. Due to insolubility, non-amenability to proteolytic cleavage, and many PTMs, poor sequence coverage and identification of PTMs of collagen ^28–30^ are also affected. However, through our in-house pipeline, our lab has been able to increase the sequence coverage and locate the accurate site-specific PTM positions on collagen chains ^5,26,27,31^. Herein, we hypothesize that the collagen PTMs heterogeneously remodel the human adrenal matrisome specific to zona glomerulosa, zona fasiculata and zona reticularis. To test this hypothesis, we have used human adrenal matrisome proteomic data (PXD051801) to decipher the comprehensive site-specific adrenal ECM collagen PTMs through our in-house developed pipeline ^2^. For the first time, we analyzed the collagenome in the matrisome of the OF and IF of the adrenal cortex. We successfully generated the PTM map of the most abundant collagen I alpha 1 (COL1A1) chain and subsequently identified several collagen PTMs of 11 different collagen chains. We found 1223 collagen-PTM sites in the adrenal matrisome (OF and IF combined). Moreover, we quantitated 8 sites of 3-HyP, 6 of HyK, 1 of GG-HyK PTM in COL1A1. As collagen IV has a crucial role in hormone production in the adrenal gland (*vide supra*), we quantitated the lysine-modified sites in COL4A1 and COL4A2. We also deciphered the variations in the occupancy levels of lysine modifications in collagen I and collagen IV. In addition, we utilized the open-source online knowledgebase, ColPTMScape, to understand the conservation of site-specific PTMs and alteration in their occupancy across different human tissues. This work lays a foundation for understanding the alteration in the collagen types and the variations in the PTMs of type I collagen across the OF and IF. We believe this discovery will potentially help us understand ECM remodeling in the pathological conditions associated with the adrenal gland.

## Results and Discussion

As previously mentioned, collagen chains are highly abundant in the ECM. In the combined ECM of OF and IF, 121 proteins were identified. The study found that 18.18% of the ECM proteins represent collagen (22 collagen chains) ^2^. This signifies the importance of collagen in structural support and cell-ECM interactions ^26^.

### Identification and Relative Abundance of Collagen Chains

A database search was conducted using the publicly available ECM data for the OF and IF of the human adrenal cortex (PXD051801) to identify different types of collagen chains (Figure 1A). In a two-step re-analyze of the dataset, we identified 25 collagen chains along with 6 additional chains (COL4A3, COL4A5, COL8A2, COL9A3, COL11A1, COL11A2) but could not locate COL4A6, COL15A1, and COL21A1 (Figure 1B). Among the 25 chains, 5 and 2 collagen chains were unique to the OF and IF, respectively. Moreover, 19 collagen chains were common to OF and IF (Figure 1B). Kremer *et al*. noted that COL2A1 and COL5A3 were present in the IF, while these two chains were also found in both the OF and IF. Additionally, we discovered COL4A5, COL9A3, and COL4A3 in the IF. In the OF, Kremer *et al*. identified 3 collagen chains. However, we did not find COL4A6 in the OF fraction but did identify 3 additional collagen chains in the OF (COL8A2, COL11A2, and COL21A1). Furthermore, Kremer et al. reported that COL15A1 was present in both OF and IF, but it was not detected in our analysis. Interestingly, we observed one extra collagen chain (COL11A1) in both OF and IF.

**Figure 1.**
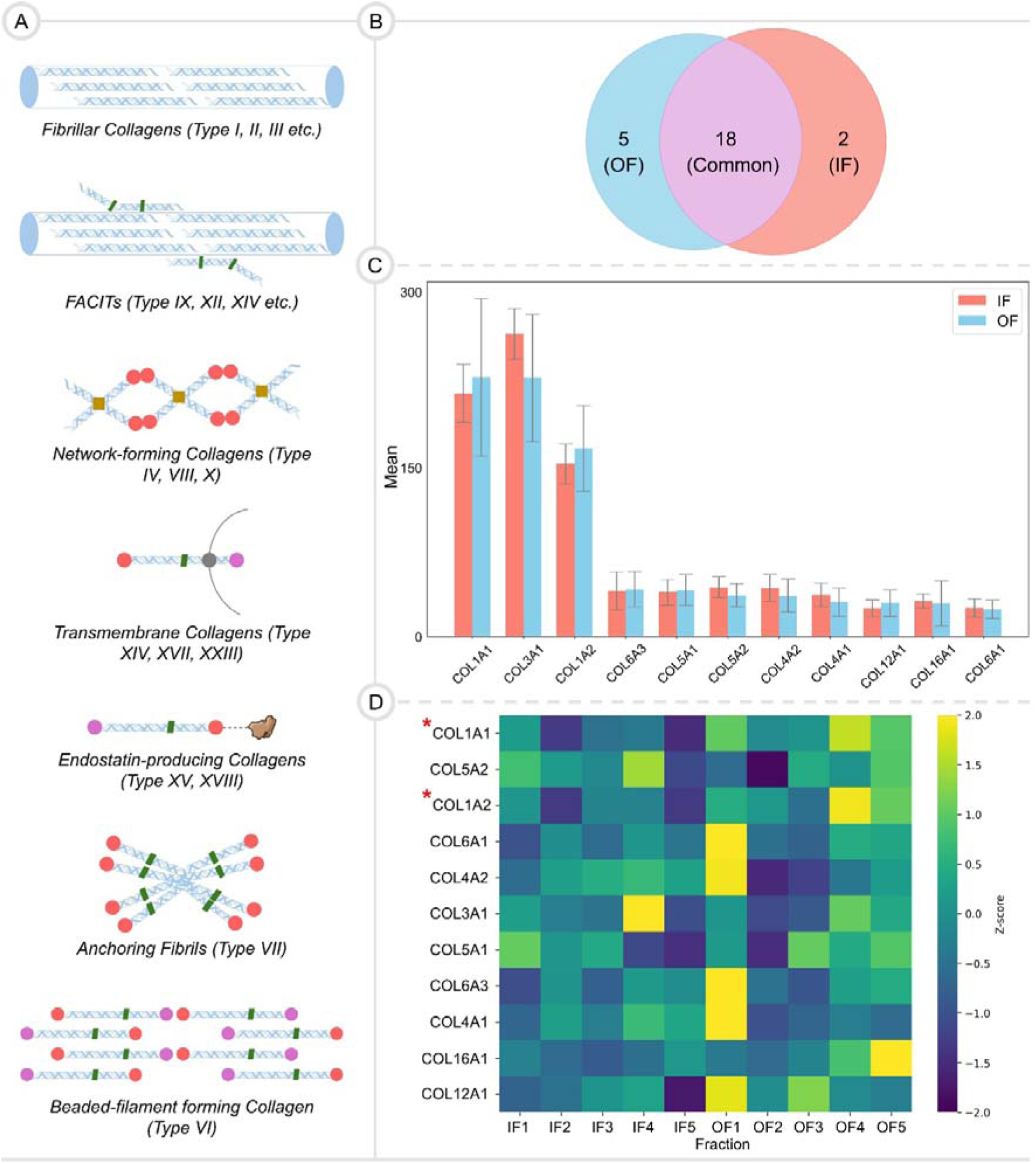
**A.** A representation illustrating the supramolecular assembly-based classification of collagens. **B.** A Venn diagram displaying the distribution of collagen chains among the OF and IF in our analysis. **C.** The bar plot shows the means and standard deviations of collagen chains for the OF and IF. **D.** The heatmap illustrates the relative abundance of collagen chains (MS^1^ quantitated) identified after the database search across the OF and IF. The correlation distance method was employed to cluster the files corresponding to OF and IF. Z-score normalization was applied to the spectral counts. Quantitation was performed for all chains, considering p < 0.05 as statistically significant.

Due to the increase in the number of discovered chains in the OF and IF, we were intrigued to look for the relative abundance of collagen chains in the OF and IF. We sorted the top 11 abundant collagen chains for OF and IF based on spectral count (Figure 1C). The descending order of collagen chains in OF: COL1A1 > COL3A1 > COL1A2 > COL6A3 > COL5A1 > COL5A2 > COL4A2 > COL4A1 > COL12A1 > COL16A1 > COL6A1 (Figure 1C). However, the descending order in IF differs and is as follows: COL3A1 > COL1A1 > COL1A2 > COL5A2 > COL4A2 > COL6A3 > COL5A1 > COL4A1 > COL16A1 > COL12A1 ∼ COL6A1 (Figure 1C). Although the value for COL1A1 was similar in both the OF and IF fractions, the other fibrillar collagen, COL3A1, was much higher in the IF fraction. In addition to this, the heatmap in Figure 1D has MS^1^-based quantitation of the top 11 collagen chains mapped from five files for both OF and IF. It was prepared by applying the Z-score normalization to quantitation data. We observed that all files in IF have fewer collagen chains than OF (Figure 1D). Two files from OF (OF1, OF5) show the higher relative abundance approximately for all collagen chains (Figure 1D).

It was observed that 5 collagen chains (COL1A1, COL1A2, COL6A3, COL5A1, and COL12A1) have higher abundance in OF, while 6 collagen chains (COL3A1, COL5A2, COL4A2, COL4A1, COL16A1, and COL6A1) in IF (Figure 1C). Among abundant 5 collagen chains, COL1A1 and COL1A2 chains displayed statistically significant abundance in OF compared to IF. As previously suggested by Kremer *et al*., the high abundance of COL1A1, COL1A2, and COL6A3 may be crucial for cell renewal in the OF of the adrenal cortex. Moreover, the higher abundance of COL4A1 in IF than OF may suggest its production requirement for HSD3B2 and cortisol (*vide supra*). As a consequence of the collagen chains relative abundance analysis, it suggests that the collagenome of both fractions is slightly varied. This prompted us to look for variations in the PTM sites of collagen chains between OF and IF.

### Quantitating the Collagen Chains in OF and IF

Including hydroxylation of proline and lysine residues in database search yields new collagen chains and provides more accurate quantitation ^5,15,27,31–34^. Classical amino acid analysis of several tissue-specific ECMs has revealed a higher abundance of hydroxyproline residues as collagens are highly modified with this PTM ^11,14,35^. Therefore, we hypothesize that incorporating the modified peptides of collagens would reflect better relative quantitation using mass spectrometry. Consequently, we anticipated a change in the fold change, particularly for the collagens among the ECM proteins showed in the previous analysis (COL1A1, COL1A2, COL6A1, COL6A3, Fibrillin-1, Heparan sulfate proteoglycan, Laminin subunit gamma 5, Laminin subunit beta 2, Laminin subunit gamma 1, Versican, Periostin, Pyruvate kinase PKM). We checked the fold change based on the MS^1^ quantitation of the proteins mentioned above and compared the values from the previous analysis. The correlation curve shows the fold change in OF, using IF as a control. We observed a significant positive correlation (r= 0.86, p=0.01) (Figure 2A, Table S1). As expected, the fold change values correlated significantly well for all the non-collagenous proteins harboring fewer PTMs (Figure 2A).

**Figure 2.**
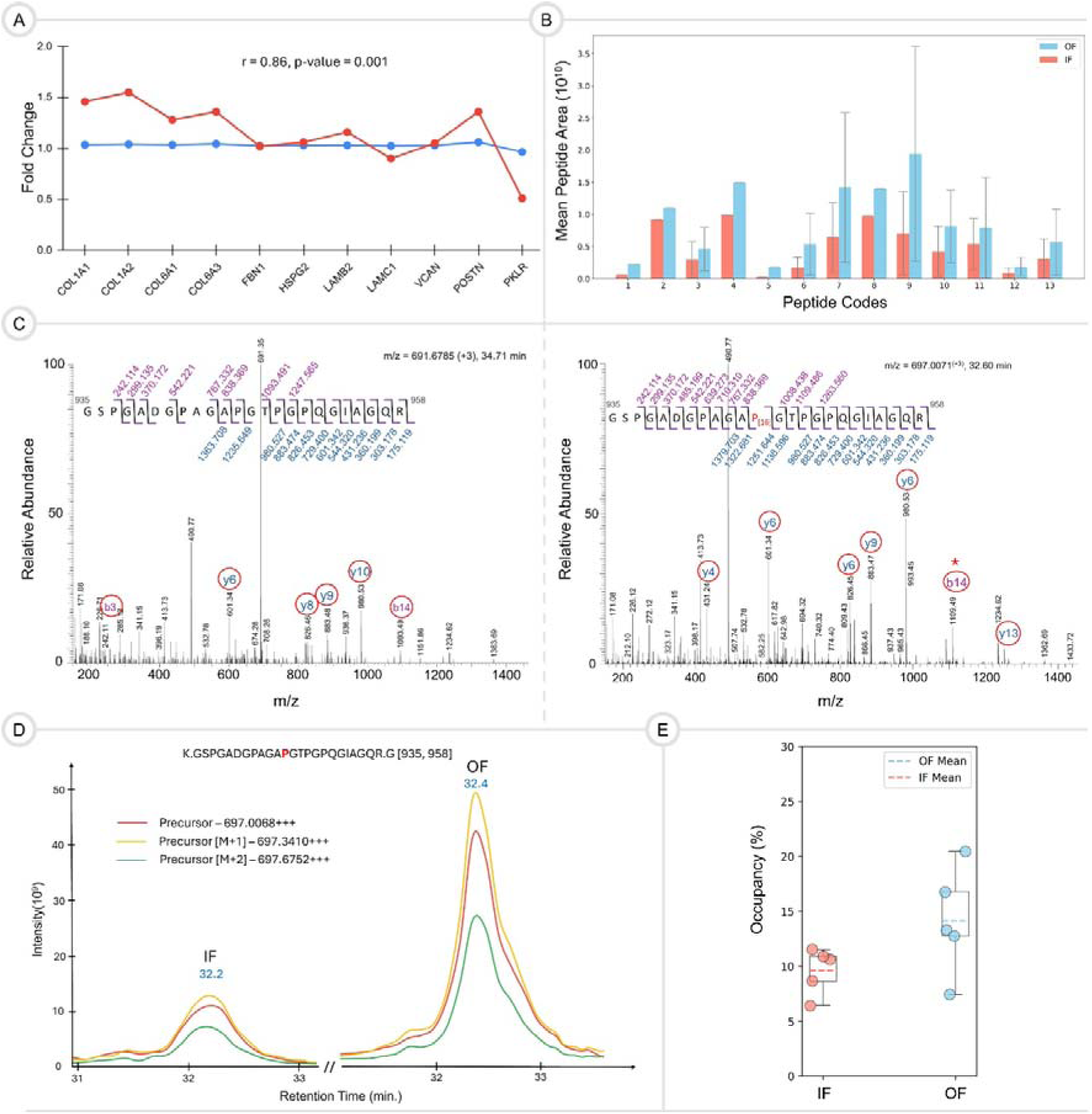
**A.** The plot represents the fold change of ECM proteins in OF compared to IF between the data analyzed by Kremer *et al.* (blue) and re-analysed data (red). **B.** The bar plot shows the peptides. The red and blue bars represent the IF and OF, respectively. The peptide 10 from OF was analyzed for PSM development and quantitation. **C.** The unmodified and modified peptide’s PSMs are shown on the left and right. **D.** The plot shows the different peptide intensities in IF and OF. **E.** The box-whisker plot represents peptide quantitation in IF and OF (non-significant, p > 0.05).

As collagen has a plethora of hydroxylation on proline and lysine as PTMs, our database search is crucial in finding the exact fold change of collagen chains in OF compared to IF by including hydroxylated collagen peptides (Figure 2A). Our analyses show an increased fold change in the collagen chains COL1A1, COL1A2, COL6A1, and COL6A3. For example, COL1A1 increased 1.5-fold compared to the 1.0-fold increase described by Kremer *et al.* (Figure 2A, Table S1).

The modified peptides (Figure 2B) were also included by including hydroxyproline in the database search, increasing the sensitivity and accuracy of collagen quantitation— consequently, the fold change value increases. In Figure 2C, we show that the peptide spectrum matches (PSM) of a 4-HyP-containing collagen-peptide, both modified and unmodified, and had the highest area (peptide 10) observed in OF. An MS^1^ peak area of this peptide from Skyline in Figure 2D highlights that though the peptide’s retention time is the same in both fractions, the area under the curve is larger in OF. As a result, we found that the abundance level of the peptide in OF is increased compared to IF, as quantitated at the MS^1^ level (Figure 2E). However, the difference was not statistically significant because the occupancy values of the peptide in two files of OF are highly sparse. As the database search through our pipeline increased the fold change of collagen chains, we became intrigued to explore the identification of site-specific PTMs in collagen across both fractions.

### Characterization of PTM Sites in Collagen and Development of PTM Maps for COL1A1 in OF and IF

As mentioned earlier, we searched for key PTM sites of collagen, such as 4-Hyp, 3-HyP, HyK, G-HyK, and GG-HyK. In OF, we found 474 4-HyP, 96 3-HyP, 51 HyK, 9 G-HyK, and 16 GG-HyK sites (Table S2). We identified 489 4-HyP, 105 3-HyP, 54 HyK, 8 G-HyK, and 19 GG-HyK sites in IF (Table S3). Figure 3A shows the modification of proline into 4-HyP and 3-HyP by different enzymes (along with their isoforms) on collagen. As 3-HyP is a rare PTM, we analyzed this PTM site across the top 11 abundant chains in both OF and IF (Figure 3B). Variations in the sites across the two fractions are discussed by examining a few chains. For instance, among the top 11 chains, COL6A3, COL12A1, and COL6A1 have only one site common in both OF and IF (Figure 3B, Table S2 and S3). In COL1A1, three exclusive prolines (P^426^, P^603^, P^690^) are 3-hydroxylated in IF (Figure 3B, Table S3). Two 3-HyP sites in IF (P^395^, P^419^) are exclusive in the case of COL1A2 (Figure 3B, Table S3). In COL3A1, six exclusive 3-HyP sites (P^334^, P^553^, P^607^, P^685^, P^700^, P^961^) are present in IF. In IF, COL4A2 has one exclusive 3-HyP site (P^975^) (Figure 3B, Table S3). The analysis supports our anticipation that there are if not significantly, a few sites that are differentially 3-hydroxylated across certain collagen chains in OF and IF.

**Figure 3.**
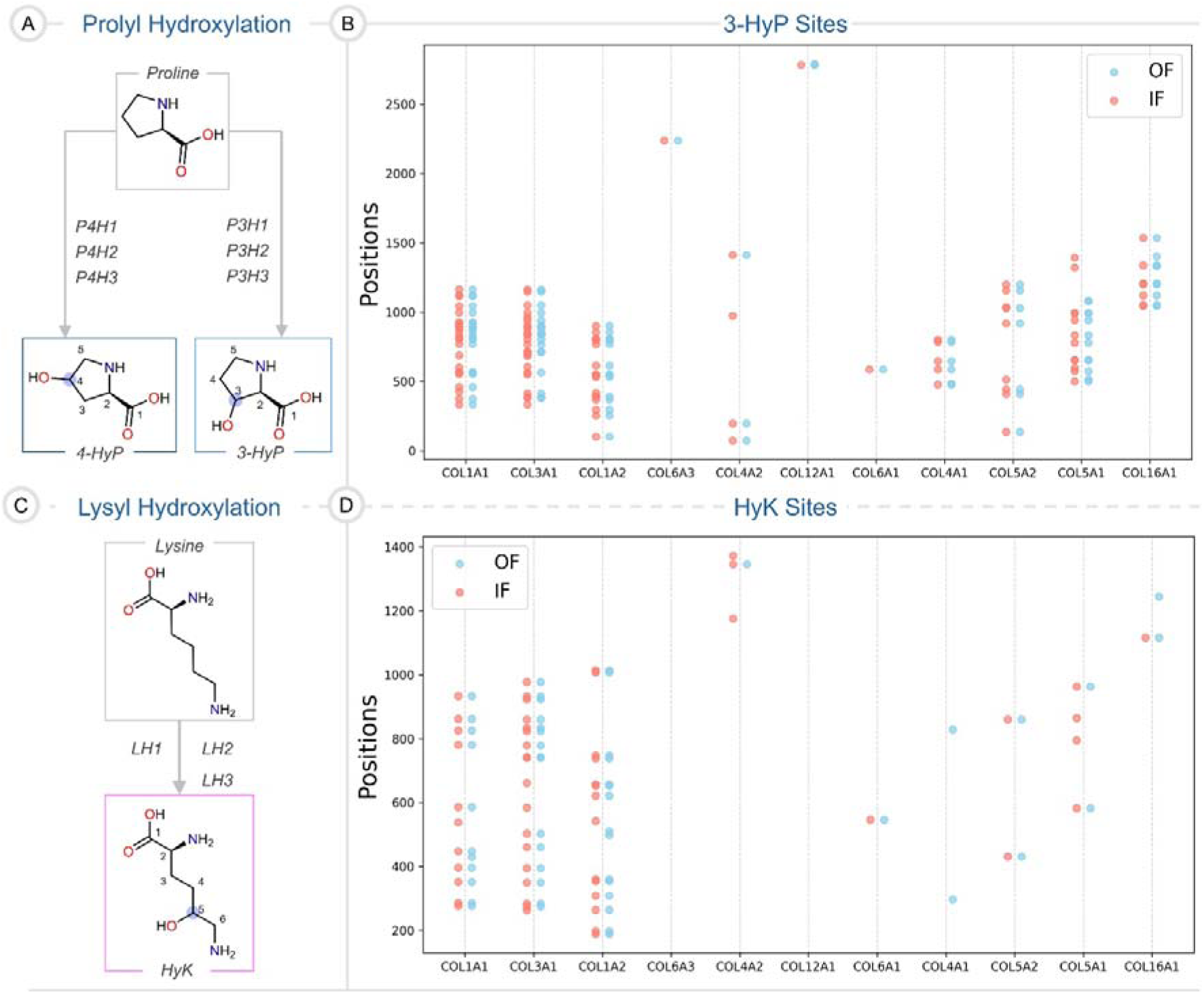
**A.** Chemical structures of prolyl hydroxylation. **B.** The bubble plot represents the positions of 3-HyP across the top 11 most abundant collagen chains in the OF and IF. **C.** Chemical structures of lysyl hydroxylation. **D.** The HyK sites across top most abundant collagen chains are shown in a bubble plot.

During collagen biosynthesis, three isoforms of lysyl hydroxylases, LH1, LH2, and LH3, can catalyse lysyl hydroxylation (Figure 3C) ^15,16^. Further, we also looked for HyK, G-HyK, and GG-HyK sites. We observed the number of sites in the top 11 collagen chains. A few of these chains are described below. The number of hydroxylated lysine residues in COL1A1, COL3A1, and COL1A2 is higher than in other collagen chains (Figure 3D). The number of HyK residues in network-forming collagen chains, COL4A1 and COL4A2, is fewer than the fibril-forming collagens (*vide supra*) (Figure 3D). Also, COL6A3 and COL12A1 have no HyK site in both fractions. Further, we checked for variation in the HyK sites between OF and IF across all collagen chains, where we came across some interesting findings. We did not observe any site in COL4A1 in IF, whereas the OF had two sites (K^298^, K^828^) (Figure 3D, Table S2). In COL4A2, we found 3 HyK sites in IF, while only 1 in OF. One site K^1346^ is common in both fractions, whereas the other two sites are unique to IF (Figure 3D, Table S3). On the contrary, in the case of COL16A1, we observed 1 site in IF and 2 in OF. Moreover, 2 additional exclusive sites are present in COL5A1 in IF, as compared to that in OF (Figure 3D, Table S2 and S3). In COL1A1, one extra lysine site is hydroxylated in each fraction, OF (K^430^) and IF (K^538^) (Figure 3D, Table S2 and S3). From the analysis, we can say there are unique HyK sites, either OF or IF.

In collagen, HyK can get further modified into G-HyK and GG-HyK by LH3 and collagen glycosyltransferases (COLGALTs) (Figure 4A) ^16,22^. Therefore, we checked for variation in glycosylation sites in both fractions (Figure 4B). The bubble plot shows that the number of GG-HyK sites is lower in fibril collagen chains compared to the network-forming collagen chains (Figure 4B). There was no HyK site in COL12A1, so we found no glycosylated HyK in this chain. However, in COL6A3, we observed 1 G-HyK and 4 GG-HyK sites in OF, while 3 GG-HyK in IF. In OF, COL1A1 has 3 G-HyK and 1 GG-HyK sites, whereas, it has 2 G-HyK and 1 GG-HyK sites in IF (Figure 4B, Table S2 and S3). In COL4A2, no G-HyK sites were observed. However, 6 GG-HyK sites in IF and 4 in OF were observed in COL4A2 (Figure 4B, Table S2 and S3). Due to the lower abundance of collagen IV, it is challenging to identify all the glycosylated sites. The analysis highlights that the PTM sites across fractions vary drastically.

**Figure 4.**
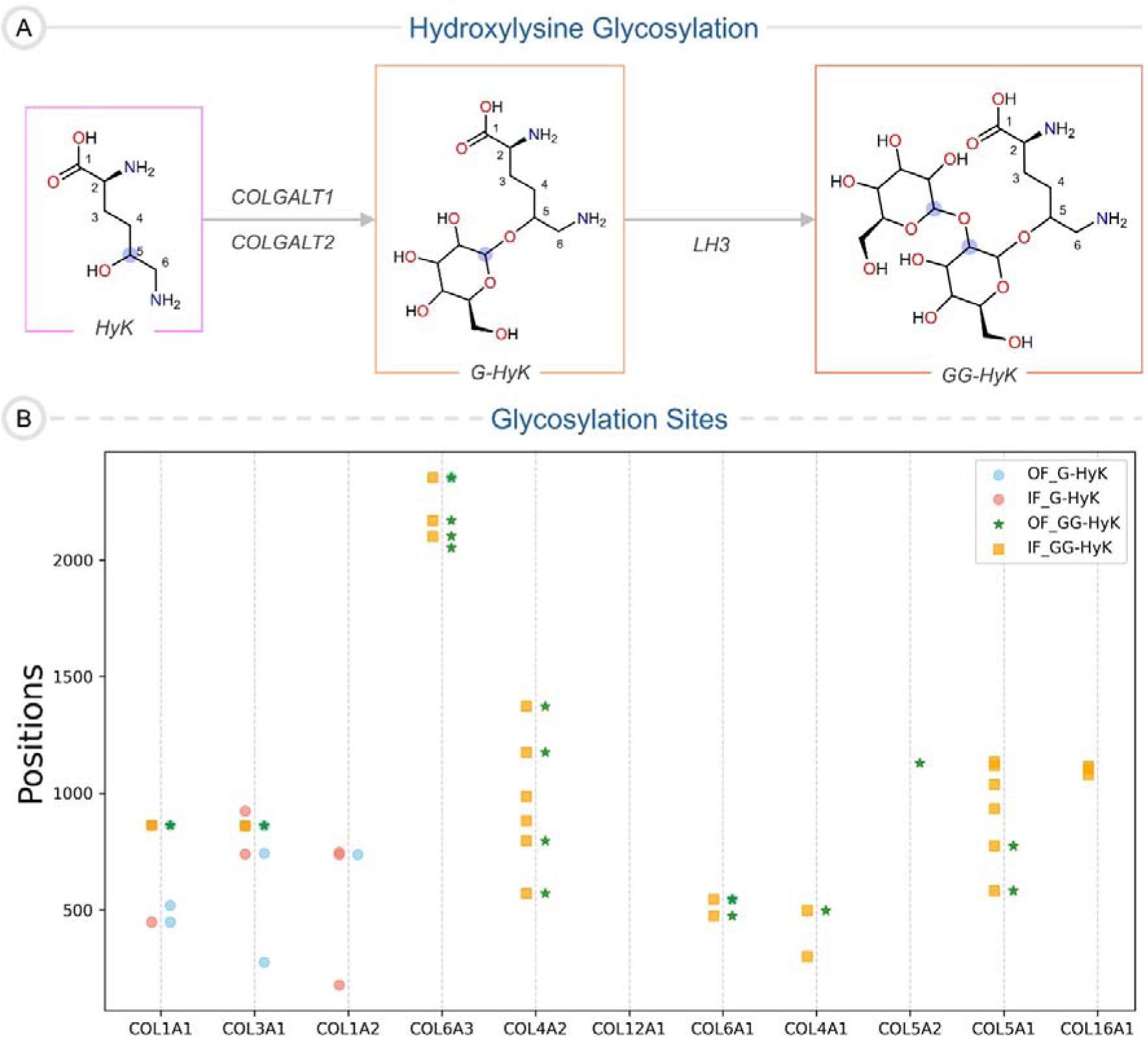
**A.** Chemical structures of galactosylation and glucosylation of a HyK residue. **B.** The bubble plot represents the positions of glycosylation sites in the most abundant collagen chains in the OF and IF.

The database search resulted in the peptides with proline and lysine modifications mapped back to the COL1A1 map. We have generated COL1A1 PTM maps for OF and IF (Figures 5A and 5B). A 72.44% and 78.31% sequence coverage were observed for COL1A1 in OF and IF, respectively. Collagen chains are polypeptides of triplet motif in the form of -Xaa-Yaa-Gly- (*vide supra*). We assigned the 4-HyP label to the modified proline present at Yaa (red P). In OF, 89 sites of 4-HyP in COL1A1 were observed (Figure 5A). Whereas the modified proline present at Xaa was assigned to 3-HyP, only if the adjacent proline was 4-hydroxylated in the -Pro-4-HyP-Gly- motif (bold red P with a blue star). A total of 21 3-HyP sites (P^333^, P^375^, P^459^, P^555^, P^567^, P^570^, P^771^, P^807^, P^816^, P^840^, P^870^, P^885^, P^894^, P^897^, P^924^, P^927^, P^996^, P^1044^, P^1119^, P^1122^, P^1164^) were mapped to COL1A1 in OF (Figure 5A). We looked for the COL1A1 PTM data of human tissues available on ColPTMScape ^36^ (https://colptmscape.iitmandi.ac.in/) to visualize the variation in the above mentioned PTMs. Of these 21, 10 3-HyP sites were the same as those observed in the PTM map of COL1A1 from human lung fibroblast ECM ^27^. These sites were P^555^, P^567^, P^690^, P^771^, P^885^, P^894^, P^897^, P^1119^, P^1122^ and P^1164^. In addition to this, 11 (P^375^, P^555^, P^567^, P^570^, P^807^, P^885^, P^897^, P^927^, P^1119^, P^1122^, P^1164^) and 1 (P^1164^) sites of 3-HyP were the same as those observed in the COL1A1 from human heart ^5^ and bone ^37^ ECM, respectively. Lysine (Lys) residue at Yaa in the -Xaa-Lys-Gly- triplet is modified to hydroxylysine (HyK), galactosyl-hydroxylysine (G-HyK), and glucosylgalactosyl-hydroxylysine (GG-HyK). We observed 11 HyK sites in COL1A1 in OF: K^277^, K^286^, K^352^, K^397^, K^430^, K^448^, K^586^, K^781^, K^826^, K^862^ and K^934^. Moreover, 3 G-HyK (K^286^, K^448^, K^862^) and 1 GG-HyK (K^862^) sites were observed. Similar to 3-HyP sites, we also checked the conservation of Lys modification sites, and we found that 9 HyK (K^277^, K^286^, K^352^, K^397^, K^586^, K^781^, K^826^, K^862^, K^934^), 1 G-HyK (K^448^), and 1 GG-HyK (K^862^) sites are the same as those in human lung fibroblast ECM. No site was common with the COL1A1 from human bone and heart.

**Figure 5.**
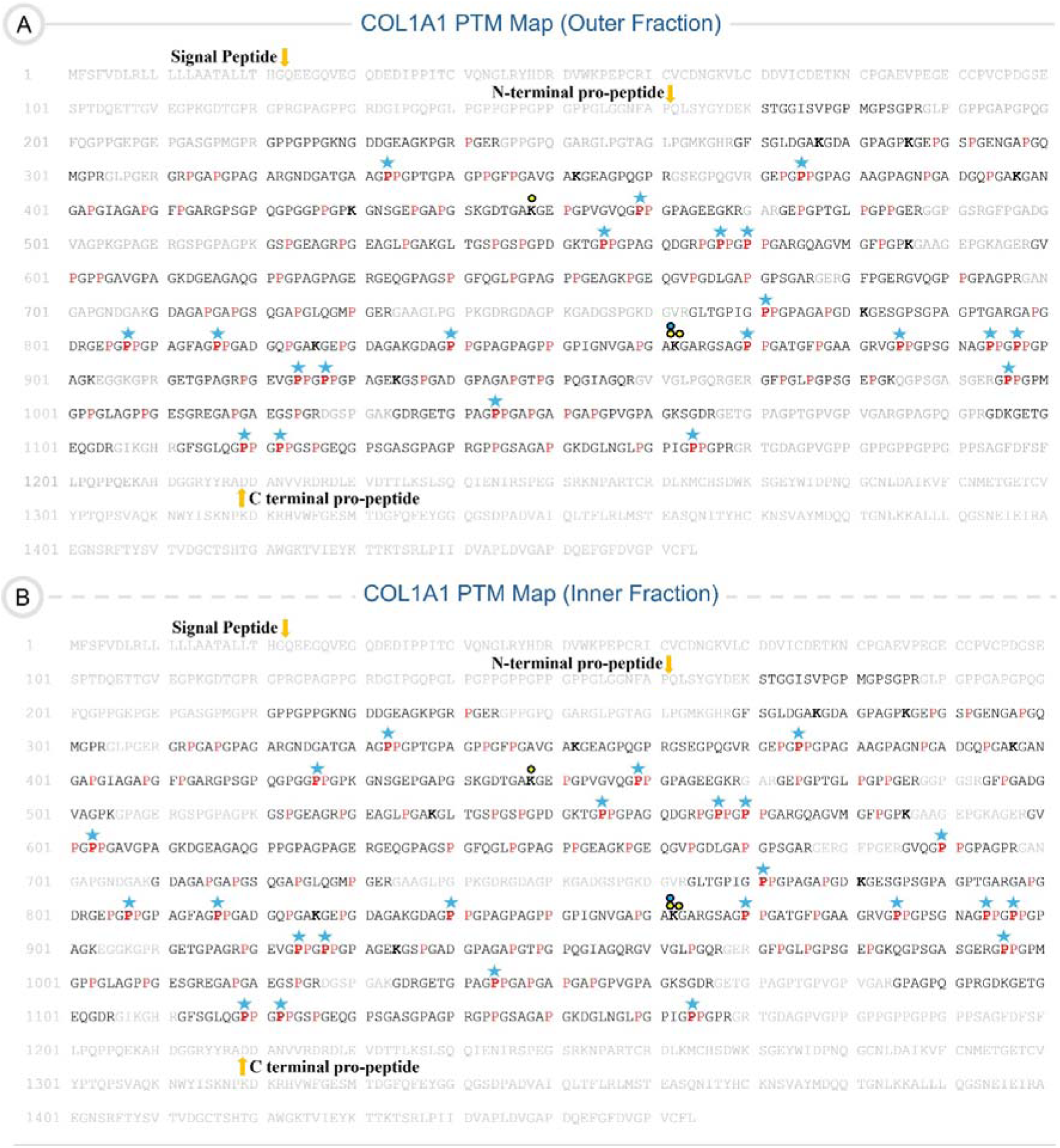
COL1A1 PTM map of the OF (**A**) and IF (**B**). The sequence coverages of OF and IF COL1A1 maps are 72.44% and 78.31%, respectively. The cleavage points for the signal peptide, N-terminal, and C-terminal propeptides are marked with the yellow arrow. The red P represents the 4-HyP, and the bold red P with a blue star highlights the presence of 3-HyP. The HyK, G-HyK, and GG-HyK are represented by bold K, bold K with yellow circles, and bold K with yellow and yellow-blue combined circles, respectively.

There are 88 4-HyP sites in COL1A1 of IF (Figure 5B). Moreover, 23 prolines at Xaa are modified to 3-HyP (P^333^, P^375^, P^426^, P^459^, P^555^, P^567^, P^570^, P^603^, P^690^, P^771^, P^807^, P^816^, P^840^, P^870^, P^885^, P^894^, P^897^, P^927^, P^996^, P^1044^, P^1119^, P^1122^, P^1164^). From -Xaa-Lys-Gly- triplet motifs, we observed 11 HyK (K^277^, K^286^, K^352^, K^397^, K^448^, K^538^, K^586^, K^781^, K^826^, K^862^, K^934^), 2 G-HyK (K^448^, K^862^), and 1 GG-HyK (K^862^) sites. While searching through the ColPTMScape knowledge base, we observed that 10 (P^555^, P^567^, P^690^, P^771^, P^885^, P^894^, P^897^, P^1119^, P^1122^, P^1164^), 11 (P^375^, P^555^, P^567^, P^570^, P^807^, P^885^, P^897^, P^927^, P^1119^, P^1122^, P^1164^), and 1 (P^1164^) sites of 3-HyP are common with COL1A1 from human lung fibroblast ^27^, heart ^5^, and bone ^37^ ECM, respectively. Moreover, 10 HyK (K^277^, K^286^, K^352^, K^397^, K^538^, K^586^, K^781^, K^826^, K^862^, K^934^), 1 G-HyK (K^448^), and 1 GG-HyK (K^862^) sites are the same as in the COL1A1 chain from human lung ^27^. In addition to COL1A1 PTM maps, we have prepared COL1A2 PTM maps for both fractions (Figure S1). We have updated the ColPTMScape knowledgebase by incorporating the collagen chains data and the PTM maps under the tissue adrenal cortex.

Additionally, we re-analyzed the rat adrenal gland MS data (PXD046828) submitted by Kremer *et al.* ^38^ to investigate the conservation of 3-HyP and HyK sites of COL1A1 within a tissue across species (human and rat). The comparative analysis revealed that human COL1A1 has 10 out of 23 (IF) and 12 out of 21 (OF) of 3-HyP sites are conserved (Figure S2). Similarly, 10 out of 11 sites of HyK in both fractions are conserved (Figure S3). The conservative nature of PTMs of collagen highlights their role in the interactions with other ECM proteins or regulation of ECM in the organ functionality.

The analysis of PTM sites of OF and IF collagen chains show the heterogeneity within a tissue similar to the quantitation of collagen chains (*vide supra*). While comparing the site-specificity of PTMs against the ColPTMScape, it also hints at conserving an extensive array of 3-HyP and Lys-modified sites across different human tissues. However, a few sites may be specific to tissue and help in the cell-ECM interaction or its functionality. Interestingly, there are variations in the modified sites of COL1A1 between OF and IF; for instance, three (P^426^, P^603^, and P^690^) sites of 3-HyP are unique to IF. This led us to two questions: (i) Will a PTM site have a different occupancy level in both fractions? (ii) Will all sites follow the same trend?

### Detecting the occupancy level of 3-HyP sites in COL1A1 of OF and IF

Although the 3-hydroxylation of proline is a rare PTM, one site (P^1164^) in collagen unmodified into 3-HyP causes the development of osteogenesis imperfecta ^39,40^. Therefore, this suggests that the occupancy of this site is different in healthy and affected individuals. As we found the variations in 3-HyP sites in COL1A1 of OF and IF, we anticipated that there would be a variation in the occupancy levels of the common 3-HyP sites. In addition to P^1164^, we tried to quantitate all common 3-HyP sites. However, due to the unavailability of all these sites in all the samples, we could quantitate only 7 sites, which are P^375^, P^570^, P^807^, P^870^, P^996^, P^1122^, and P^1164^. We quantitated all these sites using the Skyline. We observed that all sites displayed statistically non-significant difference across fractions. However,

Out of these 8 sites, the occupancy level of 2 sites (P^807^ and P^1122^) was higher in OF than in IF (Figure 6A, Table 1). The P^807^ site has 95.18 ± 4.75% occupancy in OF, whereas that is 75.46 ± 31.61% in IF. Therefore, this site is more 3-hydroxylated in the OF. Similarly, we observed that the P^1122^ site has higher occupancy in OF (42.83 ± 23.88%) compared to the IF (28.98 ± 22.61%). In human lung tissues, this site is significantly less 3-hydroxylated (9.4 ± 4.9%) ^27^. We also quantified the occupancy level of 3-hydroxylation for the P^1164^ site (^1153^DGLNGLPGPIG**P**PGPR^1168^) associated with the osteogenesis imperfecta (*vide supra*). This site is 99.69 ± 0.31% and 99.40 ± 0.64% 3-hydroxylated in the OF and IF, respectively (Figure 6A, Table 1). The occupancy level is similar across fractions. Similar to our observations, the same site is 96-99% and 42.7 ± 11% 3-hydroxylated in human bone ^37^ and lung tissues ^27^. In Figure 6B, we demonstrated two PSMs. The upper and lower ones reflect the unmodified and modified P^1164^ site-containing peptide, respectively. The mass difference of y6 ion in the modified peptide PSM distinctly highlights the modification. These results suggest two-fold inferences: (i) the occupancy of these sites can vary within a tissue-like adrenal gland, and (ii) the occupancy varies across tissues within a species.

**Figure 6.**
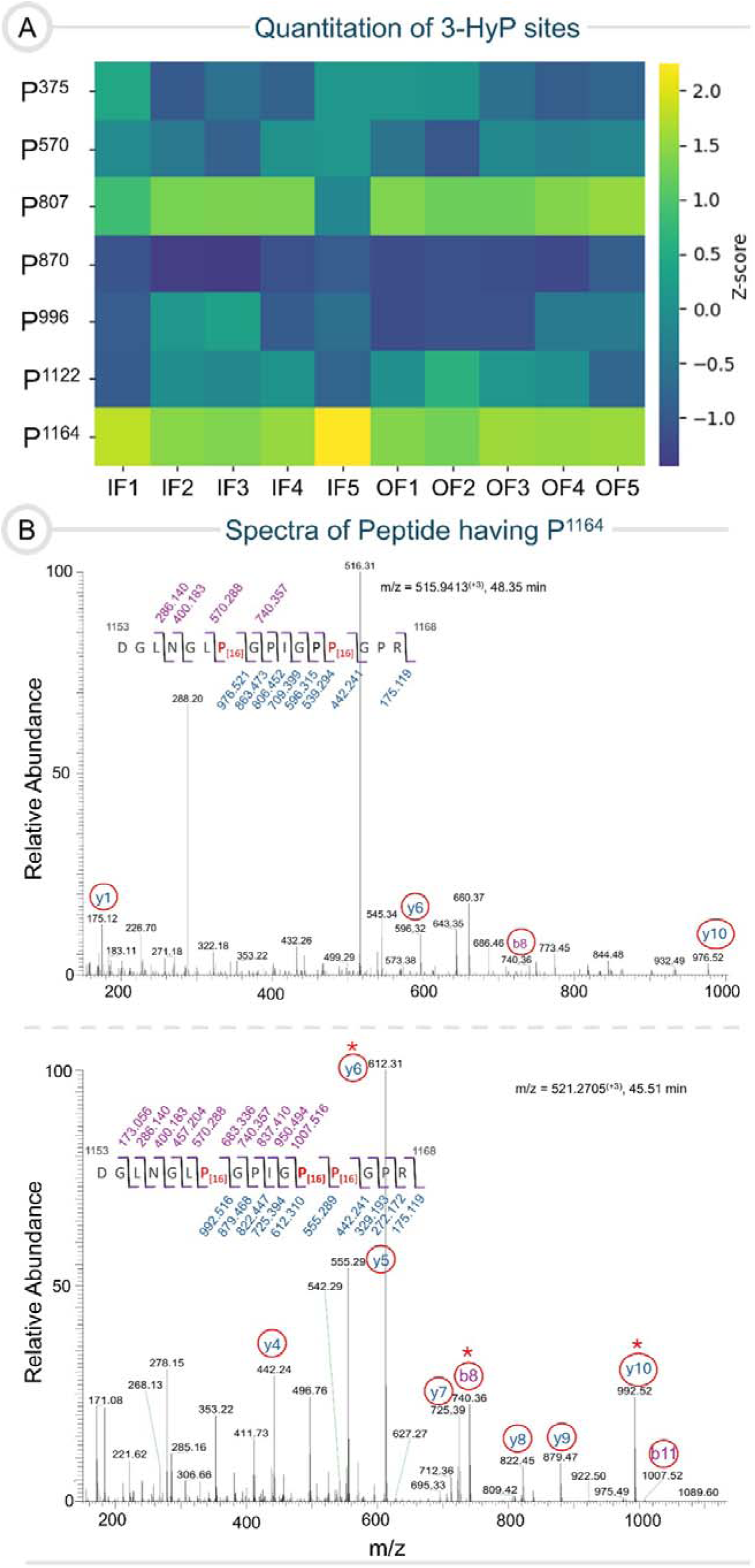
**A.** The heatmap represents the comparison between the quantitative levels of 3-HyP sites of COL1A1 across IF and OF. 3-HyP sites are mentioned on the left side of the heatmap. The range on the right side represents the occupancy of 3-HyP sites in each sample. The difference with a value of p < 0.05 is considered to be statistically significant. **B.** The peptide spectrums (unmodified and modified) for the 3-HyP site associated with the osteogenesis imperfecta are shown.

**Table 1.** Occupancy values (%) of 3-HyP in COL1A1 (*p < 0.05; ^ns^p > 0.05)

| Sites | IF | OF |
| --- | --- | --- |
| **P<sup>375</sup>** | $28.88 \pm 16.43$ | $27.43 \pm 21.48^{\text{ns}}$ |
| **P<sup>570</sup>** | $33.04 \pm 7.49$ | $25.86 \pm 13.13^{\text{ns}}$ |
| **P<sup>807</sup>** | $75.46 \pm 31.61$ | $95.18 \pm 4.75^{\text{ns}}$ |
| **P<sup>870</sup>** | $0.63 \pm 0.59$ | $0.28 \pm 0.33^{\text{ns}}$ |
| **P<sup>1122</sup>** | $28.98 \pm 22.61$ | $42.83 \pm 23.88^{\text{ns}}$ |
| **P<sup>1164</sup>** | $99.40 \pm 0.64$ | $99.69 \pm 0.31^{\text{ns}}$ |

### Quantitating the HyK and O-glycosylation of HyK sites in collagen type I, IV, and VI of OF and IF

Lysine in the Xaa-Lys-Gly tripeptide motif can be post-translationally hydroxylated, which is crucial for collagen to assemble and form a fibril ^41,42^. Therefore, it would be significant to quantify the occupancy levels of HyK sites across fractions. We performed this analysis on COL1A1. We could quantify the occupancy for 6 HyK sites (K^277^, K^352^, K^397^, K^586^, K^862^) and observed that no sites displayed the statistically significant difference. However, we detected that 2 sites have higher occupancy in OF, while the other 3 sites are more hydroxylated in IF. The K^277^ has 12.45 ± 6.09% occupancy in OF, while 5.94 ± 2.11% in IF (Figure 7A, Table 2). The occupancy of HyK^397^ in OF is 33.86 ± 34.11%, whereas it is 26.92 ± 16.82% in IF (Figure 7A, Table 2). These sites have higher occupancy levels in OF compared to IF. The K^352^ is 37.41 ± 15.54% hydroxylated in OF, while 49.23 ± 11.96% in IF (Figure 7A, Table 2). In line with previous observations, K^586^ is less hydroxylated in OF (7.97 ± 9.06%) than in IF (9.08 ± 6.88%) (Figure 7A, Table 2). Similarly, the occupancy level of the K^862^ site in OF and IF is 18.85 ± 3.81% and 22.77 ± 4.83%, respectively (Figure 7A, Table 2). The K^586^ site was also quantitated in human lung fibroblast ^27^ with an occupancy of 3.6 ± 1.1%. We have presented one unmodified (upper) and one modified (lower) peptide’s PSMs for site K^352^ (Figure 7B). Furthermore, we also quantitated one GG-HyK site (K^862^). We observed a higher occupancy value in the IF of a GG-HyK site (K^862^). The occupancy of GG-HyK^862^ in OF is 1.68 ± 0.75%, while the occupancy in IF is 2.34 ± 0.83% (Figure 8A, Table 3). This site was identified in human lung fibroblast; however, the quantitation was unavailable. The quantitation indicates the variation in the hydroxylation and glycosylation of lysine sites is possible within a tissue. Moreover, the occupancy of these sites can also vary across different tissues.

**Figure 7.**
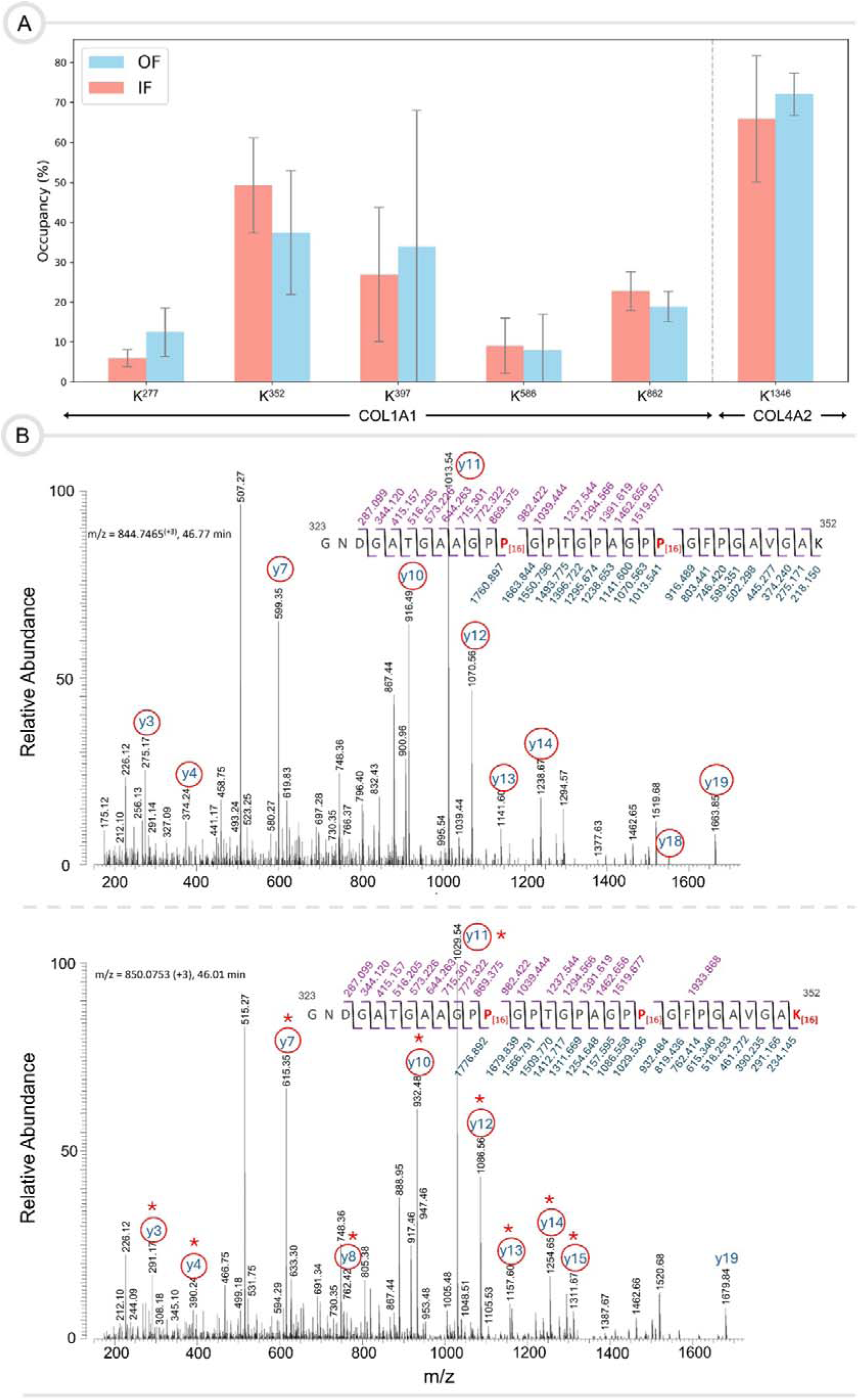
**A.** The bar plot represents the occupancy of lysine sites in COL1A1 and COL4A2. The statistically significant difference is considered if p < 0.05. **B.** The unmodified (upper) and modified (lower) PSMs of the K^352^ site are shown.

**Figure 8.**
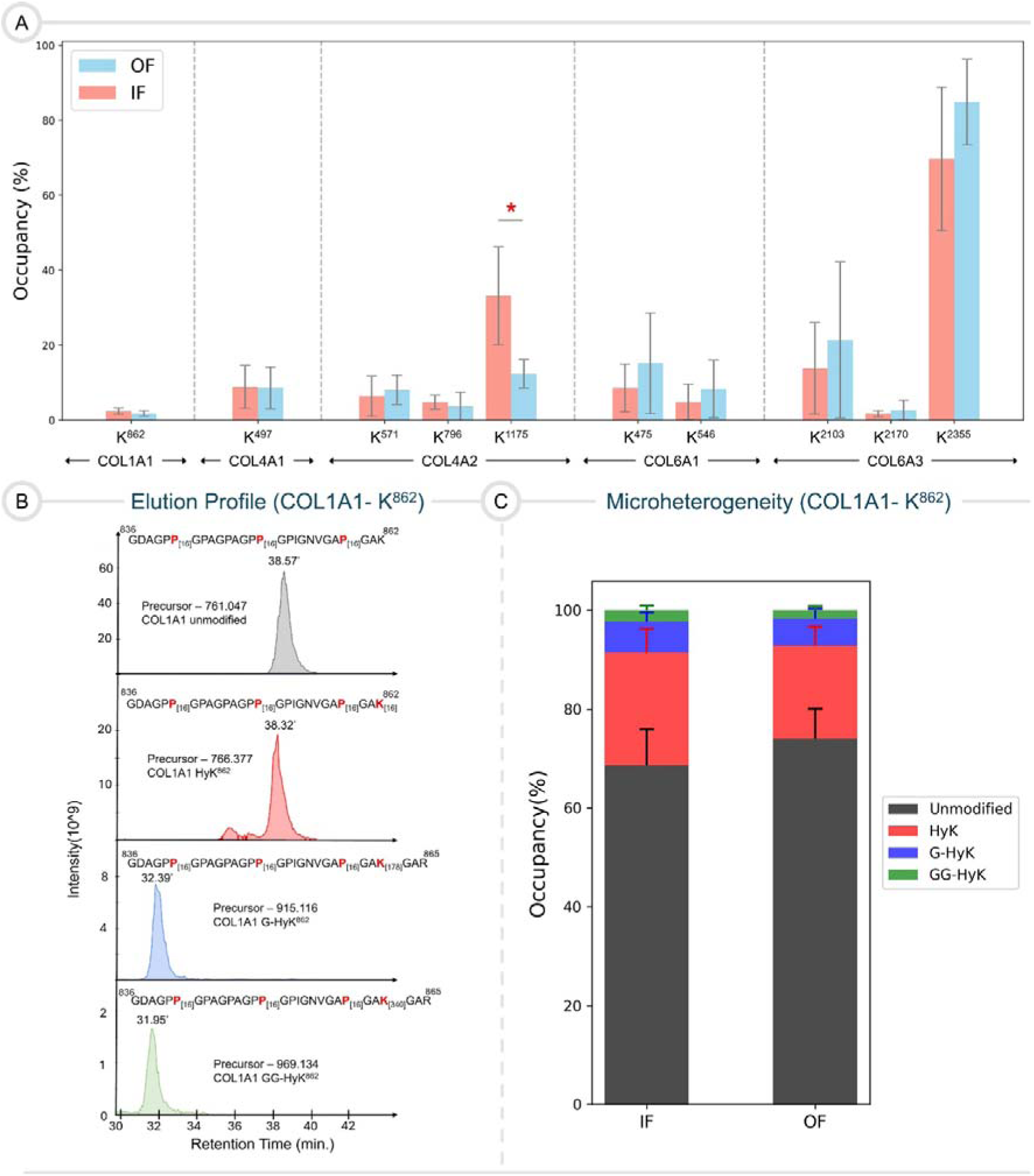
**A.** The bar plot represents the occupancy of GG-HyK sites in COL1A1, COL4A1, and COL4A2. **B.** The elution profile of the K^862^ site is shown. **C.** The bar plot shows the microheterogeneity in modifications of K^862^. The colors of grey, red, blue, and green represent the unmodified lysine, HyK, G-HyK, and GG-HyK, respectively. The difference in quantitation is considered statistically significant if p < 0.05.

**Table 2.** Occupancy values (%) of HyK in COL1A1 and COL4A2 (*p < 0.05; ^ns^p > 0.05)

| COL1A1 |  |  |
| --- | --- | --- |
| Sites | IF | OF |
| K<sup>277</sup> | 5.94 ± 2.11 | 12.45 ± 6.09<sup>ns</sup> |
| K<sup>352</sup> | 49.23 ± 11.96 | 37.41 ± 15.54<sup>ns</sup> |
| K<sup>397</sup> | 26.92 ± 16.82 | 33.86 ± 34.11<sup>ns</sup> |
| K<sup>586</sup> | 9.08 ± 6.88 | 7.97 ± 9.06<sup>ns</sup> |
| K<sup>862</sup> | 22.77 ± 4.83 | 18.85 ± 3.81<sup>ns</sup> |
| COL4A2 |  |  |
| K<sup>1346</sup> | 72.09 ± 5.28 | 65.87 ± 15.80<sup>ns</sup> |

**Table 3.** Occupancy values (%) table of GG-HyK in COL1A1, COL4A1, COL4A2, COL6A1 and COL6A3 (*p < 0.05; ^ns^p > 0.05)

| Collagen | Sites | IF | OF |
| --- | --- | --- | --- |
| **COL1A1** | K<sup>862</sup> | $2.34 \pm 0.83$ | $1.68 \pm 0.75^{\text{ns}}$ |
| **COL4A1** | K<sup>497</sup> | $8.80 \pm 5.68$ | $8.53 \pm 5.52^{\text{ns}}$ |
| **COL4A2** | K<sup>571</sup> | $6.32 \pm 5.38$ | $8.01 \pm 3.91^{\text{ns}}$ |
| | K<sup>796</sup> | $4.74 \pm 1.81$ | $3.62 \pm 3.78^{\text{ns}}$ |
| | K<sup>1175</sup> | $33.16 \pm 13.01$ | $12.36 \pm 3.79^*$ |
| **COL6A1** | K<sup>475</sup> | $8.49 \pm 6.31$ | $15.15 \pm 13.42^{\text{ns}}$ |
| | K<sup>546</sup> | $4.83 \pm 4.77$ | $8.28 \pm 7.68^{\text{ns}}$ |
| **COL6A3** | K<sup>2103</sup> | $13.79 \pm 12.28$ | $21.33 \pm 20.89^{\text{ns}}$ |
| | K<sup>2170</sup> | $1.65 \pm 0.77$ | $2.62 \pm 2.61^{\text{ns}}$ |
| | K<sup>2355</sup> | $69.68 \pm 19.10$ | $84.82 \pm 11.44^{\text{ns}}$ |

As lysine hydroxylation and glycosylation are prevalent in collagen IV, we quantified a few sites with these modifications. We could quantitate one site, K^1346^, of COL4A2 for hydroxylation. The K^1346^ is 65.87 ± 15.80% hydroxylated in OF, while 72.09 ± 5.28% in IF (Figure 7A, Table 2). To our knowledge, this site was not previously identified and quantitated in other human tissues. Further, we quantitated 1 (K^497^) and 3 (K^571^, K^796^, K^1175^) GG-HyK sites of COL4A1 and COL4A2, respectively (Figure 8A). The GG-HyK site (K^497^) have approximately similar occupancy level in OF (8.53 ± 5.52%) and IF (8.80 ± 5.68%) with a non-significant statistical difference (Figure 8A, Table 3). The quantitation of this site in human lung ^27^ and lens capsule ^26,43^ tissues was undefined (ColPTMScape). In COL4A2, among 3 sites, K^1175^ has the highest occupancy level and demonstrated the statistically significant difference across fractions (Figure 8A, Table 3). This site is highly glycosylated in IF as compared to OF (Figure 8A, Table 3). The K^571^ and K^796^ sites were not identified in human lungs, whereas the occupancy of K^1175^ sites was undefined ^27^. These observations indicate a high probability of lysine residues of collagen IV always being more glycosylated than collagen I, even in case of intra-tissue heterogeneity such as Adrenal gland (Figure 8A, Table 3) However, the hydroxylation of lysine residues can vary. It will be very interesting to look at the microheterogeneity of a lysine residue. Collagen IV has been experimentally shown to be more favorable than laminin and fibronectin for cortisol production from the human fetal adrenal glands following adrenocorticotropic hormone (ACTH) stimulation ^44–46^. It is highly likely that the alterations in the occupancy levels of PTMs, such as o-glycosylation of lysine residues, may have some functional role in steroid secretion. As observed from previous ^2^ and this work, COL6A1 and COL6A3 comprise top collagen abundant chains in the adrenal ECM. Collagen type VI interacts with several proteins ^47^, can play a role in cell differentiation ^2^ and cellular pluripotency ^2^, and can promote fibrosis ^47^. This intrigued us to quantitate GG-HyK sites of COL6A1 and COL6A3. Consequently, we quantitated 2 (K^475^, K^546^) and 3 (K^2103^, K^2170^, K^2355^) GG-HyK sites of COL6A1 and COL6A3, respectively. All sites from both chains displayed a statistically non-significant difference across fractions. In COL6A1 and COL6A3, sites are more glycosylated in OF compared to IF (Figure 8A, Table 3). The K^2355^ site in COL6A3 is highly glycosylated in OF as well as IF. This site was not identified in human lungs, and the other two sites of COL6A3 were not quantified. Similarly, in COL6A1, K^475^ was not identified in the human lungs, and the quantitation of K^546^ was undefined. The results highlights that these modified sites may contribute to the functions of OF.

### Microheterogeneity of Lysine Modifications in COL1A1 of OF and IF

The defects in the glycosylation of hydroxylated lysine residues can cause severe diseases ^48–50^. In the collagen chain, lysine residues can be available as unmodified or bear three modifications: hydroxylation (HyK), galactosyl-hydroxylation (G-HyK), or glucosylgalactosyl-hydroxylation (GG-HyK). Using our in-house pipeline, we observed the microheterogeneity on 1 lysine site (K^862^) in COL1A1 across both fractions. The quantitation of microheterogeneity demonstrated that there is no statistically significant difference across fractions. The initial hint of microheterogeneity can be followed by analyzing the chromatographic elution profile of such lysine sites. Through analyzing the elution profile of K^862^, we observed, as expected, the elution of peptides in the following order in minutes: GG-HyK containing peptide (31.95) > G-HyK containing peptide (32.39) > hydroxylated peptide (38.32) > unmodified peptide (38.57) (Figure 8B). The microheterogeneity observed for K^862^ highlights a high hydroxylation level in OF. In contrast, the level of o-glycosylation (G-HyK and GG-HyK) is high in IF (Figure 8C, Table 4). Similar to the previous site-specific quantitation, microheterogeneity in lysine modifications is different within a tissue.

**Table 4.** Microheterogeneity (%) in HyK^862^ residue in COL1A1 (*p < 0.05; ^ns^p > 0.05).

| Sites | IF | OF |
| --- | --- | --- |
| **Unmodified** | $68.5939 \pm 7.3064$ | $73.9489 \pm 6.0965^{\text{ns}}$ |
| **HyK** | $22.7712 \pm 4.8285$ | $18.8539 \pm 3.8126^{\text{ns}}$ |
| **G-HyK** | $6.2914 \pm 1.8539$ | $5.5175 \pm 1.9882^{\text{ns}}$ |
| **GG-HyK** | $2.3435 \pm 0.8287$ | $1.6797 \pm 0.7472^{\text{ns}}$ |

## Conclusion

In summary, we presented the analysis of collagen PTMs of the ECM of the human adrenal gland through this study. With the help of our in-house pipeline, we detected a more significant number of collagen chains in the adrenal gland than previously identified. We performed the MS^1^-based quantitation for collagen chains and defined the relative abundance of collagen chains using spectral numbers across both fractions. The relative abundance curve highlights higher levels of fibril and network-forming collagen chains in OF and IF, respectively, highlighting the importance of considering the PTM-modified collagen peptides. We comprehensively presented all the identified site-specific PTMs for the top 11 most abundant collagen chains across OF and IF. In this study, we generated the annotated COL1A1 PTM maps for OF and IF with a higher sequence coverage. We quantitated 3-HyP and lysine modification sites of COL1A1 to understand if there is an alteration in the occupancy level across OF and IF. For 3-HyP sites, we find no sites with the same occupancy in both fractions. 3 sites with higher occupancy in IF compared to OF are P^375^, P^570^, and P^870^. Interestingly, the osteogenesis imperfecta-linked site (P^1164^) has similar occupancy levels in both fractions (∼99% in IF and OF) than the bone (96-99%). The relative quantitation of HyK sites of COL1A1 has the same pattern as 3-HyP sites. As anticipated, comparing the hydroxylation levels of COL1A1 sites with COL4A2 shows the dominance of hydroxylation levels in COL4A2. For GG-HyK sites, we observed that the collagen IV sites were way more glycosylated than the only site (K^862^) available in COL1A1. The quantitation of thi modification sites in COL6A1 and COL6A3 demonstrated that the occupancy levels of some sites are higher than even COL4A1 and COL4A2, which raises a question regarding their role in the structural and biological functions of collagen type VI. The microheterogeneity was observed for this site, which clearly showed a higher level of hydroxylation in OF and glycosylation in IF. The change in level highlights that OF and IF may have different crosslinking patterns. To get an idea about the conservation of the inter-species’ heterogeneity of collagen within a tissue, we compared the 3-HyP and HyK sites of IF and OF of COL1A1 from human and rat adrenal cortex. We observed the conservation of ∼50% 3-HyP and ∼90% HyK sites of human COL1A1. Although the analysis primitively hints the essence of PTMs in collagen-ECM proteins interactions, the requirement of large-scale conservation of site-specific PTMs throughout the evolution needs to be explored. Collectively, we show the heterogeneity presented by collagen chains and PTMs within a healthy human adrenal (adult) gland, and it can differentially remodel the matrisome. The alteration in the heterogeneity during the pathophysiology of adrenal gland would provide detailed insights on the ECM remodelling. Further, our study not only generates a picture of collagenome within the human adrenal but also opens new avenues to understand the correlation of collagen PTMs’ role with the core function of the adrenal gland.

## Methods

### Mass Spectrometry Data Source

In the current study, we used previously published dataset, PXD051801, submitted to ProteomeXchange by Kremer *et al.* ^2^. The dataset contains the raw MS files of the ECM from the human adrenal cortex. They segmented the adrenal cortex into two fractions, the OF and IF (*vide supra*). The adrenal fractions’ ECM was enriched by decellularization using Triton-X and sodium deoxycholate ^2^. The MS analysis was performed on Orbitrap Exploris 480 (Thermo Fisher Scientific) connected to a nano-LC Vanquish Neo (Thermo Fisher Scientific) chromatograph. They performed data-dependent acquisition during MS analysis ^2^. We downloaded 10 .raw files, 5 for each fraction, OF and IF. Subsequently, we performed the MS database search.

### Database Search

To re-analyze the dataset, we performed the database search in two steps using MyriMatch ^51^ on the PARAM Himalaya Supercomputer at the Indian Institute of Technology Mandi. In the first step, a general search on the .RAW files was executed against the *Homo sapiens* database from Uniprot, which contains 20,220 entries. We set the precursor and fragment ion tolerances at 10 and 20 ppm, respectively. A static modification of carbamidomethylation (+57.0236) on cysteine and dynamic modifications of methionine oxidation (+15.9949) and prolyl hydroxylation (+15.9949) were incorporated in the search. We selected the option for fully tryptic protein digestion, allowing a maximum of two missed cleavages. The minimum and maximum peptide lengths were set to 5 and 40 residues, respectively. With this, the search generated .pepXML files. A parsimonious grouping of proteins identified in the search was performed after importing the .pepXML files in the IDPicker. We set the false discovery rate (FDR) to less than 1% for proteins, peptides, and peptide spectrum matches (PSMs). Consequently, a subset of the FASTA database was generated from IDPicker and used in the second PTM search.

For the PTM search, we used the same static modification as the previous search i,e, carbamidomethylating (+57.0236) on cysteine. However, the number of variable modifications was increased from two to five, and those were methionine oxidation (+15.9949), prolyl hydroxylation (+15.9949), lysyl hydroxylation (+15.9949), galactosyl-hydroxylysine (+178.0477), and glucosyl-galactosyl-hydroxylysine (+340.1006). For these modifications, we considered the tripeptide motifs of collagen as Gly-Xaa-Yaa. The prolyl hydroxylation at Yaa was considered 4-HyP, whereas that at Xaa was assigned as 3-HyP only if 4-HyP was available at Yaa in the motif. Similarly, the hydroxylation and glycosylations of lysine were assumed to be present in the Gly-Xaa-K motif. In addition to the modifications, the precursor and fragment ion tolerances were set to 10 and 20 ppm, respectively. In this search, we allowed for a maximum of three missed cleavages and a peptide length of 40 residues. Similar to the previous search, the parsimonious grouping of proteins was performed, and the false discovery rate (FDR) was kept at less than 1% for proteins, peptides, and PSMs. We manually inspected the PSMs and carefully assessed the MS/MS spectra before assigning them to the PTMs. The MS^1^-based relative quantitation for the collagen chains between OF and IF was performed using Skyline ^27^.

### Relative Quantitation of PTMs

The occupancy of collagen PTMs was quantitated for COL1A1, COL4A1, COL4A2, COL6A1, and COL6A3 using the previous pipeline ^27^. The PeptideProhet was utilized to generate the probability scores using the.pep.XML files. Subsequently, the spectral libraries were generated in Skyline (24.1, 64-bit) using .pep.XML and .RAW files using the previously described procedure ^27^. The MS/MS spectra were manually inspected for the unmodified and modified proline and lysine sites. For some of the PTM site’s quantitation, we have used Thermo XCalibur (2.2 SP1) in order to extract the ion-chromatogram and calculate the MS^1^peak area. The targeted extraction of the MS^1^ peak area for unmodified and modified proline and lysine sites was performed. As discussed, these peak areas were utilized to calculate the relative quantitation and microheterogeneity ^27^.

### Statistical Analysis

To confirm the significance of the change in PTM site occupancy, we performed a t-test using the SciPy library in Python and considered a p-value < 0.05 to be statistically significant. Subsequently, we plotted the relative abundance and occupancy plots graph using Matplotlib and the Seaborn library in Python.

## Supplementary Material Description

The supplementary file associated with this work contains tables related to collagen PTMs, figures related to conserved sites between rat and humans and COL1A2 PTM map, and PSMs associated with modifications of COL1A1.

## Author’s contribution

AJ and AN re-analyzed and generated the data, prepared the figures, and wrote the manuscript. CFPL and JLK generated the proteomic data in their lab. AJ, BM, and TB worked on summarizing and reviewing the manuscript. TB conceptualized the overall structure of the manuscript. All authors have made substantial intellectual contributions to this work.

## Funding

T.B. is grateful for the funding support from the Anusandhan National Research Foundation-Science and Engineering Research Board (ANRF-SERB) in the form of Core Research Grant CRG/2022/006204.

## Acknowledgment

AJ and AN are grateful for the fellowships from the Ministry of Education (MoE, Govt. of India). BM and TB acknowledge the CHCI centre of IIT-mandi. We acknowledge National Supercomputing Mission (NSM) for providing computing resources of ‘PARAM Himalaya’ at IIT Mand, which is implemented by C-DAC and supported by the Ministry of Electronics and Information Technology (MeitY) and Department of Science and Technology (DST), Government of India.

## Data Availability

The publicly available dataset PXD051801 was utilized in this study. The dataset can be accessed through the ProteomeXchange repository.

## Notes

### Competing Interest Statement

The authors have declared no competing interest.

