## Supporting Information for "Deciphering the intra-tissue specific Collagen PTMs site-specific heterogeneity in Human Adrenal extracellular-matrix"

Affiliations

Lab page- Proteomics Lab@IIT-Mandi

Table S1. Fold change comparison of ECM proteins of human adrenal gland

| Protein Gene | FC (Kremer et al.) | FC (This Work) |
| --- | --- | --- |
| COL1A1 | 1.03 | 1.46 |
| COL1A2 | 1.04 | 1.55 |
| COL6A1 | 1.03 | 1.28 |
| COL6A3 | 1.04 | 1.36 |
| FBN1 | 1.02 | 1.02 |
| HSPG2 | 1.02 | 1.06 |
| LAMB2 | 1.03 | 1.16 |
| LAMC1 | 1.02 | 0.9 |
| VCAN | 1.02 | 1.05 |
| POSTN | 1.06 | 1.36 |
| PKLR | 0.96 | 0.51 |

Table S2. PTMs identified across the top 11 collagen chains in the Outer Fraction (OF) of the Human Adrenal Gland

| Collagen chain | 3-HyP | HyK | G-HyK | GG-HyK | 4-HyP |
| --- | --- | --- | --- | --- | --- |
| COL1A1 | P³³³, P³⁷⁵, P⁴⁵⁹, P⁵⁵⁵, P⁵⁶⁷, P⁵⁷⁰, P⁷⁷¹, P⁸⁰⁷, P⁸¹⁶, P⁸⁴⁰, P⁸⁷⁰, P⁸⁸⁵, P⁸⁹⁴, P⁸⁹⁷, P⁹²⁴, P⁹²⁷, P⁹⁹⁶, P¹⁰⁴⁴, P¹¹¹⁹, P¹¹²², P¹¹⁶⁴ | K²⁷⁷, K²⁸⁶, K³⁵², K³⁹⁷, K⁴³⁰, K⁴⁴⁸, K⁵⁸⁶, K⁷⁸¹, K⁸²⁶, K⁸⁶², K⁹³⁴ | K²⁸⁶, K⁴⁴⁸, K⁸⁶² | K⁸⁶² | P²⁴¹, P²⁸⁹, P²⁹², P²⁹⁸, P³¹³, P³¹⁶, P³³⁴, P³⁴³, P³⁴⁶, P³⁷³, P³⁷⁶, P³⁸⁸, P³⁹⁴, P⁴⁰³, P⁴⁰⁹, P⁴¹², P⁴²⁷, P⁴³⁶, P⁴³⁹, P⁴⁵¹, P⁴⁶⁰, P⁴⁷⁵, P⁴⁸¹, P⁴⁸⁴, P⁵²³, P⁵²⁹, P⁵³⁵, P⁵⁴⁴, P⁵⁴⁷, P⁵⁵⁶, P⁵⁶⁵, P⁵⁶⁸, P⁵⁷¹, P⁵⁸³, P⁶⁰¹, P⁶⁰⁴, P⁶²², P⁶⁴⁰, P⁶⁴⁶, P⁶⁵², P⁶⁵⁸, P⁶⁶⁴, P⁶⁷⁰, P⁶⁹¹, P⁷¹⁵, P⁷¹⁸, P⁷²⁴, P⁷³⁰, P⁷⁷², P⁷⁷⁸, P⁷⁹⁹, P⁸⁰⁵, P⁸⁰⁸, P⁸¹⁷, P⁸²³, P⁸²⁹, P⁸⁴¹, P⁸⁵⁰, P⁸⁵⁹, P⁸⁷¹, P⁸⁷⁷, P⁸⁸⁶, P⁸⁹⁵, P⁸⁹⁸, P⁹¹⁹, P⁹²⁵, P⁹²⁸, P⁹³⁷, P⁹⁴⁶, P⁹⁴⁹, P⁹⁷³, P⁹⁷⁶, P⁹⁸², P⁹⁹⁷, P¹⁰⁰³, P¹⁰⁰⁹, P¹⁰¹⁸, P¹⁰²⁴, P¹⁰⁴⁵, P¹⁰⁴⁸, P¹⁰⁵¹, P¹⁰⁵⁴, P¹¹²⁰, P¹¹²³, P¹¹²⁶, P¹¹⁴⁴, P¹¹⁵⁰, P¹¹⁵⁹, P¹¹⁶⁵ |
| COL3A1 | P³⁸², P³⁸⁵, P⁴¹⁵, P⁵⁶⁵, P⁷¹², P⁷¹⁵, P⁷⁶⁹, P⁸⁰⁵, P⁸³⁸, P⁸⁴⁴, P⁸⁵³, P⁸⁸³, P⁸⁹², P⁹⁰⁴, P⁹⁴⁰, P⁹⁹⁴, P¹⁰⁵¹, P¹¹⁴⁷, P¹¹⁶² | K²⁷⁵, K²⁸⁴, K³⁵⁰, K³⁹⁵, K⁴⁶¹, K⁵⁰³, K⁷⁴⁰, K⁷⁴³, K⁷⁷⁹, K⁸²⁴, K⁸³³, K⁸⁶⁰, K⁹²³, K⁹³², K⁹⁷⁷ | K²⁷⁵, K⁷⁴³, K⁸⁶⁰ | K⁸⁶⁰ | P²⁸¹, P²⁹⁰, P²⁹⁶, P³³², P³³⁵, P³³⁸, P³⁴⁴, P³⁴⁷, P³⁵⁹, P³⁶⁵, P³⁷¹, P³⁸³, P³⁸⁶, P³⁹², P⁴⁰⁴, P⁴⁰⁷, P⁴¹⁶, P⁴²⁵, P⁴⁵⁵, P⁴⁵⁸, P⁴⁷⁰, P⁴⁷³, P⁴⁷⁹, P⁵⁰⁰, P⁵³⁹, P⁵⁴², P⁵⁴⁵, P⁵⁵¹, P⁵⁵⁴, P⁵⁶³, P⁵⁶⁶, P⁵⁷⁵, P⁵⁸¹, P⁵⁹⁹, P⁶⁰², P⁶⁰⁸, P⁶³⁵, P⁶⁴⁴, P⁶⁵⁰, P⁶⁸⁶, P⁶⁹², P⁷⁰¹, P⁷¹³, P⁷¹⁶, P⁷²², P⁷²⁸, P⁷³⁷, P⁷⁴⁶, P⁷⁴⁹, P⁷⁵⁵, P⁷⁷⁰, P⁷⁷⁶, P⁷⁸⁵, P⁷⁸⁸, P⁷⁹⁷, P⁸⁰⁶, P⁸¹², P⁸¹⁵, P⁸²¹, P⁸³⁰, P⁸³⁹, P⁸⁴⁵, P⁸⁵⁴, P⁸⁶⁶, P⁸⁶⁹, P⁸⁷⁵, P⁸⁸¹, P⁸⁸⁴, P⁸⁹⁰, P⁸⁹³, P⁸⁹⁹, P⁹⁰⁵, P⁹¹⁴, P⁹¹⁷, P⁹²⁹, P⁹³⁵, P⁹⁴¹, P⁹⁴⁴, P⁹⁷¹, P⁹⁸³, P⁹⁹⁵, P¹⁰⁰¹, P¹⁰¹⁰, P¹⁰¹⁶, P¹⁰²², P¹⁰⁴⁰, P¹⁰⁴³, P¹⁰⁴⁶, P¹⁰⁴⁹, P¹⁰⁵², P¹⁰⁷⁶, P¹¹¹², P¹¹¹⁵, P¹¹¹⁸, P¹¹²¹, P¹¹³³, P¹¹⁴⁸, P¹¹⁵⁷, P¹¹⁶³ |
| COL1A2 | P¹⁰¹, P²⁵⁴, P²⁹⁶, P³⁷¹, P³⁹², P⁵³³, P⁵⁴⁸, P⁵⁵¹, P⁶¹⁴, P⁷⁷⁰, P⁷⁹⁷, P⁸⁰⁶, P⁸⁰⁹, P⁸⁵⁷, P⁹⁰² | K¹⁸⁹, K¹⁹⁸, K²⁶⁴, K³⁰⁹, K³⁵⁴, K³⁶⁰, K⁴⁹⁸, K⁵¹⁰, K⁶²¹, K⁶⁵⁴, K⁶⁵⁷, K⁷³⁸, K⁷⁴⁷, K¹⁰⁰⁸, K¹⁰¹⁴ | K⁷³⁸ | None | P¹⁰², P¹⁰⁸, P¹²⁰, P¹²³, P¹⁴⁷, P¹⁵⁰, P¹⁵³, P¹⁹², P¹⁹⁵, P²⁰¹, P²⁰⁴, P²¹⁰, P²⁵⁵, P²⁵⁸, P²⁶¹, P²⁸⁸, P²⁹⁷, P³⁰⁰, P³¹⁵, P³²¹, P³²⁴, P³³⁰, P³⁴⁸, P³⁶³, P³⁷², P³⁹³, P³⁹⁶, P⁴⁴¹, P⁴⁴⁴, P⁴⁵³, P⁴⁵⁶, P⁴⁷¹, P⁴⁷⁷, P⁵³⁴, P⁵⁴⁹, P⁵⁵², P⁵⁵⁸, P⁵⁷⁰, P⁵⁸², P⁵⁹⁴, P⁶¹⁵, P⁶²⁴, P⁶⁴², P⁶⁵¹, P⁶⁶⁰, P⁶⁶⁹, P⁶⁷⁸, P⁶⁸⁴, P⁷³⁵, P⁷⁷¹, P⁷⁸³, P⁷⁸⁹, P⁷⁹⁸, P⁸⁰⁷, P⁸¹⁰, P⁸⁴⁰, P⁸⁵⁸, P⁸⁶¹, P⁸⁷⁰, P⁸⁷⁶, P⁸⁸⁵, P⁸⁹⁴, P⁹⁰³, P⁹⁰⁹, P⁹¹⁵, P⁹²¹, P⁹³⁰, P⁹³⁶, P⁹⁵¹, P⁹⁶³, P¹⁰¹¹, P¹⁰³², P¹⁰⁴⁴, P¹⁰⁷¹, P¹⁰⁹², P¹⁰⁹⁵, P¹⁰⁹⁸, P¹¹⁰¹ |
| COL6A3 | P^2238^ | None | K²³⁵⁵ | K²⁰⁵², K²¹⁰³, K²¹⁷⁰, K²³⁵⁵ | P²¹⁰⁰, P²¹⁵⁷, P²²³⁹, P²³⁵⁸, P²³⁶¹ |
| COL5A1 | P⁵⁰⁰, P⁵¹⁵, P⁵⁷⁵, P⁶⁵³, P⁶⁵⁶, P⁷⁷⁹, P⁸³³, P⁹⁴⁴, P⁹⁹², P⁹⁹⁵, P¹⁰⁷⁹, P¹⁰⁸² | K⁵⁸², K⁹⁶³ | None | K⁵⁸², K⁷⁷⁴ | P⁵⁰¹, P⁵⁰⁴, P⁵⁰⁷, P⁵¹³, P⁵¹⁶, P⁵⁷⁰, P⁵⁷⁶, P⁵⁸⁵, P⁶⁰⁰, P⁶⁰⁶, P⁶³⁹, P⁶⁴⁸, P⁶⁵⁴, P⁶⁵⁷, P⁷⁵⁰, P⁷⁵³, P⁷⁵⁶, P⁷⁶², P⁷⁶⁵, P⁷⁷¹, P⁷⁸⁰, P⁷⁸⁹, P⁸⁷⁰, P⁸⁷³, P⁸⁷⁶, P⁹⁶⁰, P⁹⁹³, P⁹⁹⁶, P¹⁰⁴¹, P¹⁰⁴⁷, P¹¹⁰⁴, P¹³⁹⁵, P¹⁴³¹, P¹⁴³⁴ |
| COL5A2 | P¹³⁶, P⁴⁰⁹, P⁴⁴², P⁹¹⁹, P¹⁰³⁰, P¹¹⁵⁶, P¹¹⁹⁸ | K⁴³¹, K⁸⁶⁰ | None | K¹¹³⁰ | P¹³¹, P¹³⁷, P²⁶⁰, P²⁶³, P²⁶⁹, P²⁷⁵, P²⁸⁴, P⁴¹⁰, P⁴¹⁶, P⁴¹⁹, P⁴²⁸, P⁴³⁷, P⁴⁴³, P⁴⁴⁹, P⁴⁵², P⁴⁸⁵, P⁴⁹⁷, P⁵⁰⁹, P⁵⁹³, P⁵⁹⁹, P⁶⁰², P⁸⁵⁷, P⁹²⁰, P⁹²⁶, P⁹³⁵, P⁹⁴¹, P¹⁰³¹, P¹⁰³⁷, P¹⁰⁴⁶, P¹⁰⁵⁸, P¹⁰⁷³, P¹⁰⁷⁹, P¹⁰⁸⁵, P¹⁰⁸⁸, P¹⁰⁹⁴, P¹¹⁵⁴, P¹¹⁵⁷, P¹¹⁶⁹, P¹¹⁷⁸, P¹¹⁹⁹ |
| COL4A2 | P⁷⁴, P¹⁹⁷, P¹⁴¹² | K¹³⁴⁶ |  | K⁵⁷¹, K⁷⁹⁶, K¹¹⁷⁵, K¹³⁷³ | P⁶³, P⁷⁵, P⁸¹, P¹⁸⁶, P¹⁸⁹, P¹⁹⁸, P²⁰¹, P²¹⁰, P²¹³, P⁴⁶², P⁴⁶⁵, P⁴⁶⁸, P⁷⁰⁷, P⁷¹⁶, P⁷³¹, P⁷³⁴, P⁷⁷³, P⁷⁹⁰, P⁹⁷³, P⁹⁷⁶, P¹¹⁷², P¹¹⁸¹, P¹¹⁸⁴, P¹¹⁸⁷, P¹¹⁹⁰, P¹²⁰⁸, P¹²¹¹, P¹²¹⁹, P¹²²², P¹²²⁵, P¹³³⁷, P¹³⁴³, P¹³⁷⁹, P¹³⁸², P¹³⁸⁸, P¹⁴¹³, P¹⁴¹⁶, P¹⁴²⁵, P¹⁴²⁸, P¹⁴⁷⁰, P¹⁴⁷³, P¹⁴⁷⁶, P¹⁴⁷⁹, P¹⁴⁸² |
| COL4A1 | P⁴⁷⁸, P⁴⁸⁴, P⁵⁸⁷, P⁶⁴⁷, P⁷⁸⁶, P⁸⁰³ | K²⁹⁸, K⁸²⁸ | None | K⁴⁹⁷ | P²¹⁰, P³⁰⁴, P³⁰⁷, P³¹⁰, P³⁷⁶, P³⁸⁴, P³⁸⁷, P³⁹⁹, P⁴⁷⁹, P⁴⁸⁵, P⁴⁹¹, P⁴⁹⁴, P⁵⁸⁸, P⁵⁹⁴, P⁶⁴⁸, P⁶⁵⁴, P⁶⁵⁷, P⁶⁶⁰, P⁷⁸⁴, P⁷⁸⁷, P⁷⁹⁰, P⁷⁹⁶, P⁷⁹⁹, P⁸⁰⁴ |
| COL12A1 | P²⁷⁸⁴, P²⁷⁹⁰ | None | None | None | P²⁷⁸⁵, P²⁷⁹¹, P²⁹⁵⁹, P²⁹⁶⁵, P²⁹⁶⁸, P²⁹⁷¹, P²⁹⁷⁷ |
| COL16A1 | P¹⁰⁴⁶, P¹⁰⁵², P¹¹²¹, P¹²⁰², P¹²⁰⁵, P¹³³³, P¹³³⁹, P¹⁴⁰², P¹⁵³⁷ | K¹¹¹⁶, K¹²⁴⁵ | None | None | P⁷⁹⁶, P⁷⁹⁹, P⁸⁰², P⁸⁰⁸, P¹⁰⁴⁴, P¹⁰⁴⁷, P¹⁰⁵³, P¹⁰⁵⁶, P¹⁰⁶², P¹⁰⁶⁵, P¹¹¹⁹, P¹¹²², P¹¹²⁸, P¹¹³¹, P¹¹⁵⁸, P¹¹⁶¹, P¹¹⁶⁴, P¹¹⁶⁷, P¹¹⁷⁰, P¹¹⁹⁷, P¹²⁰⁰, P¹²⁰³, P¹²⁰⁶, P¹²³⁰, P¹²³⁶, P¹²⁴², P¹²⁶³, P¹²⁶⁶, P¹²⁷², P¹²⁷⁸, P¹³²⁸, P¹³³¹, P¹³³⁴, P¹³³⁷, P¹³⁴⁰, P¹³⁴³, P¹⁴⁰³, P¹⁴⁸⁴, P¹⁴⁸⁷, P¹⁴⁹⁰, P¹⁴⁹³, P¹⁴⁹⁶, P¹⁵³⁴, P¹⁵³⁷, P¹⁵⁴³ |
| COL6A1 | P⁵⁸⁸ | K⁵⁴⁶ | K⁵⁴⁶ | K⁴⁷⁵, K⁵⁴⁶ | P⁴⁴⁵, P⁴⁶³, P⁴⁶⁶, P⁴⁹⁹, P⁵⁰², P⁵⁴³, P⁵⁵⁵, P⁵⁸³, P⁵⁸⁹ |

Table S3. PTMs identified across the top 11 collagen chains in the Inner Fraction (IF) of the Human Adrenal Gland

| Collagen chain | 3-HyP | HyK | G-HyK | GG-HyK | 4-HyP |
| --- | --- | --- | --- | --- | --- |
| COL1A1 | P³³³, P³⁷⁵, P⁴²⁶, P⁴⁵⁹, P⁵⁵⁵, P⁵⁶⁷, P⁵⁷⁰, P⁶⁰³, P⁶⁹⁰, P⁷⁷¹, P⁸⁰⁷, P⁸¹⁶, P⁸⁴⁰, P⁸⁷⁰, P⁸⁸⁵, P⁸⁹⁴, P⁸⁹⁷, P⁹²⁷, P⁹⁹⁶, P¹⁰⁴⁴, P¹¹¹⁹, P¹¹²², P¹¹⁶⁴ | K²⁷⁷, K²⁸⁶, K³⁵², K³⁹⁷, K⁴⁴⁸, K⁵³⁸, K⁵⁸⁶, K⁷⁸¹, K⁸²⁶, K⁸⁶², K⁹³⁴ | K⁴⁴⁸, K⁸⁶² | K⁸⁶² | P²⁴¹, P²⁸⁹, P²⁹², P²⁹⁸, P³¹³, P³¹⁶, P³³⁴, P³⁴³, P³⁴⁶, P³⁷³, P³⁷⁶, P³⁸⁸, P³⁹⁴, P⁴⁰³, P⁴⁰⁹, P⁴¹², P⁴²⁷, P⁴⁵¹, P⁴⁶⁰, P⁴⁷⁵, P⁴⁸¹, P⁴⁸⁴, P⁴⁹⁶, P⁵²³, P⁵²⁹, P⁵³⁵, P⁵⁴⁴, P⁵⁴⁷, P⁵⁵⁶, P⁵⁶⁵, P⁵⁶⁸, P⁵⁷¹, P⁵⁸³, P⁶⁰¹, P⁶⁰⁴, P⁶⁴⁰, P⁶⁴⁶, P⁶⁵², P⁶⁵⁸, P⁶⁶⁴, P⁶⁷⁰, P⁶⁹¹, P⁷¹⁵, P⁷¹⁸, P⁷²⁴, P⁷³⁰, P⁷⁷², P⁷⁷⁸, P⁷⁹⁹, P⁸⁰⁵, P⁸⁰⁸, P⁸¹⁷, P⁸²³, P⁸²⁹, P⁸⁴¹, P⁸⁵⁰, P⁸⁵⁹, P⁸⁷¹, P⁸⁷⁷, P⁸⁸⁶, P⁸⁹⁵, P⁸⁹⁸, P⁹¹⁹, P⁹²⁵, P⁹²⁸, P⁹³⁷, P⁹⁴⁶, P⁹⁴⁹, P⁹⁶⁴, P⁹⁷³, P⁹⁷⁶, P⁹⁸², P⁹⁹⁷, P¹⁰⁰³, P¹⁰⁰⁹, P¹⁰¹⁸, P¹⁰²⁴, P¹⁰⁴⁵, P¹⁰⁴⁸, P¹⁰⁵¹, P¹⁰⁵⁴, P¹¹²⁰, P¹¹²³, P¹¹²⁶, P¹¹⁴⁴, P¹¹⁵⁰, P¹¹⁵⁹, P¹¹⁶⁵ |
| COL3A1 | P³³⁴, P³⁸², P³⁸⁵, P⁴¹⁵, P⁵⁵³, P⁵⁶⁵, P⁶⁰⁷, P⁶⁸⁵, P⁷⁰⁰, P⁷¹², P⁷¹⁵, P⁷⁶⁹, P⁸⁰⁵, P⁸³⁸, P⁸⁴⁴, P⁸⁵³, P⁸⁸³, P⁸⁹², P⁹⁰⁴, P⁹⁴⁰, P⁹⁶¹, P⁹⁹⁴, P¹⁰⁵¹, P¹¹⁴⁷, P¹¹⁶² | K²⁶³, K²⁷⁵, K²⁸⁴, K³⁵⁰, K³⁹⁵, K⁴⁶¹, K⁵⁰³, K⁵⁸⁴, K⁶⁶², K⁷⁴⁰, K⁷⁴³, K⁷⁷⁹, K⁸²⁴, K⁸³³, K⁸⁶⁰, K⁹²³, K⁹³², K⁹⁷⁷ | K⁷⁴⁰, K⁸⁶⁰, K⁹²³ | K⁸⁶⁰ | P²⁵⁷, P²⁶⁰, P²⁸¹, P²⁹⁰, P²⁹⁶, P³¹¹, P³¹⁴, P³³², P³³⁵, P³³⁸, P³⁴⁴, P³⁴⁷, P³⁵⁹, P³⁶⁵, P³⁷¹, P³⁸³, P³⁸⁶, P³⁹², P⁴⁰⁴, P⁴⁰⁷, P⁴¹⁶, P⁴²⁵, P⁴⁵⁵, P⁴⁵⁸, P⁴⁷⁰, P⁴⁷³, P⁴⁷⁹, P⁵⁰⁰, P⁵³⁹, P⁵⁴², P⁵⁴⁵, P⁵⁵¹, P⁵⁵⁴, P⁵⁶³, P⁵⁶⁶, P⁵⁷⁵, P⁵⁸¹, P⁵⁹⁹, P⁶⁰², P⁶⁰⁸, P⁶³⁵, P⁶⁴⁴, P⁶⁵⁰, P⁶⁶⁸, P⁶⁷¹, P⁶⁸⁶, P⁶⁹², P⁷⁰¹, P⁷¹³, P⁷¹⁶, P⁷²², P⁷²⁸, P⁷³⁷, P⁷⁴⁶, P⁷⁴⁹, P⁷⁵⁵, P⁷⁷⁰, P⁷⁷⁶, P⁷⁸⁵, P⁷⁸⁸, P⁸⁰⁶, P⁸¹², P⁸¹⁵, P⁸²¹, P⁸³⁰, P⁸³⁹, P⁸⁴⁵, P⁸⁵⁴, P⁸⁶⁶, P⁸⁶⁹, P⁸⁷⁵, P⁸⁸¹, P⁸⁸⁴, P⁸⁹⁰, P⁸⁹³, P⁸⁹⁹, P⁹⁰⁵, P⁹¹⁴, P⁹¹⁷, P⁹²⁹, P⁹³⁵, P⁹⁴¹, P⁹⁴⁴, P⁹⁶², P⁹⁶⁵, P⁹⁷¹, P⁹⁸³, P⁹⁹⁵, P¹⁰⁰¹, P¹⁰¹⁰, P¹⁰¹⁶, P¹⁰²², P¹⁰⁴⁰, P¹⁰⁴³, P¹⁰⁴⁶, P¹⁰⁴⁹, P¹⁰⁵², P¹⁰⁷⁶, P¹⁰⁸⁵, P¹¹¹², P¹¹¹⁵, P¹¹¹⁸, P¹¹²¹, P¹¹³³, P¹¹⁴⁸, P¹¹⁵⁷, P¹¹⁶³ |
| COL1A2 | P¹⁰¹, P²⁵⁴, P²⁹⁶, P³⁷¹, P³⁹², P³⁹⁵, P⁴¹⁹, P⁵³³, P⁵⁴⁸, P⁵⁵¹, P⁶¹⁴, P⁷⁷⁰, P⁷⁹⁷, P⁸⁰⁶, P⁸⁰⁹, P⁸⁵⁷, P⁹⁰² | K¹⁸⁹, K¹⁹⁸, K²⁶⁴, K³⁰⁹, K³⁵⁴, K³⁶⁰, K⁵⁴³, K⁶²¹, K⁶⁵⁴, K⁶⁵⁷, K⁷³⁸, K⁷⁴⁷, K¹⁰⁰⁸, K¹⁰¹⁴ | K¹⁷⁷, K⁷³⁸, K⁷⁴⁷ | None | P¹⁰², P¹⁰⁸, P¹²⁰, P¹²³, P¹⁴⁷, P¹⁵⁰, P¹⁵³, P¹⁶⁸, P¹⁷¹, P¹⁷⁴, P¹⁹², P¹⁹⁵, P²⁰¹, P²⁰⁴, P²¹⁰, P²²⁸, P²⁵⁵, P²⁵⁸, P²⁶¹, P²⁸⁸, P²⁹⁷, P³⁰⁰, P³¹⁵, P³²¹, P³²⁴, P³³⁰, P³⁴⁸, P³⁶³, P³⁷²,P³⁹³, P³⁹⁶, P⁴²⁰, P⁴⁴¹, P⁴⁴⁴, P⁴⁵³, P⁴⁵⁶, P⁴⁷¹, P⁴⁷⁷, P⁴⁸⁹, P⁴⁹⁵, P⁵²², P⁵³⁴, P⁵⁴⁹, P⁵⁵², P⁵⁵⁸, P⁵⁷⁰, P⁵⁸², P⁵⁹⁴, P⁶¹⁵, P⁶²⁴, P⁶⁴², P⁶⁷⁸, P⁶⁸⁴, P⁷³⁵, P⁷⁷¹, P⁷⁸³, P⁷⁸⁹, P⁷⁹⁸, P⁸⁰⁷, P⁸¹⁰, P⁸⁴⁰, P⁸⁵⁸, P⁸⁶¹, P⁸⁷⁰, P⁸⁷⁶, P⁸⁸⁵, P⁸⁹⁴, P⁹⁰³, P⁹⁰⁹, P⁹¹⁵, P⁹²¹, P⁹³⁰, P⁹³⁶, P⁹⁵¹, P⁹⁶³, P¹⁰¹¹, P¹⁰³², P¹⁰⁴⁴, P¹⁰⁷¹ |
| COL6A3 | P²²³⁸ | None | None | K²¹⁰³, K²¹⁷⁰, K²³⁵⁵ | P²⁰⁶⁴, P²¹⁰⁰, P²¹⁵⁷, P²¹⁸², P²²³⁹, P²³⁵⁸, P²³⁶¹ |
| COL5A1 | P⁵⁰¹, P⁵⁷⁵, P⁵⁹⁹, P⁶⁵³, P⁶⁵⁶, P⁷⁷⁹, P⁸³³, P⁹⁴⁴, P⁹⁹², P⁹⁹⁵, P¹³²², P¹³⁹⁴ | K⁵⁸², K⁷⁹⁵, K⁸⁶⁴, K⁹⁶³ | None | K⁵⁸², K⁷⁷⁴, K¹⁰³⁸, K¹¹²⁵ | P⁵⁰¹, P⁵⁰⁴, P⁵⁰⁷, P⁵¹³, P⁵¹⁶, P⁵⁷⁰, P⁵⁷⁶, P⁵⁸⁵, P⁶⁰⁰, P⁶⁰⁶, P⁶⁴⁸, P⁶⁵⁴, P⁶⁵⁷, P⁷⁵⁰, P⁷⁵³, P⁷⁵⁶, P⁷⁶², P⁷⁶⁵, P⁷⁷¹, P⁷⁸⁰, P⁷⁸⁹, P⁸⁵⁵, P⁸⁶¹, P⁸⁷⁰, P⁸⁷³, P⁸⁷⁶, P⁹⁶⁰, P⁹⁹³, P⁹⁹⁶, P¹¹⁰⁴, P¹³⁹⁵, P¹⁴³¹, P¹⁴³⁴ |
| COL5A2 | P¹³⁶, P⁴⁰⁹, P⁴⁴², P⁵¹⁴, P⁹¹⁹, P¹⁰³⁰, P¹⁰³⁶, P¹¹⁵⁶, P¹¹⁹⁸ | K⁴³¹, K⁸⁶⁰ | None | None | P¹³¹, P¹³⁷, P¹⁶⁴, P¹⁶⁷, P¹⁷⁰, P¹⁷³, P¹⁷⁶, P¹⁷⁹, P²⁶⁰, P²⁶³, P²⁶⁹, P²⁷⁵, P²⁸⁴, P³⁵³, P⁴¹⁰, P⁴¹⁶, P⁴¹⁹, P⁴²⁸, P⁴³⁷, P⁴⁴³, P⁴⁴⁹, P⁴⁵², P⁴⁸⁵, P⁴⁹⁷, P⁵⁰⁹, P⁵¹⁵, P⁵⁷⁸, P⁵⁹³, P⁵⁹⁹, P⁶⁰², P⁸⁵⁷, P⁹²⁰, P⁹²⁶, P⁹³⁵, P⁹⁶², P⁹⁶⁵, P⁹⁷¹, P⁹⁷⁷, P⁹⁸³, P¹⁰⁰⁷, P¹⁰¹, P¹⁰¹⁶, P¹⁰³¹, P¹⁰³⁷, P¹⁰⁴⁶, P¹⁰⁵⁸, P¹⁰⁷³, P¹⁰⁷⁹, P¹⁰⁸⁵, P¹⁰⁸⁸, P¹⁰⁹⁴, P¹¹⁵⁴, P¹¹⁵⁷, P¹¹⁶⁹ |
| COL4A2 | P⁷⁴, P¹⁹⁷, P⁹⁷⁵, P¹⁴¹² | K¹¹⁷⁵, K¹³⁴⁶, K¹³⁷³ | None | K⁵⁷¹, K⁷⁹⁶, K⁸⁸³, K⁹⁸⁷, K¹¹⁷⁵, K¹³⁷³ | P⁶³, P⁷⁵, P⁸¹, P¹¹⁴, P¹²⁰, P¹²³, P¹⁸⁶, P¹⁸⁹, P¹⁹⁸, P²⁰¹, P²¹³, P⁴⁶², P⁴⁶⁵, P⁴⁶⁸, P⁷⁰⁷, P⁷¹⁶, P⁷³¹, P⁷³⁴, P⁷⁷³, P⁷⁸¹, P⁷⁹⁰, P⁷⁹³, P⁸⁷¹, P⁸⁷⁷, P⁹⁷³, P⁹⁷⁶, P¹¹⁷², P¹¹⁸¹, P¹¹⁸⁴, P¹¹⁸⁷, P¹¹⁹⁰, P¹²⁰⁸, P¹²¹¹, P¹²¹⁹, P¹²²², P¹²²⁵, P¹³³⁷, P¹³⁴³, P¹³⁷⁹, P¹³⁸², P¹³⁸⁸, P¹⁴¹³, P¹⁴¹⁶, P¹⁴²⁵, P¹⁴²⁸, P¹⁴⁷³, P¹⁴⁷⁶, P¹⁴⁷⁹, P¹⁴⁸², P¹⁶⁸⁸ |
| COL4A1 | P⁴⁷⁸, P⁵⁸⁷, P⁶⁴⁷, P⁷⁸⁶, P⁸⁰³ | None | None | K²⁹⁸, K⁴⁹⁷ | P³⁰¹, P³⁰⁴, P³⁰⁷, P³¹⁰, P³⁹⁹, P⁴⁷⁹, P⁴⁸⁵, P⁴⁹¹, P⁴⁹⁴, P⁵⁸⁸, P⁵⁹⁴, P⁶⁴⁸, P⁶⁵⁴, P⁶⁵⁷, P⁶⁶⁰, P⁷⁸⁴, P⁷⁸⁷, P⁷⁹⁰, P⁷⁹⁶, P⁷⁹⁹, P⁸⁰⁴ |
| COL12A1 | P²⁷⁸⁴ | None | None | None | P²⁷⁸⁵, P²⁷⁹¹, P²⁹⁵⁹, P²⁹⁶⁵, P²⁹⁶⁸, P²⁹⁷¹, P²⁹⁷⁷ |
| COL16A1 | P¹⁰⁴⁶, P¹⁰⁵², P¹¹²¹, P¹²⁰², P¹²⁰⁵, P¹³³⁹, P¹⁵³⁶ | K¹¹¹⁶ | None | None | P¹⁰⁴⁴, P¹⁰⁴⁷, P¹⁰⁵³, P¹⁰⁵⁶, P¹⁰⁶², P¹⁰⁶⁵, P¹¹¹⁹, P¹¹²², P¹¹²⁸, P¹¹³¹, P¹¹⁷⁶, P¹¹⁷⁹, P¹¹⁹⁷, P¹²⁰⁰, P¹²⁰³, P¹²⁰⁶, P¹²⁶³, P¹²⁶⁶, P¹²⁷², P¹²⁷⁸, P¹³²⁸, P¹³³¹, P¹³³⁴, P¹³³⁷, P¹³⁴⁰, P¹³⁴³, P¹⁴⁸⁴, P¹⁴⁸⁷, P¹⁴⁹⁰, P¹⁴⁹³, P¹⁴⁹⁶, P¹⁵³⁴, P¹⁵³⁷, P¹⁵⁴³ |
| COL6A1 | P⁵⁸⁸ | K⁵⁴⁶ | None | K⁴⁷⁵, K⁵⁴⁶ | P⁴⁴⁵, P⁴⁶³, P⁴⁶⁶, P⁴⁹³, P⁴⁹⁹, P⁵⁰², P⁵⁴³, P⁵⁵⁵, P⁵⁸³, P⁵⁸ |

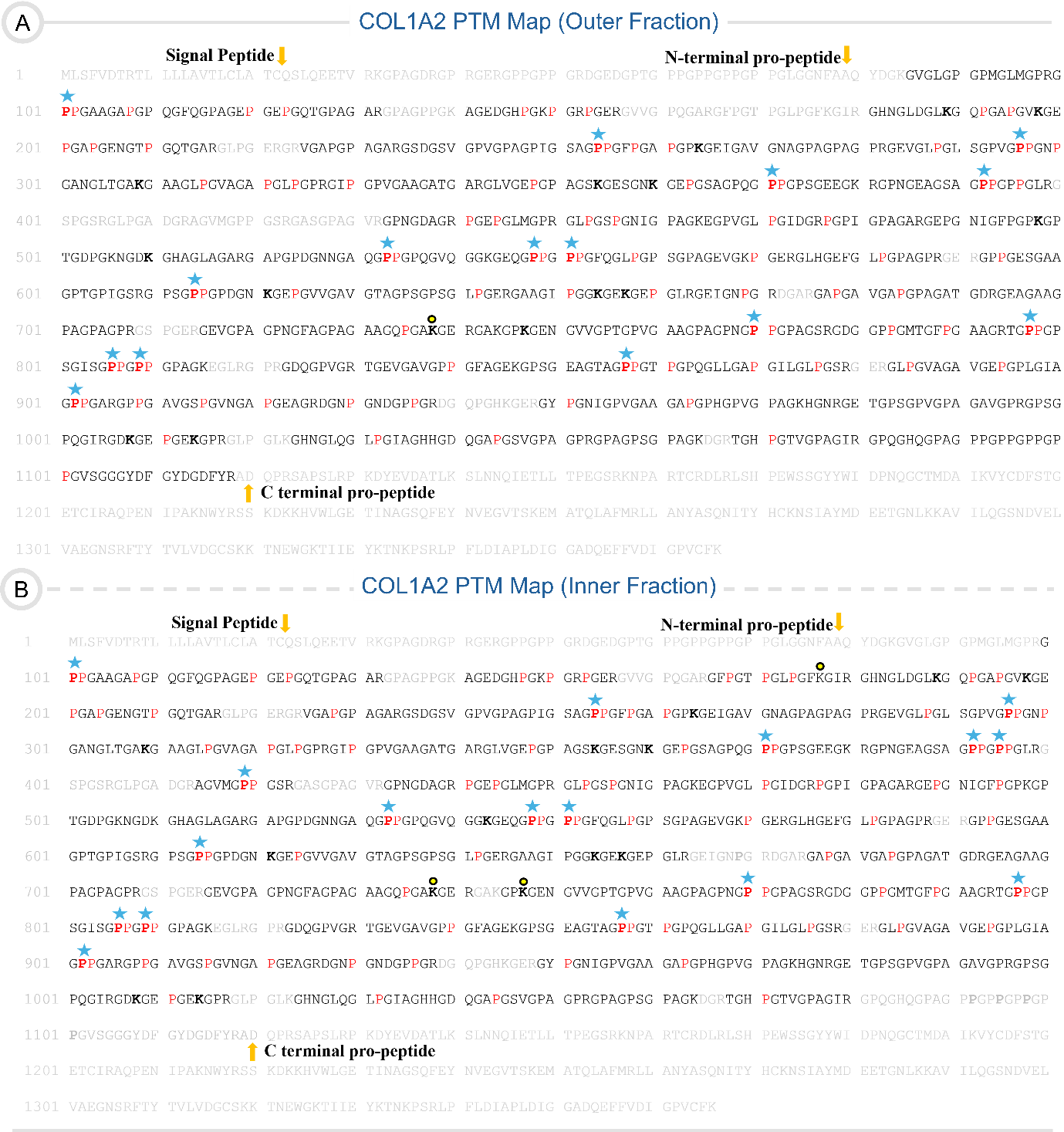

Figure S1. COL1A2 PTM map of the OF (A) and IF (B). The sequence coverages of OF and IF COL1A2 maps are 88.28% and 84.63%, respectively. The cleavage points for the signal peptide, N-terminal, and C-terminal propeptides are marked with the yellow arrow. The red P represents the 4-HyP, and the bold red P with a blue star highlights the presence of 3-HyP. The HyK, G-HyK, and GG-HyK are represented by bold K, bold K with yellow circles, and bold K with yellow and yellow-blue combined circles, respectively.

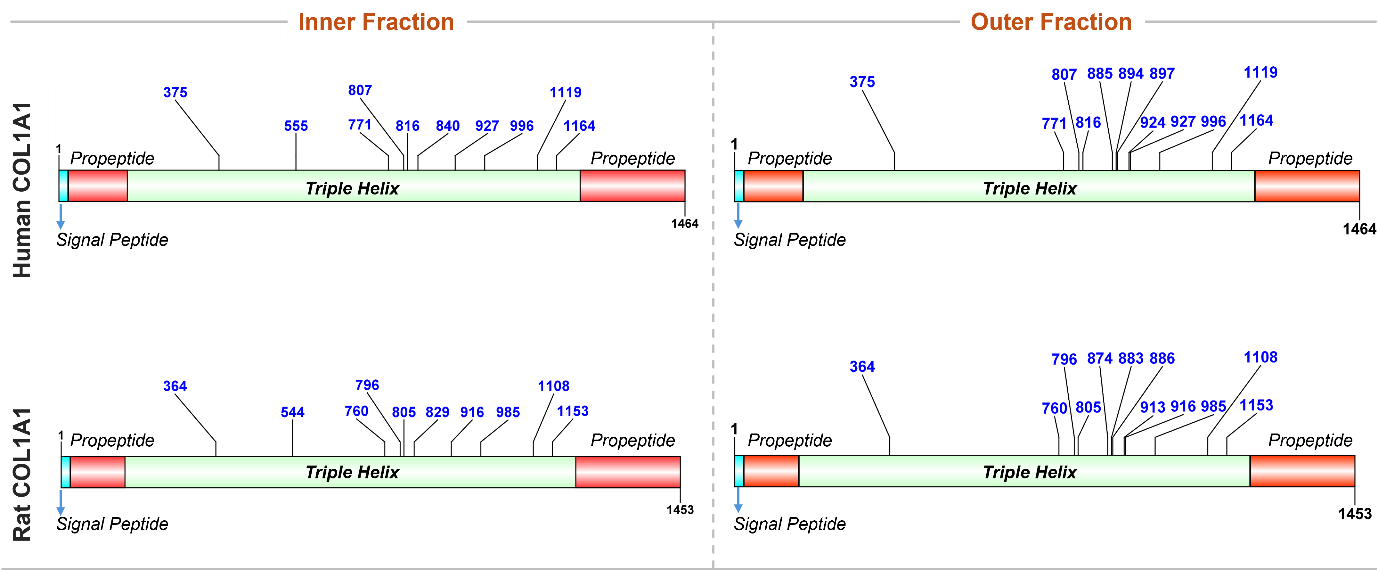

Figure S2. Comparative analysis of conserved 3-HyP sites between human and rat COL1A1 chains. The rectangular box represents the COL1A1 chain that has signal peptide (light blue) propeptide (orange), and triple helix (light green) domains. All conserved sites (site numbers in blue) are depicted with the black lines.

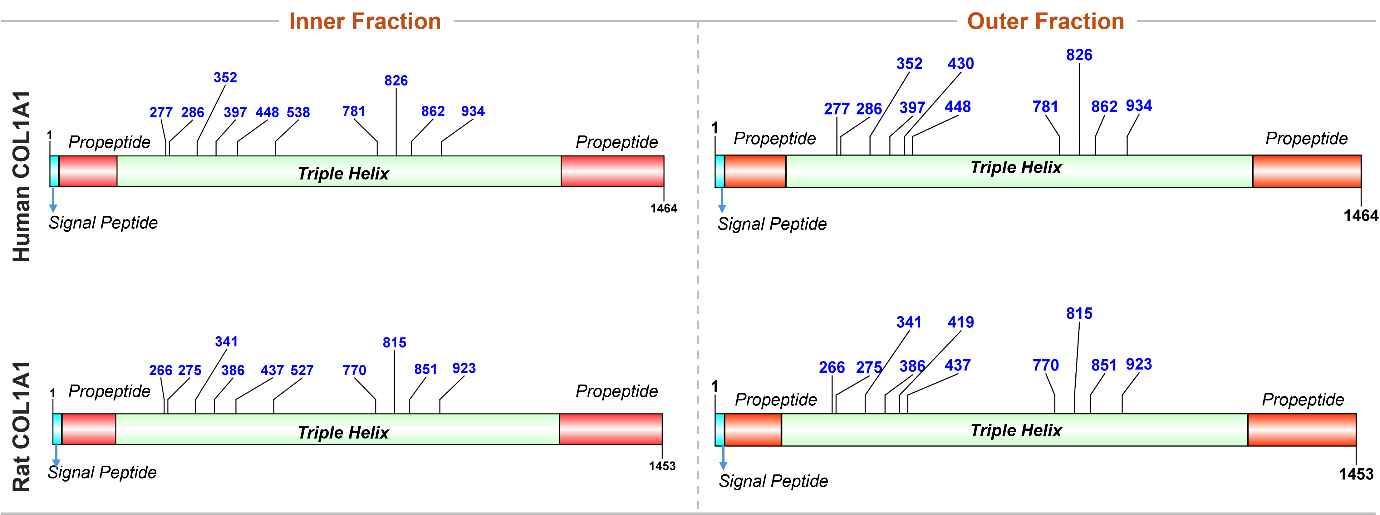

Figure S3. Comparative analysis of conserved HyK sites between human and rat COL1A1 chains. The rectangular box represents the COL1A1 chain that has signal peptide (light blue) propeptide (orange), and triple helix (light green) domains. All conserved sites (site numbers in blue) are depicted with the black lines.

**PSMs for OF, COL1A1**

**Modification: 3-HyP**

Site: P^333^

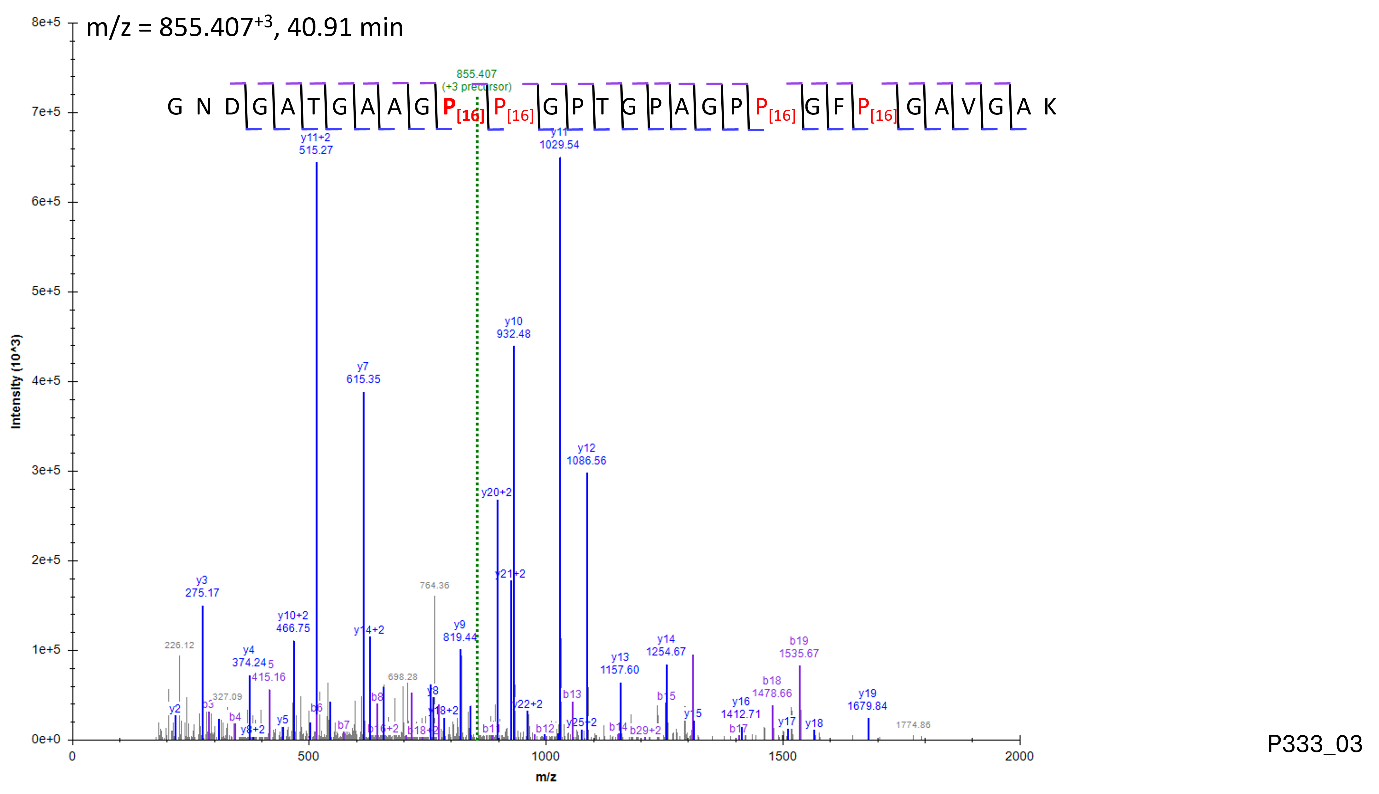

Site: P^375^

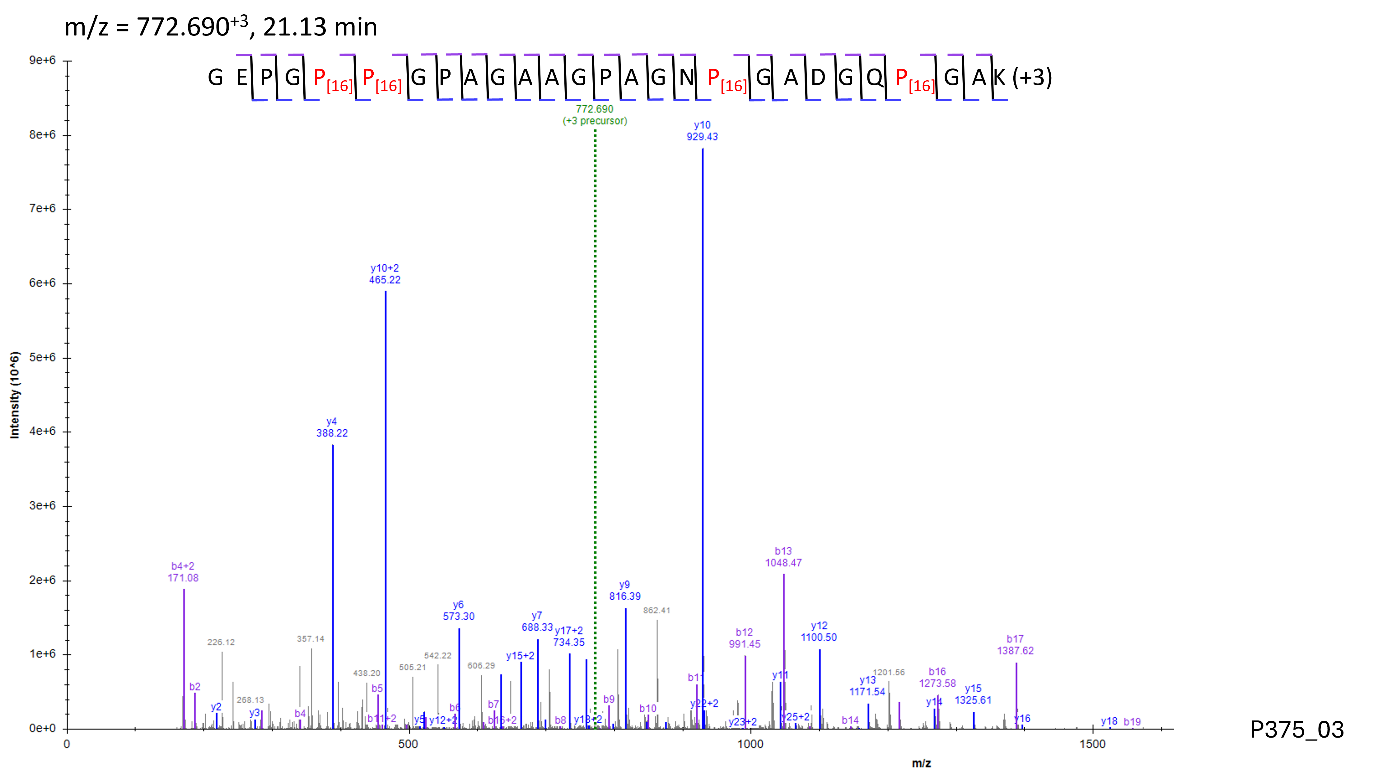

Site: P^459^

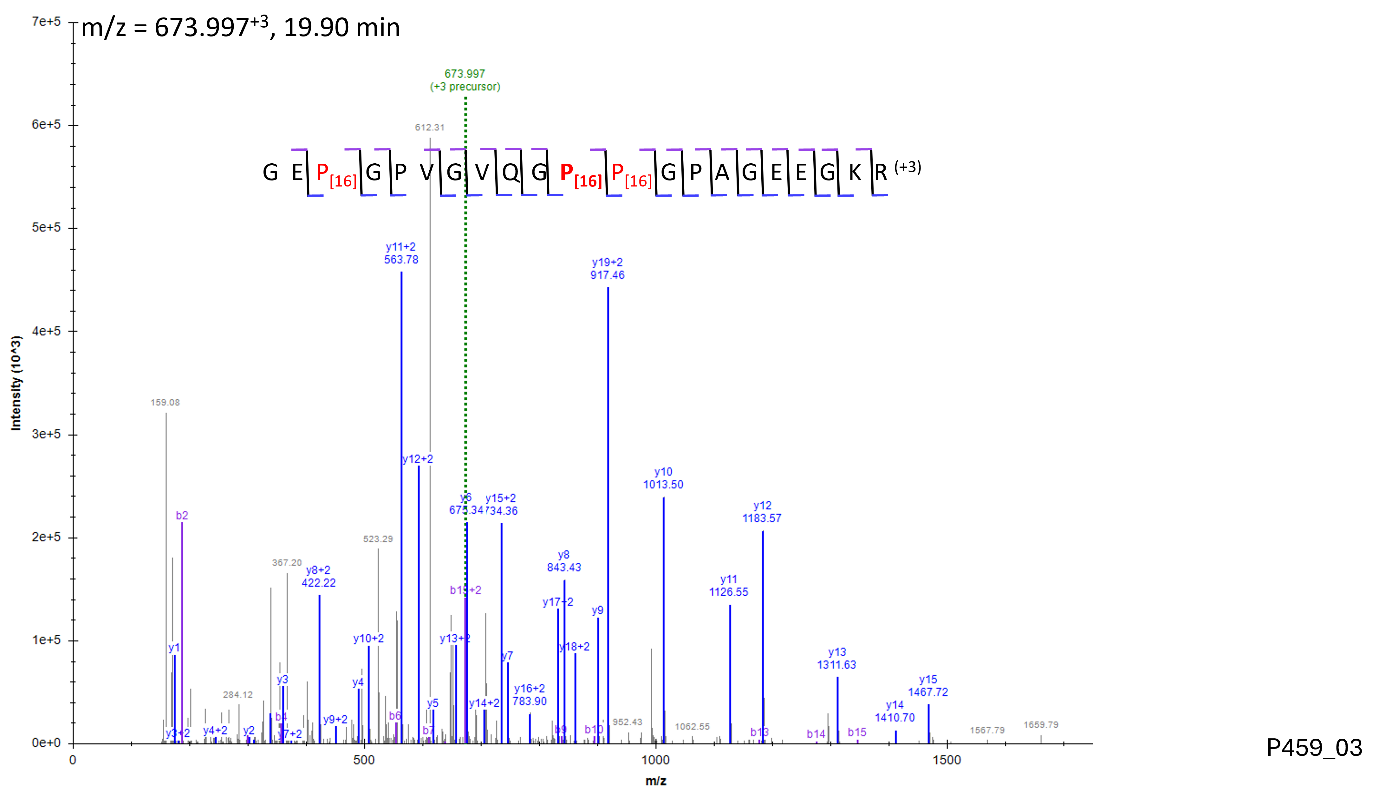

Site: P^555^

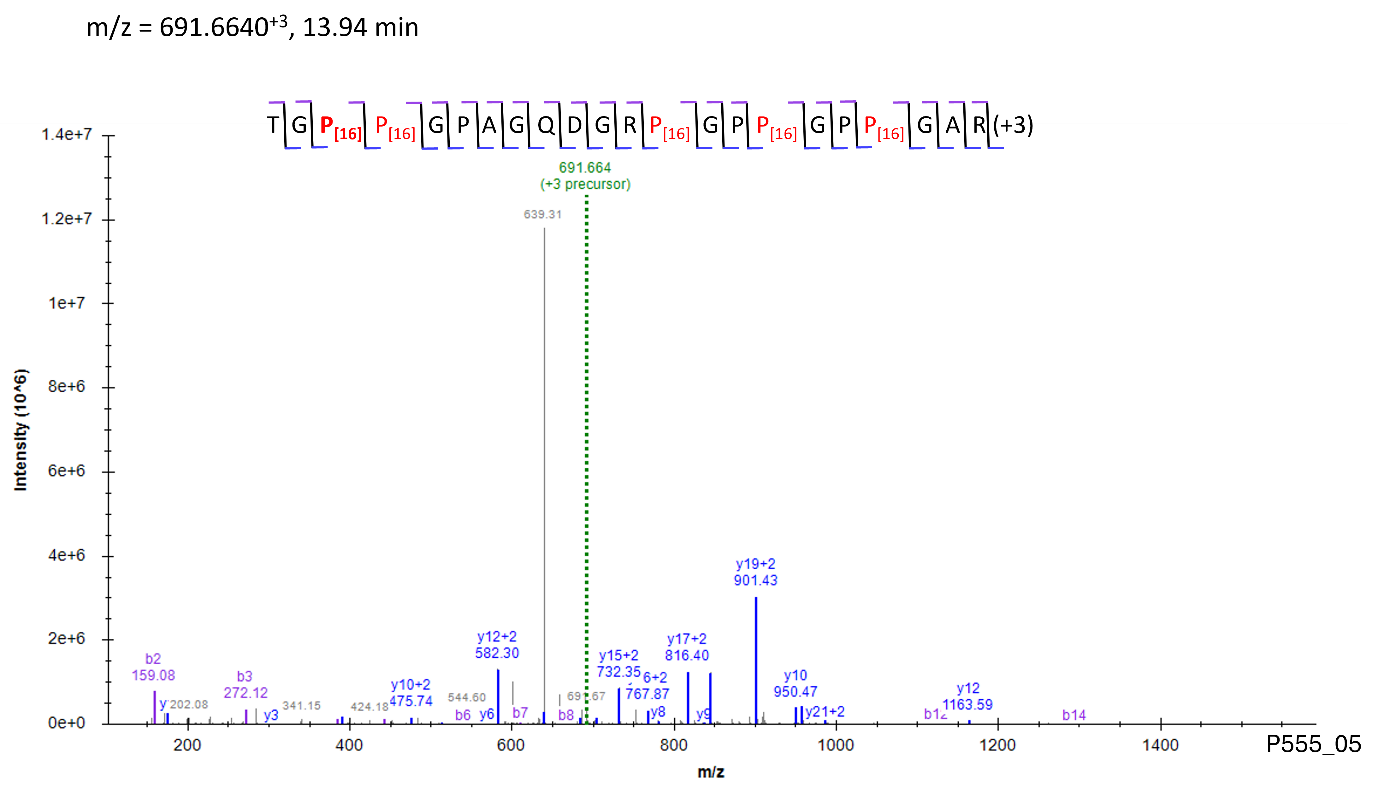

Site: P^567^

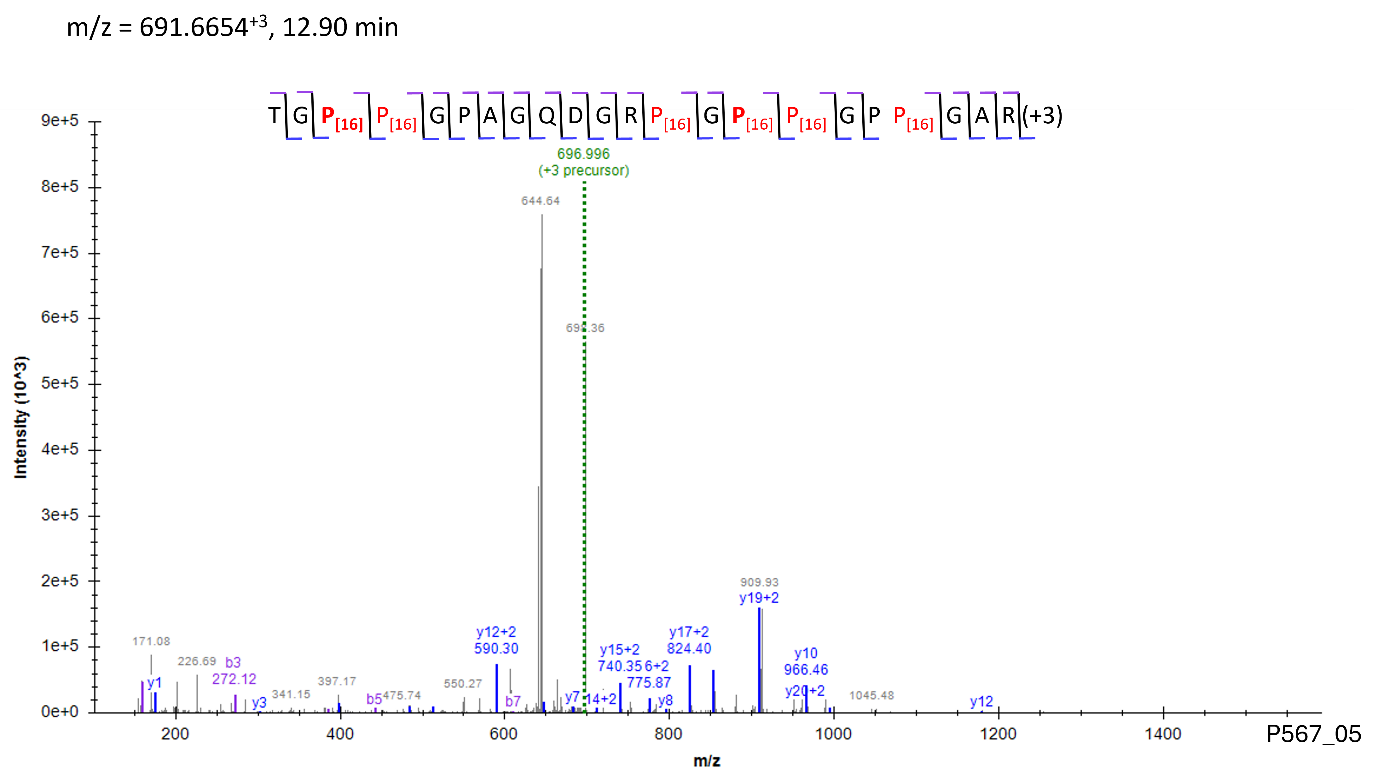

Site: P^570^

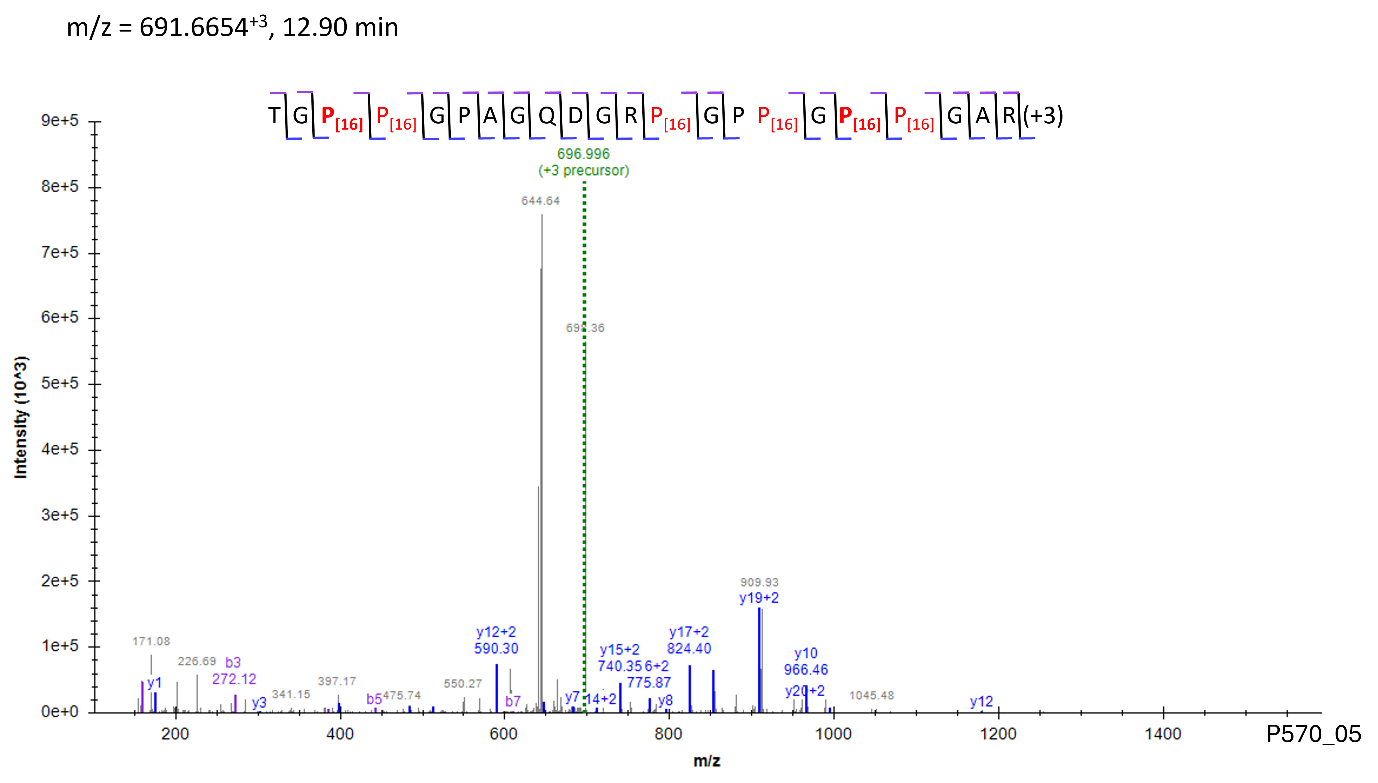

Site: P^771^

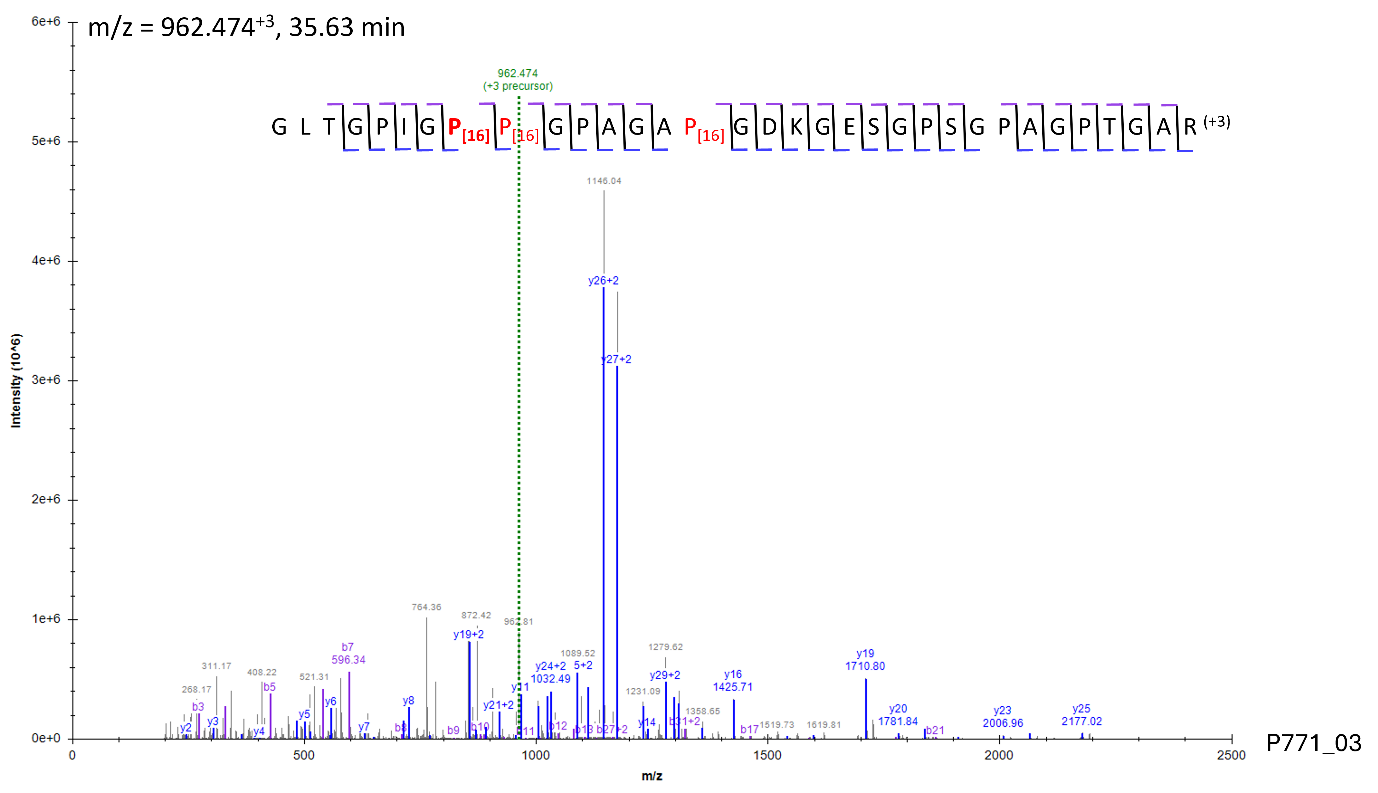

Site: P^807^

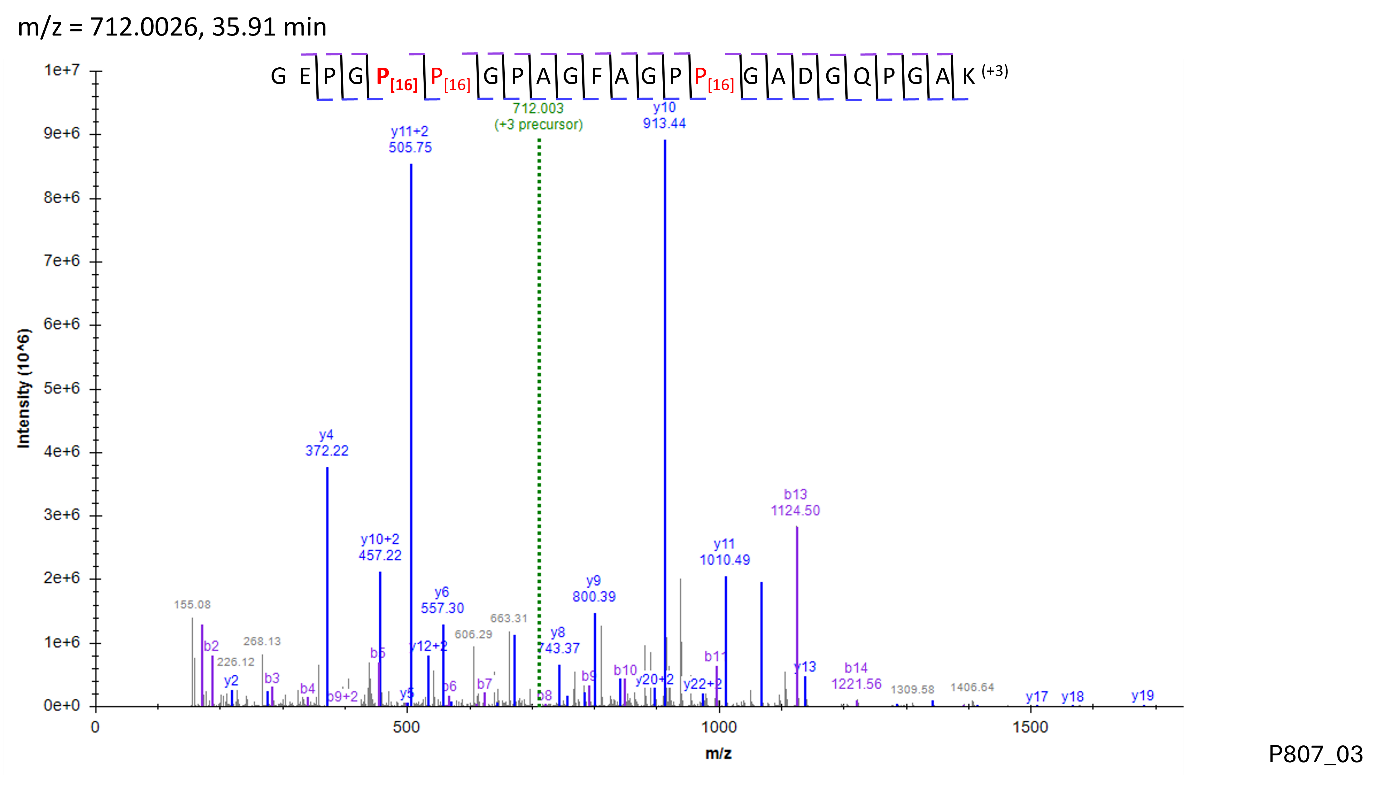

Site: P^816^

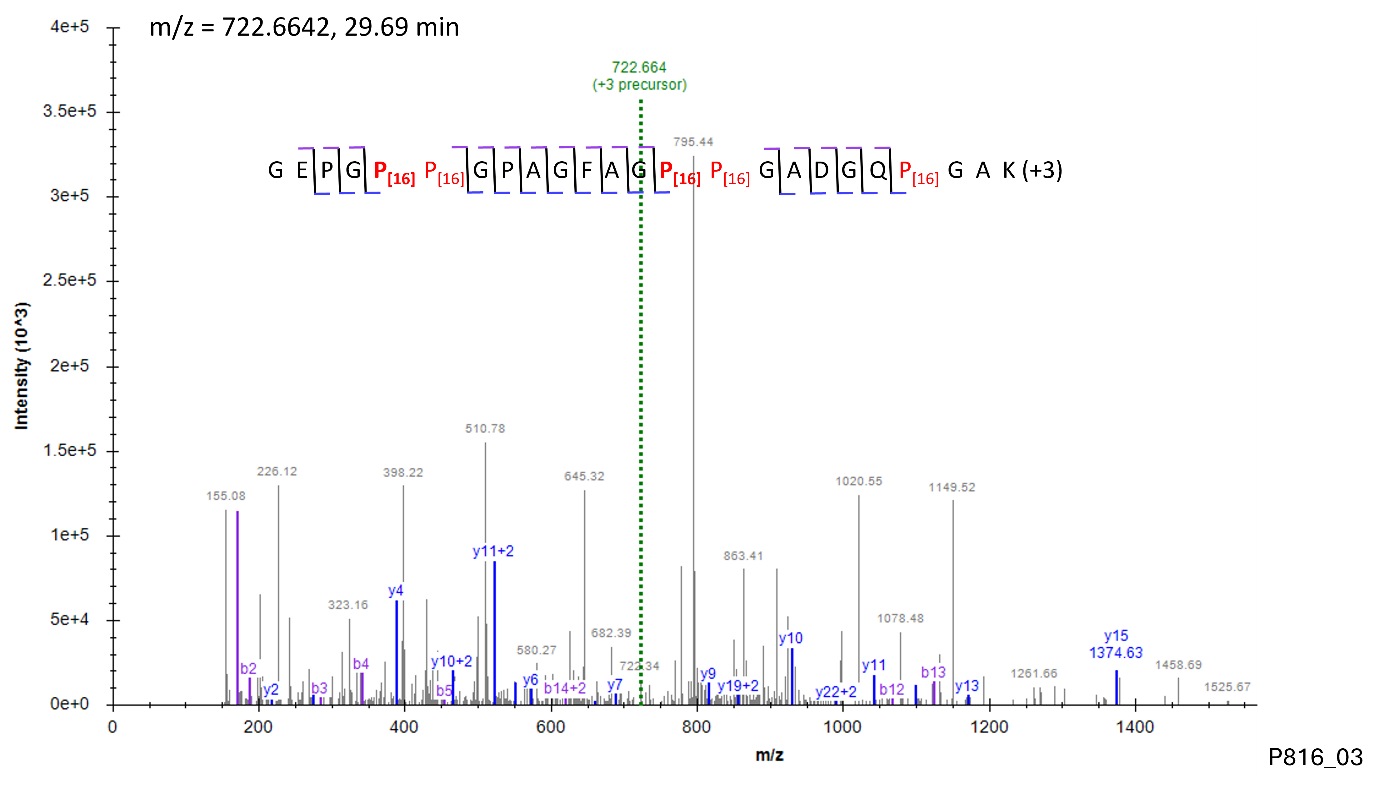

Site: P^840^

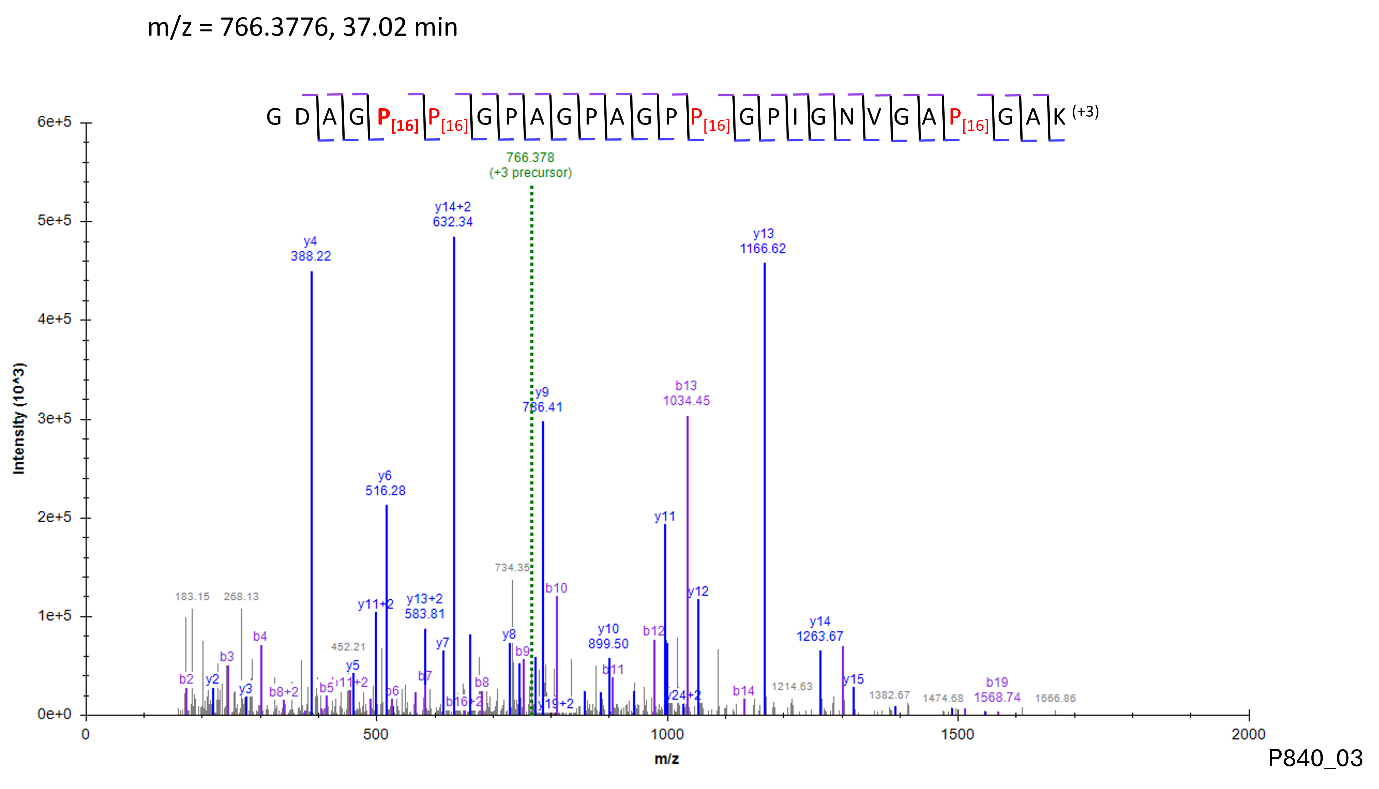

Site: P^870^

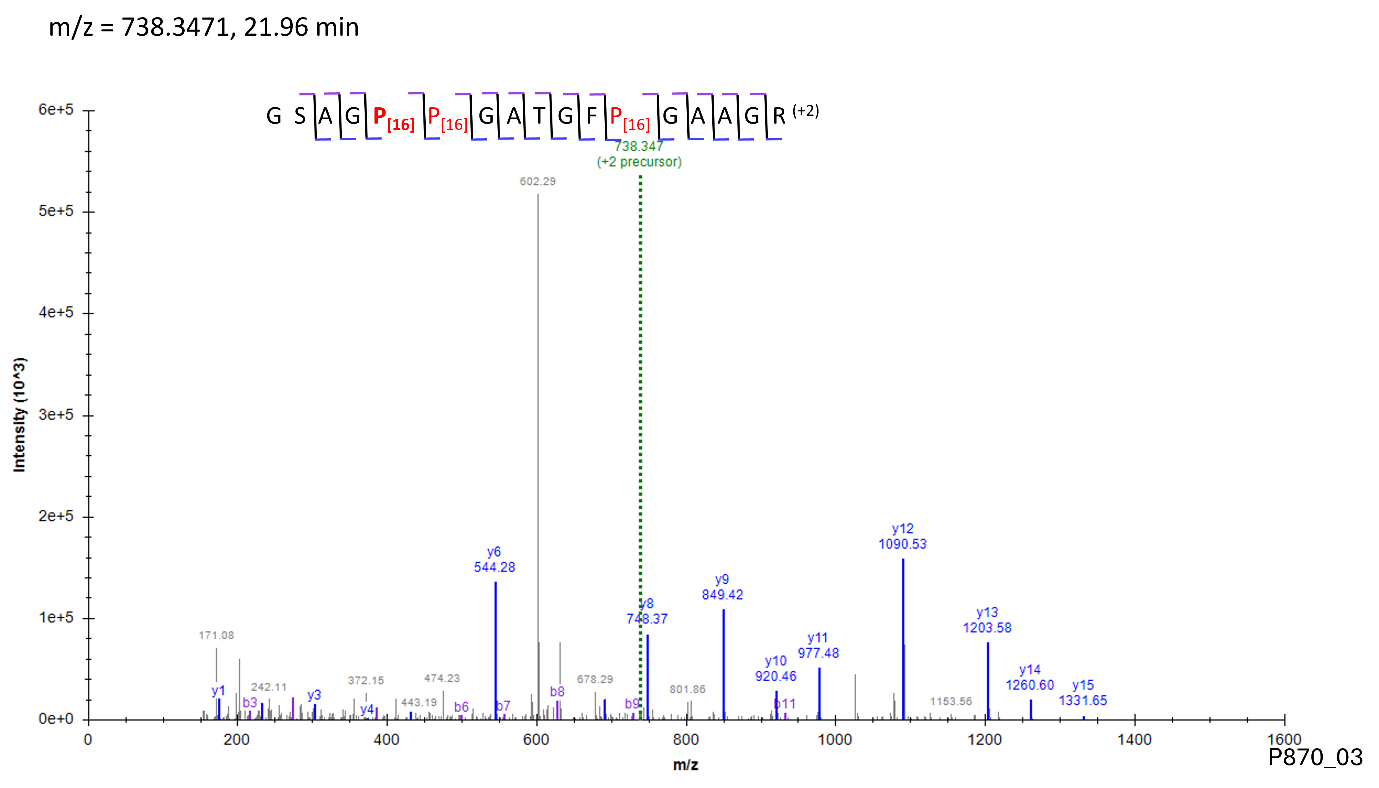

Site: P^885^

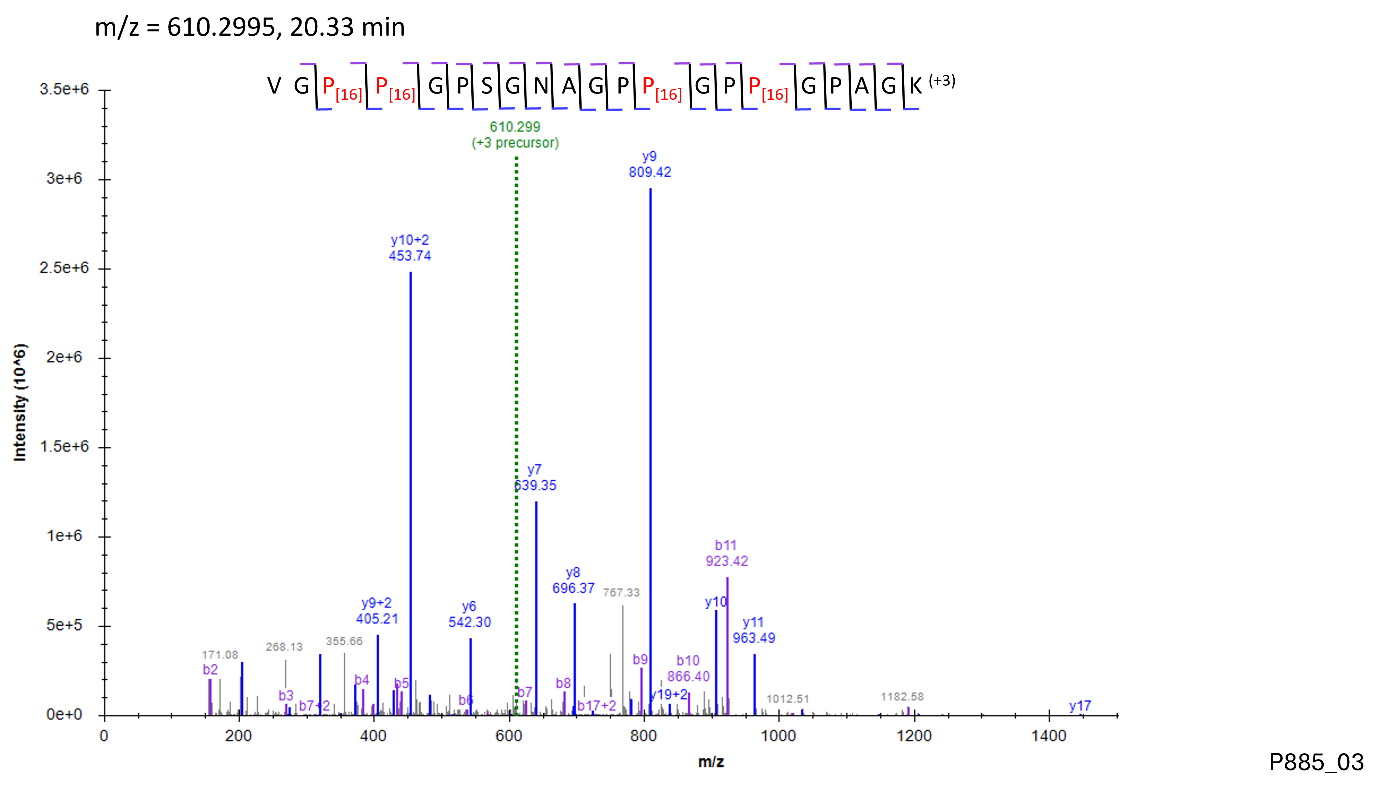

Site: P^894^

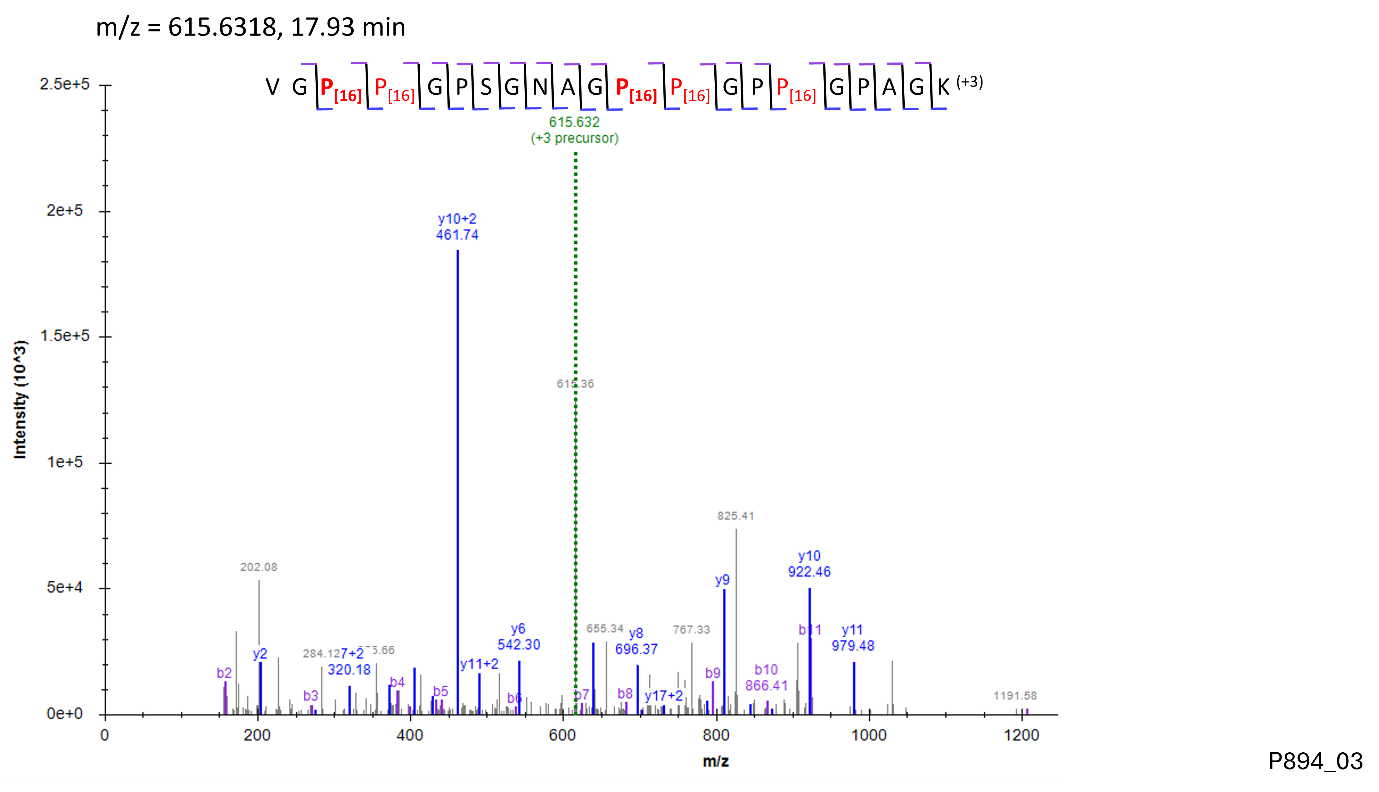

Site: P^897^

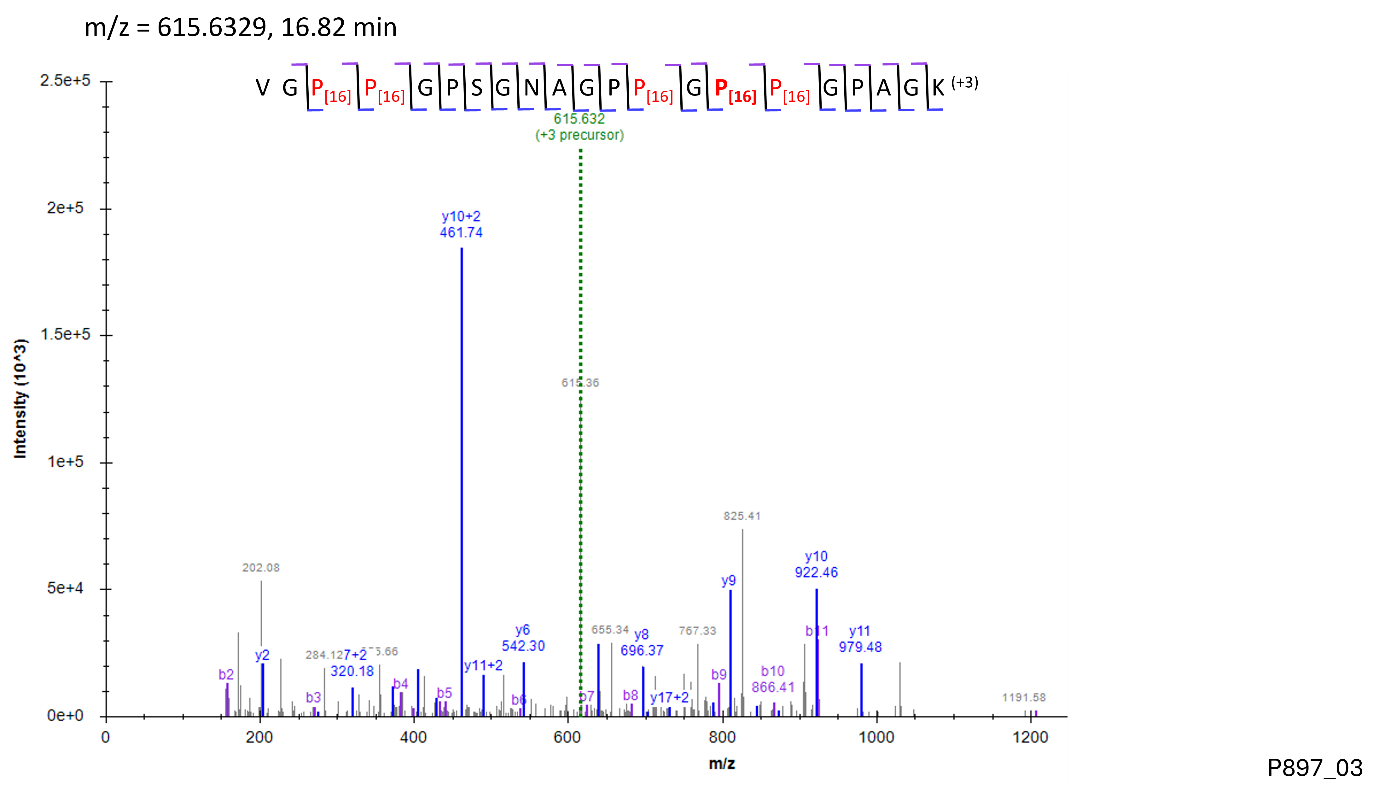

Site: P^924^

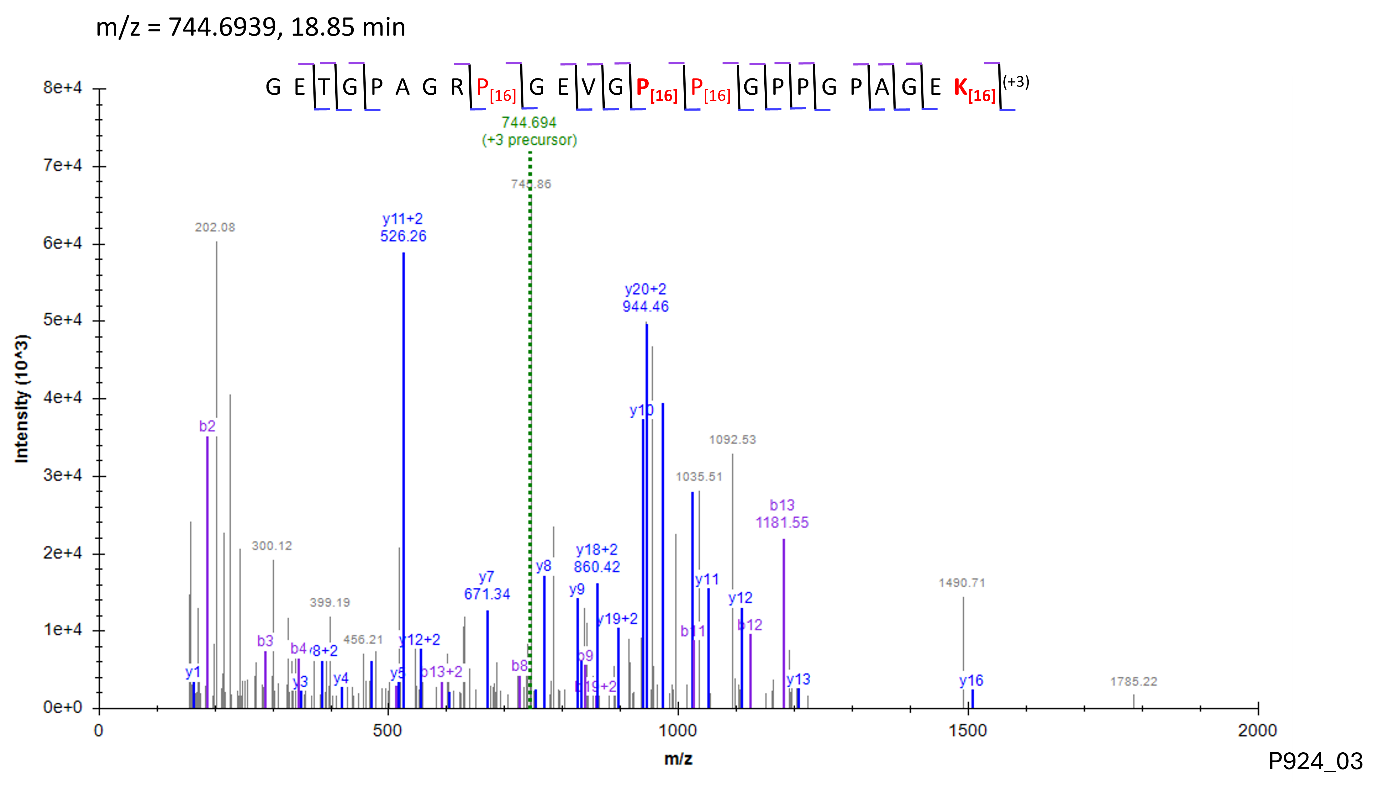

Site: P^927^

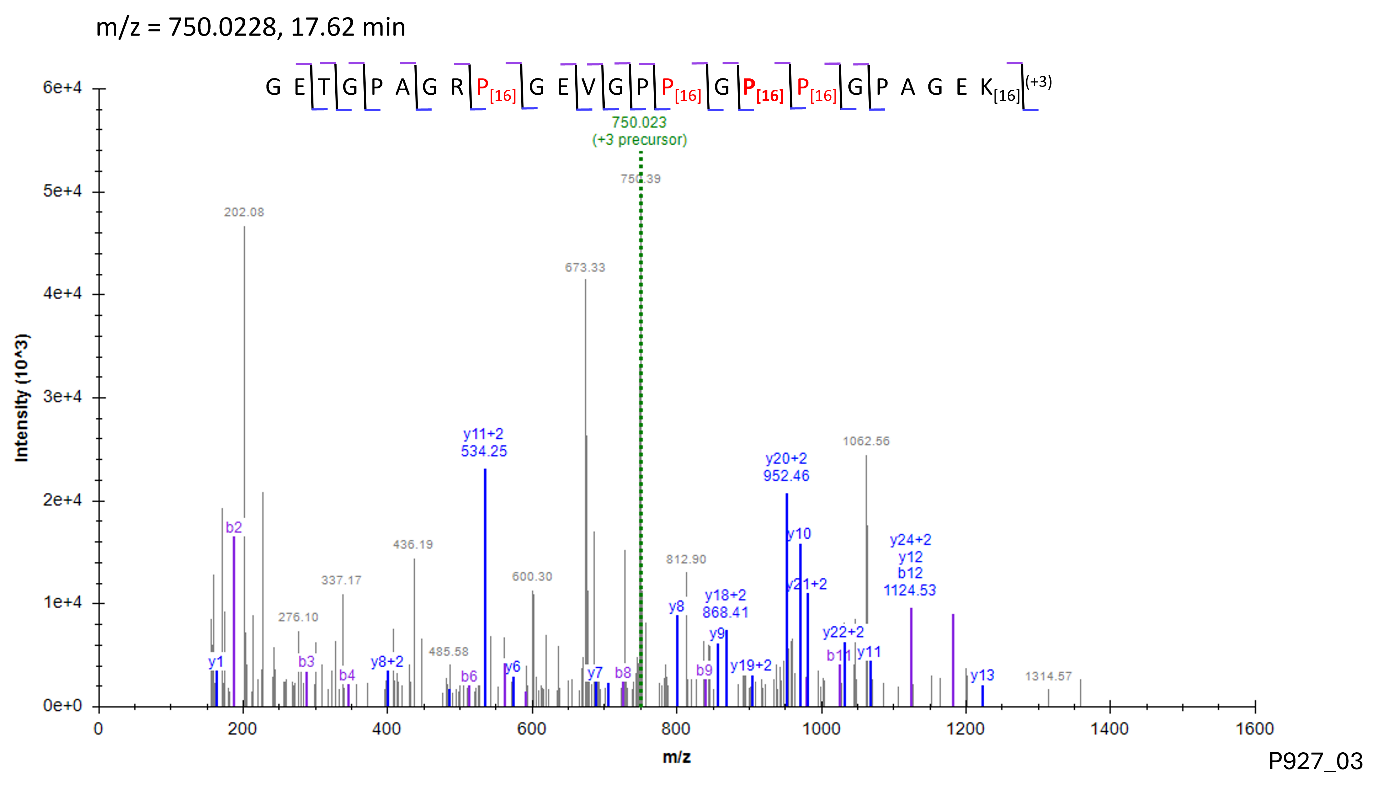

Site: P^996^

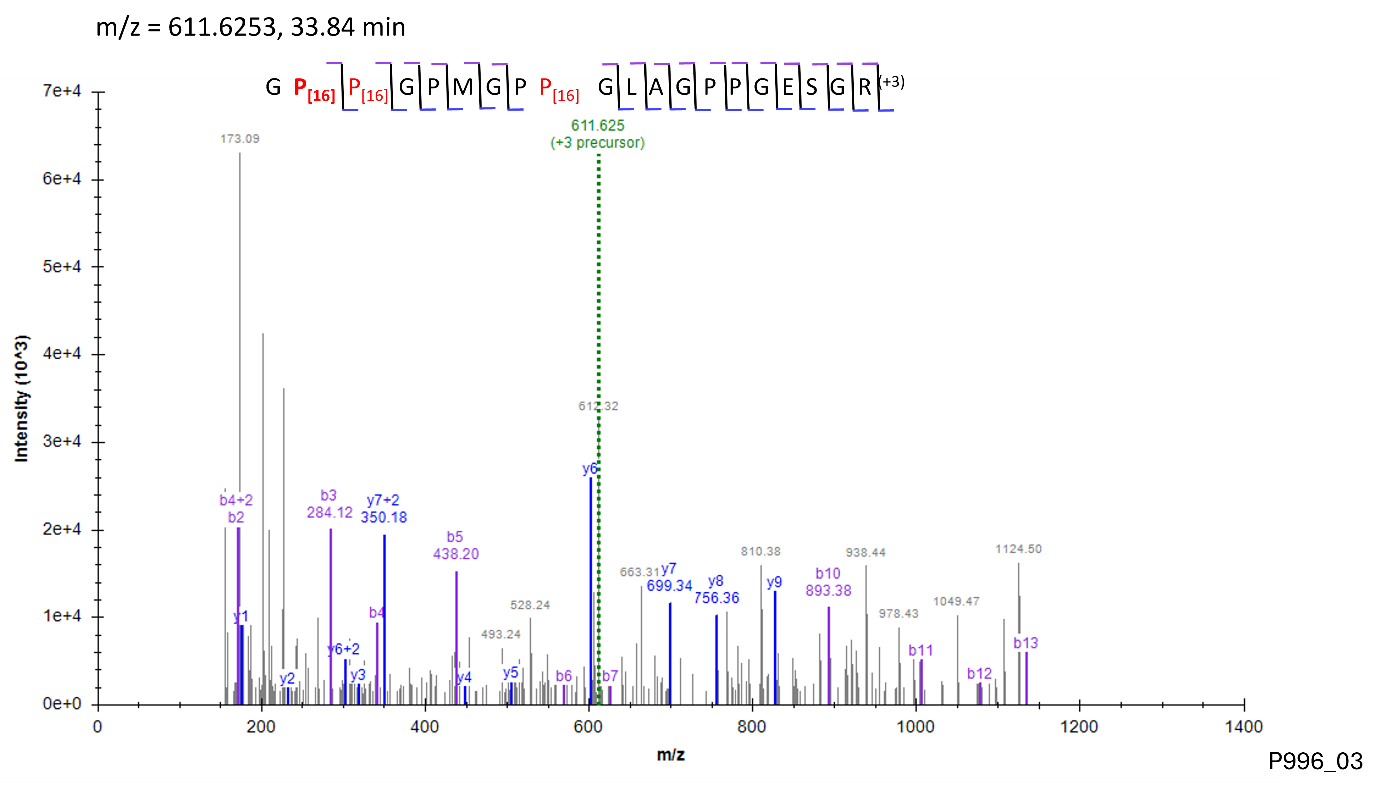

Site: P^1044^

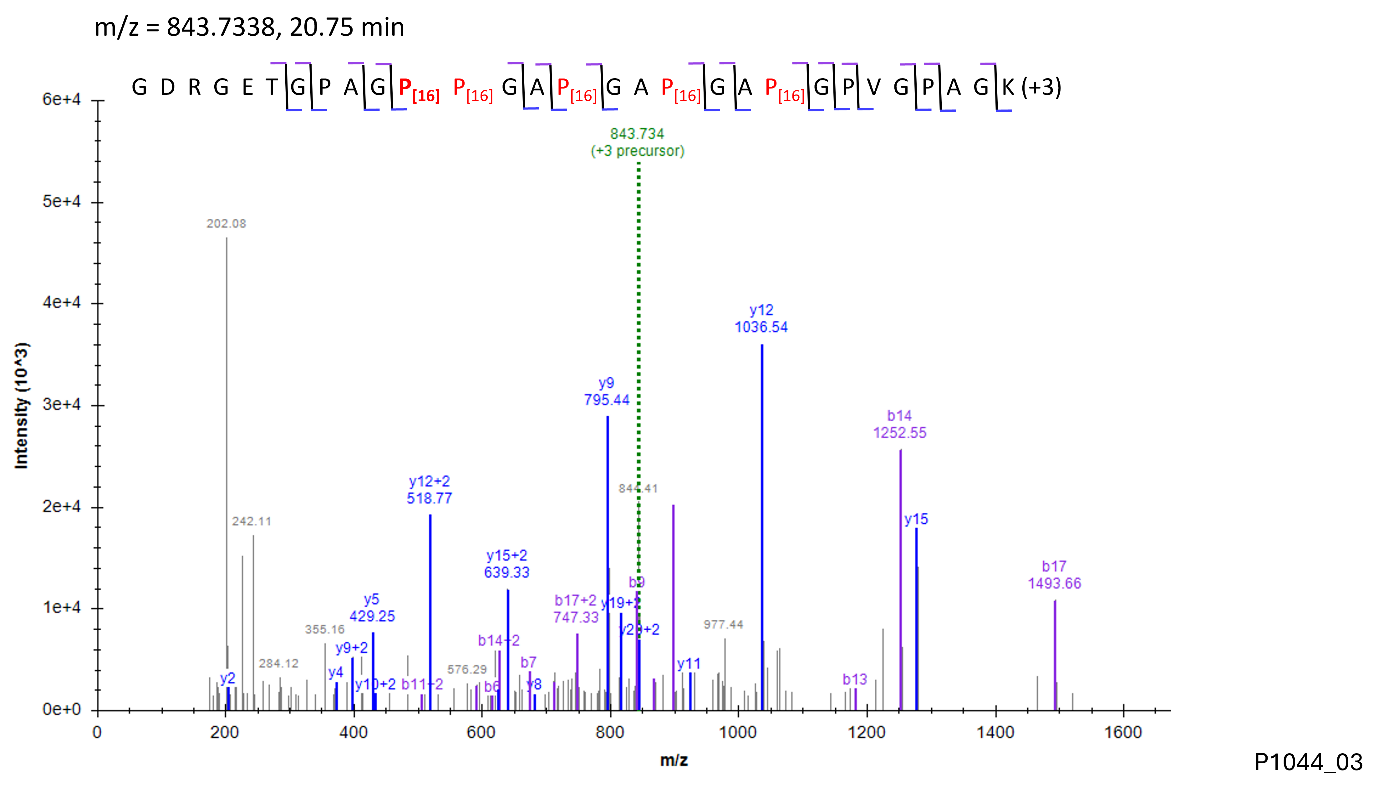

Figure – S18

Site: P^1119^, Site: P^1122^

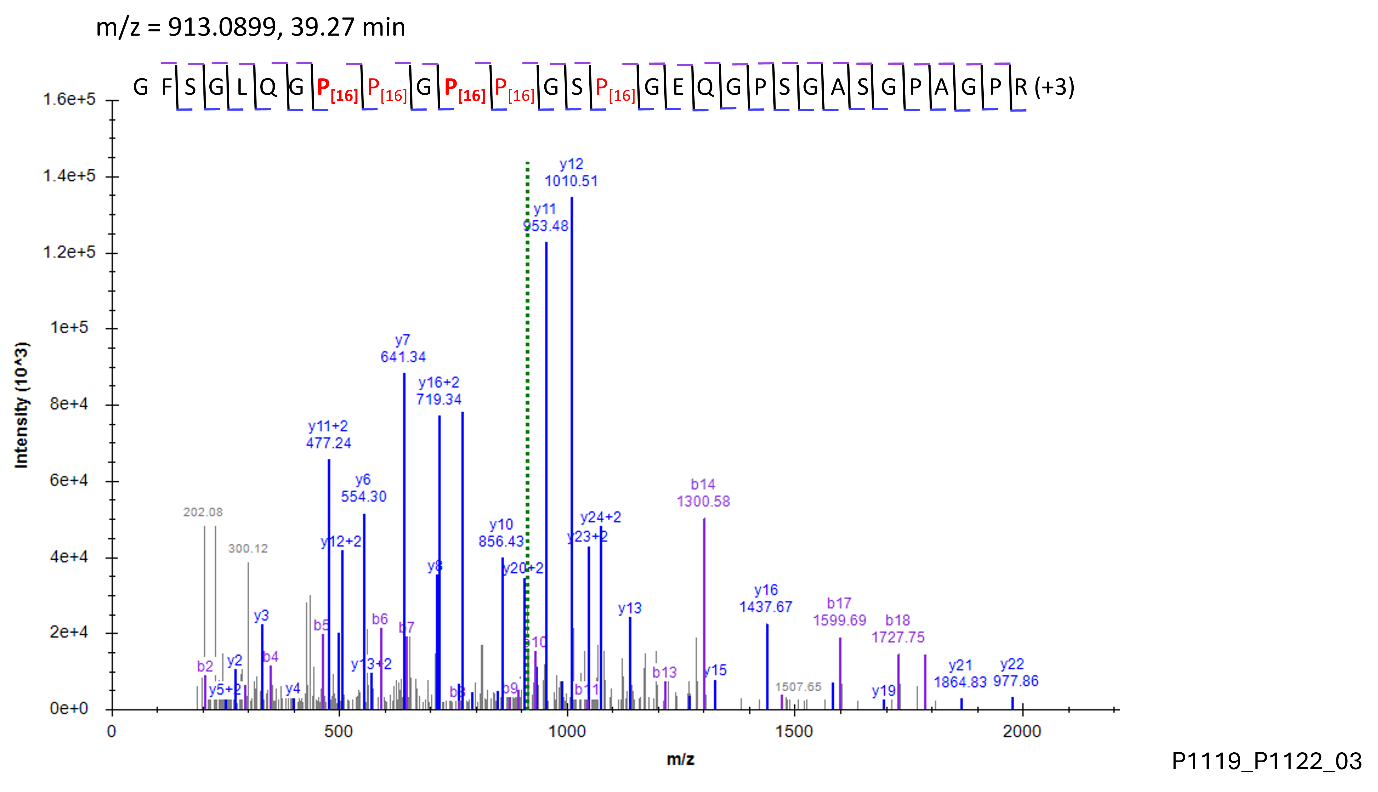

**Modification: HyK**

Site: K^277^

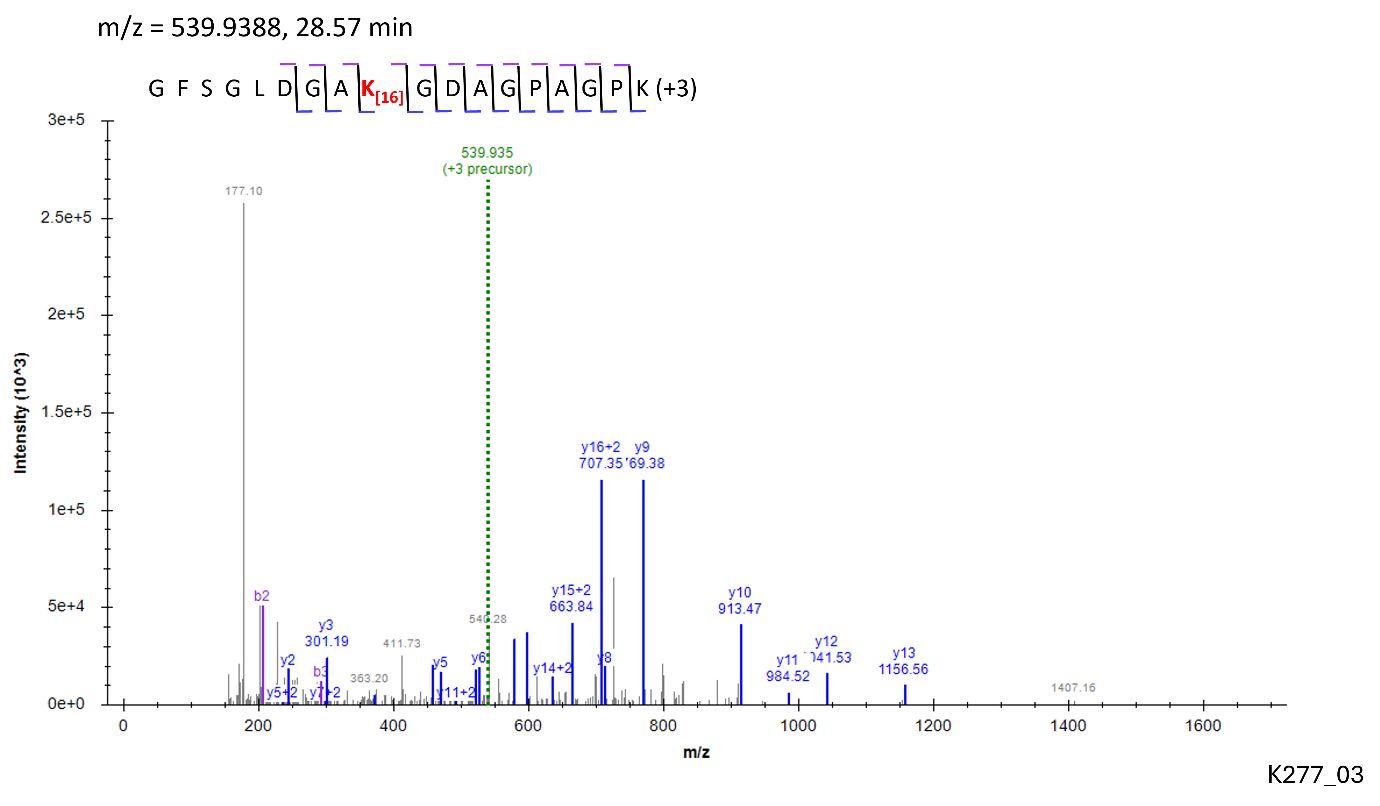

Site: K^286^

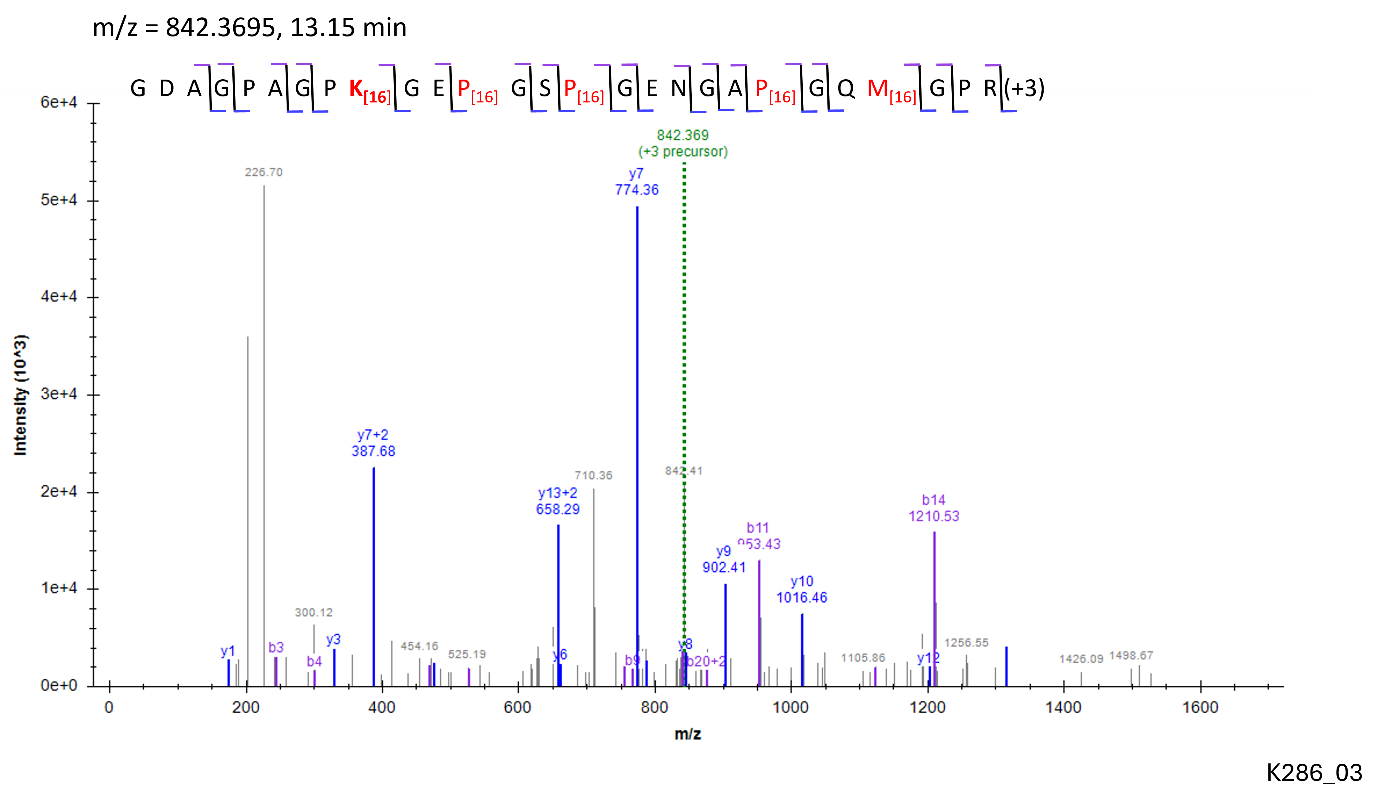

Site: K^397^

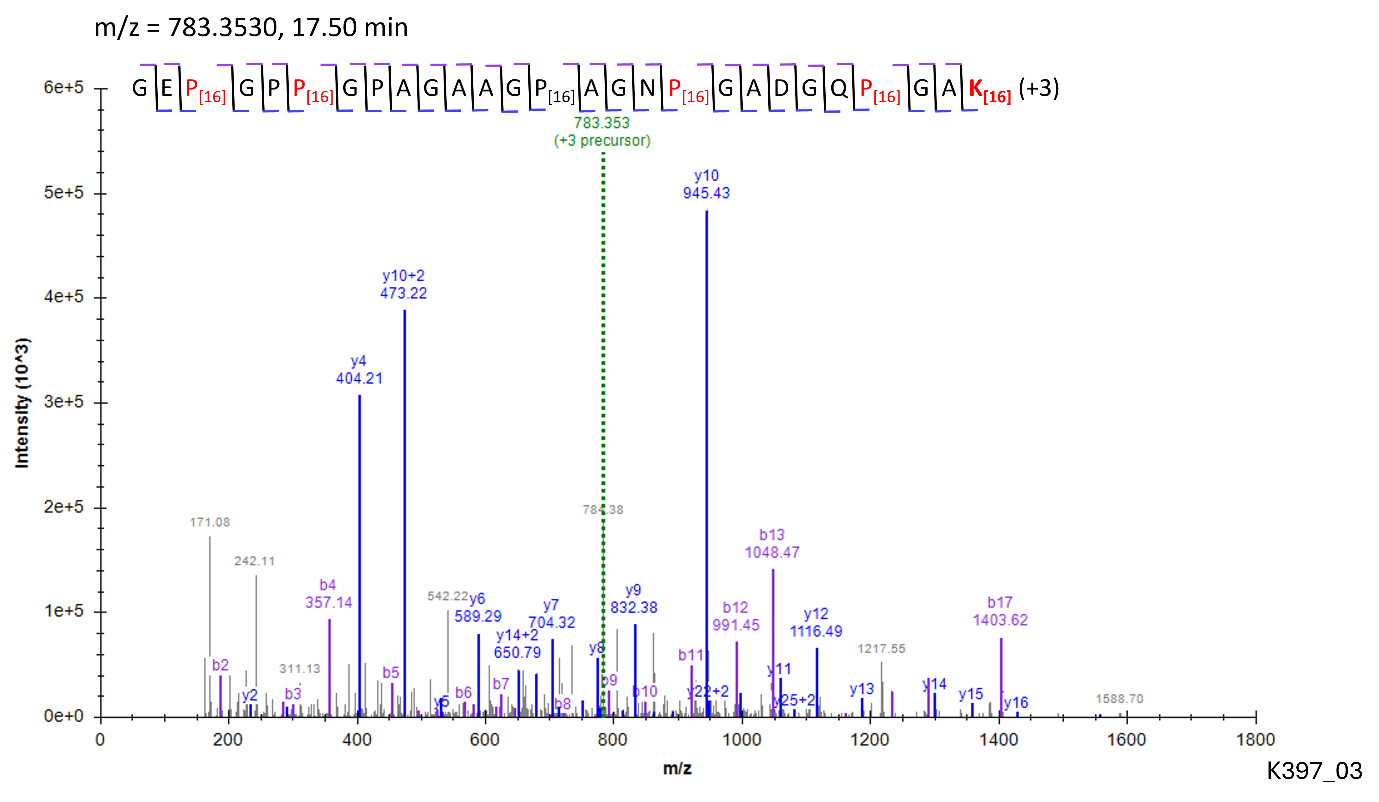

Site: K^430^

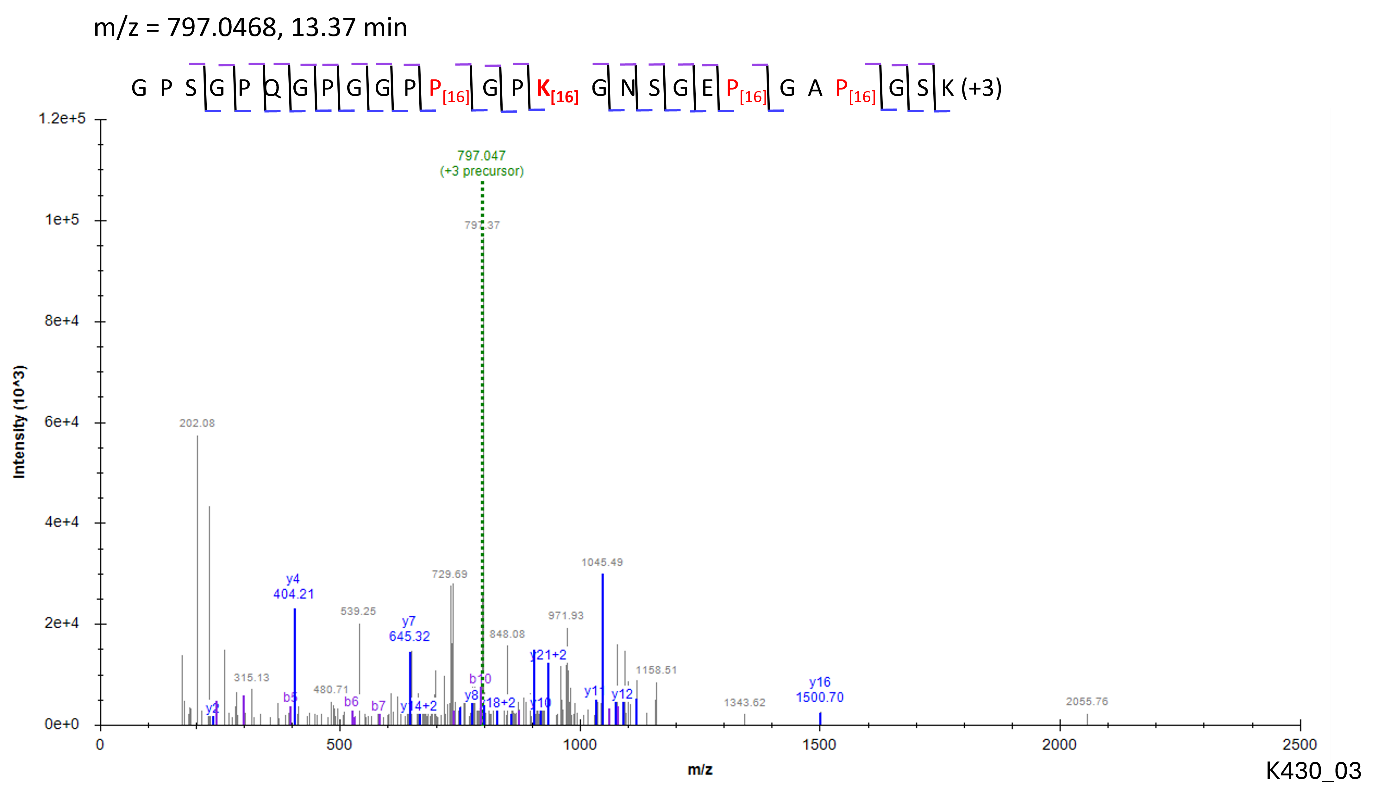

Site: K^448^

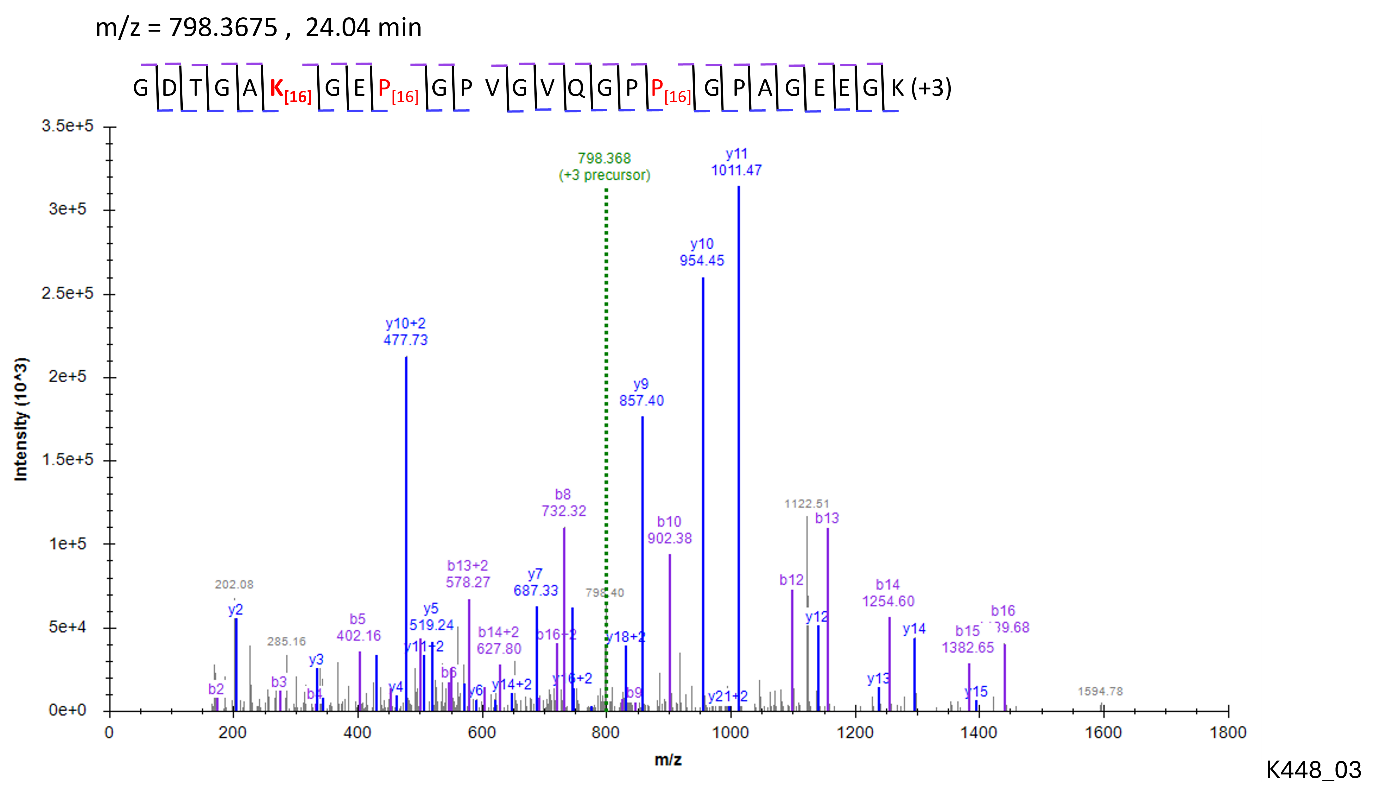

Site: K^586^

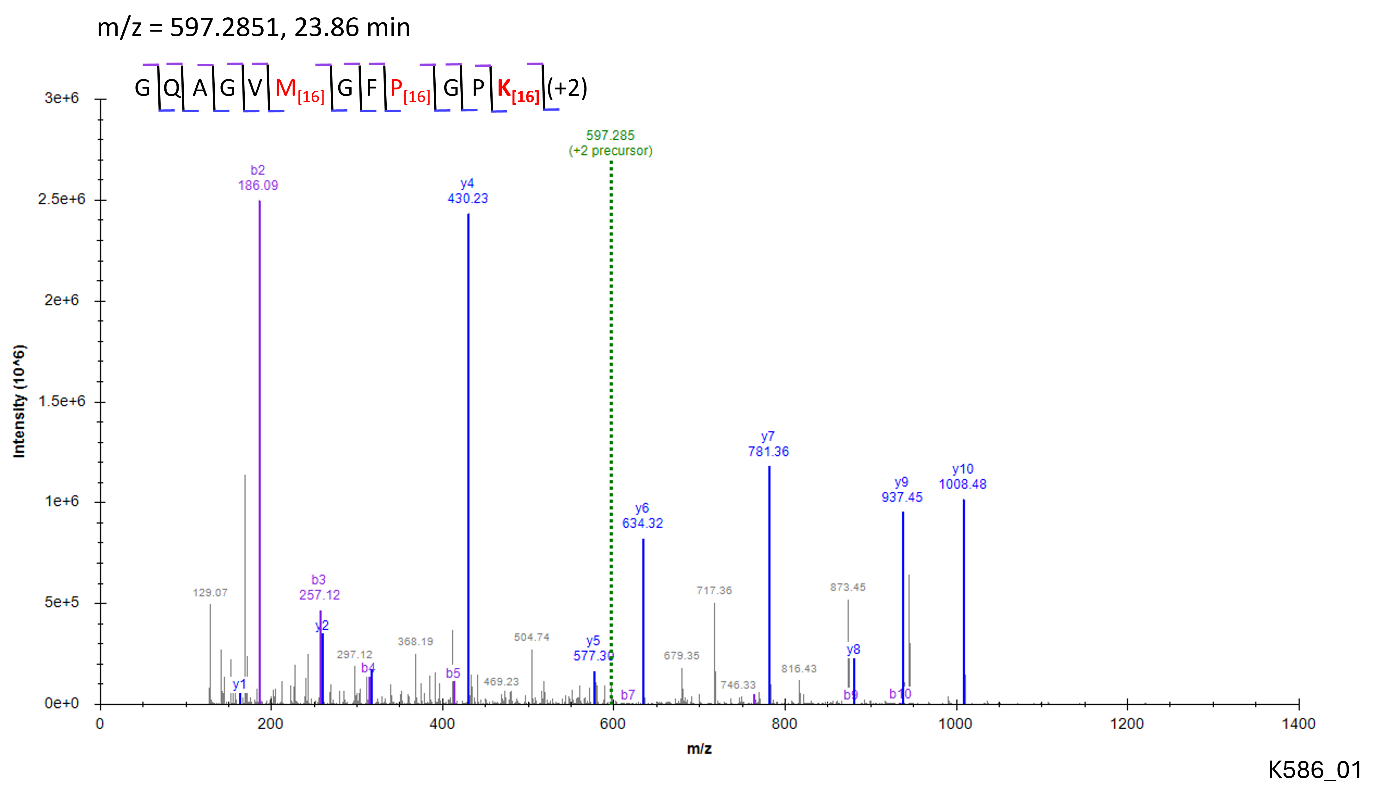

Site: K^781^

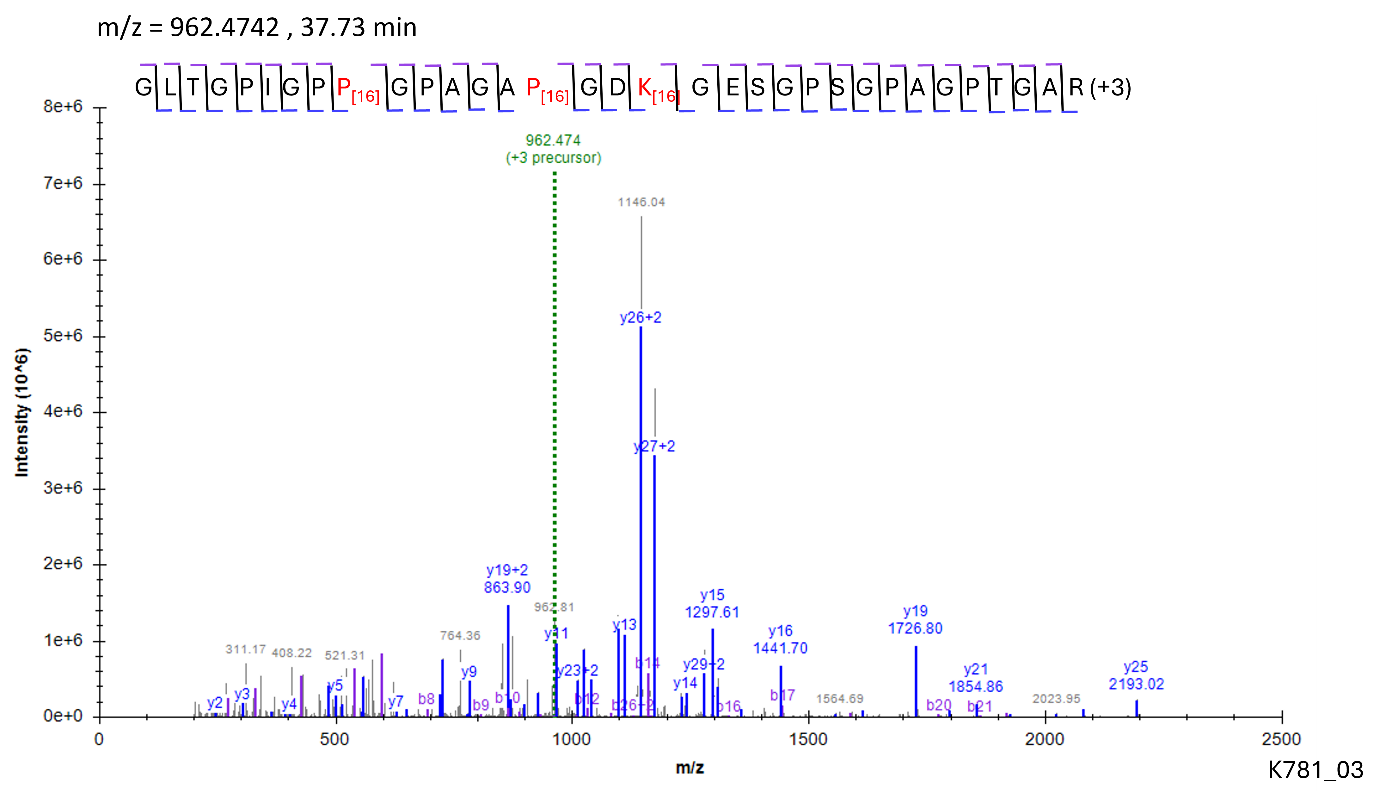

Site: K^826^

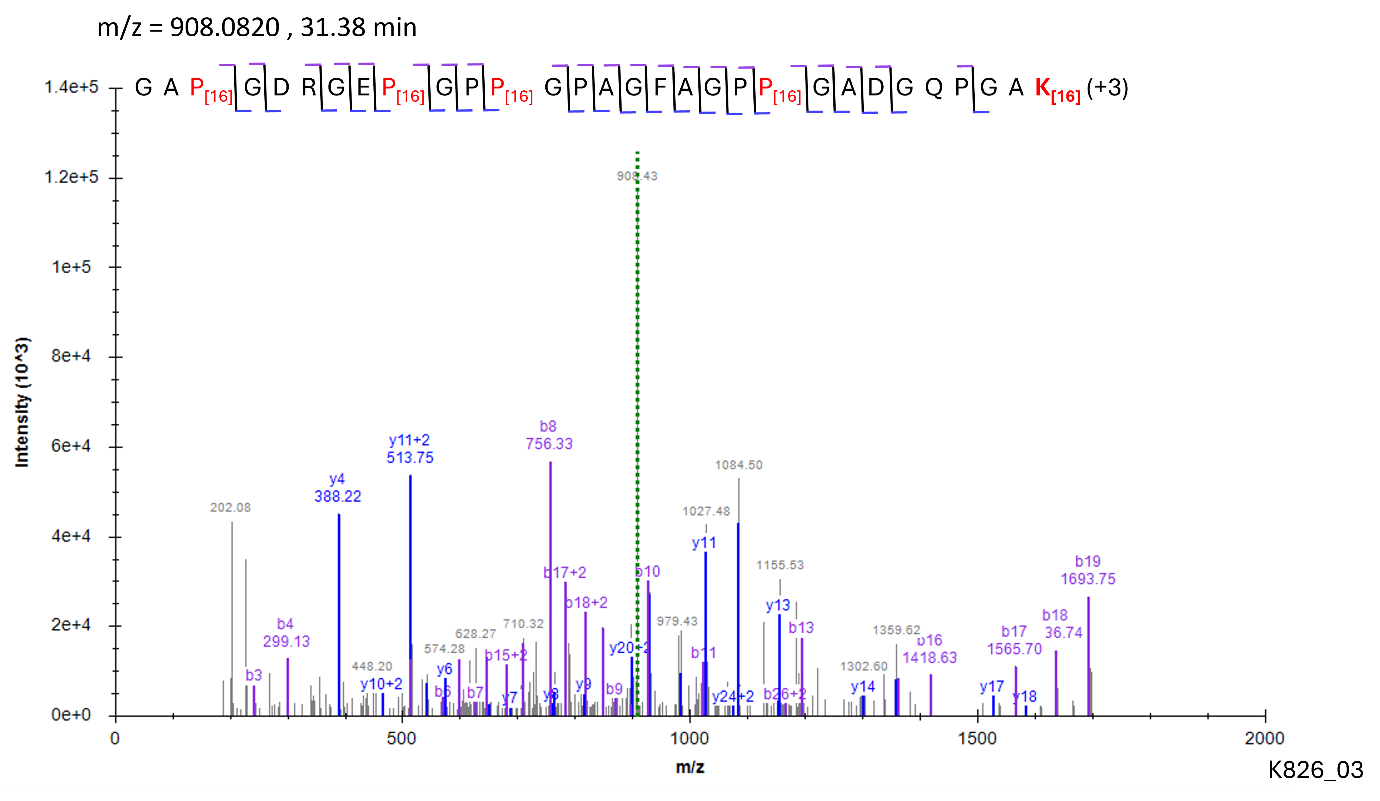

Site: K^862^

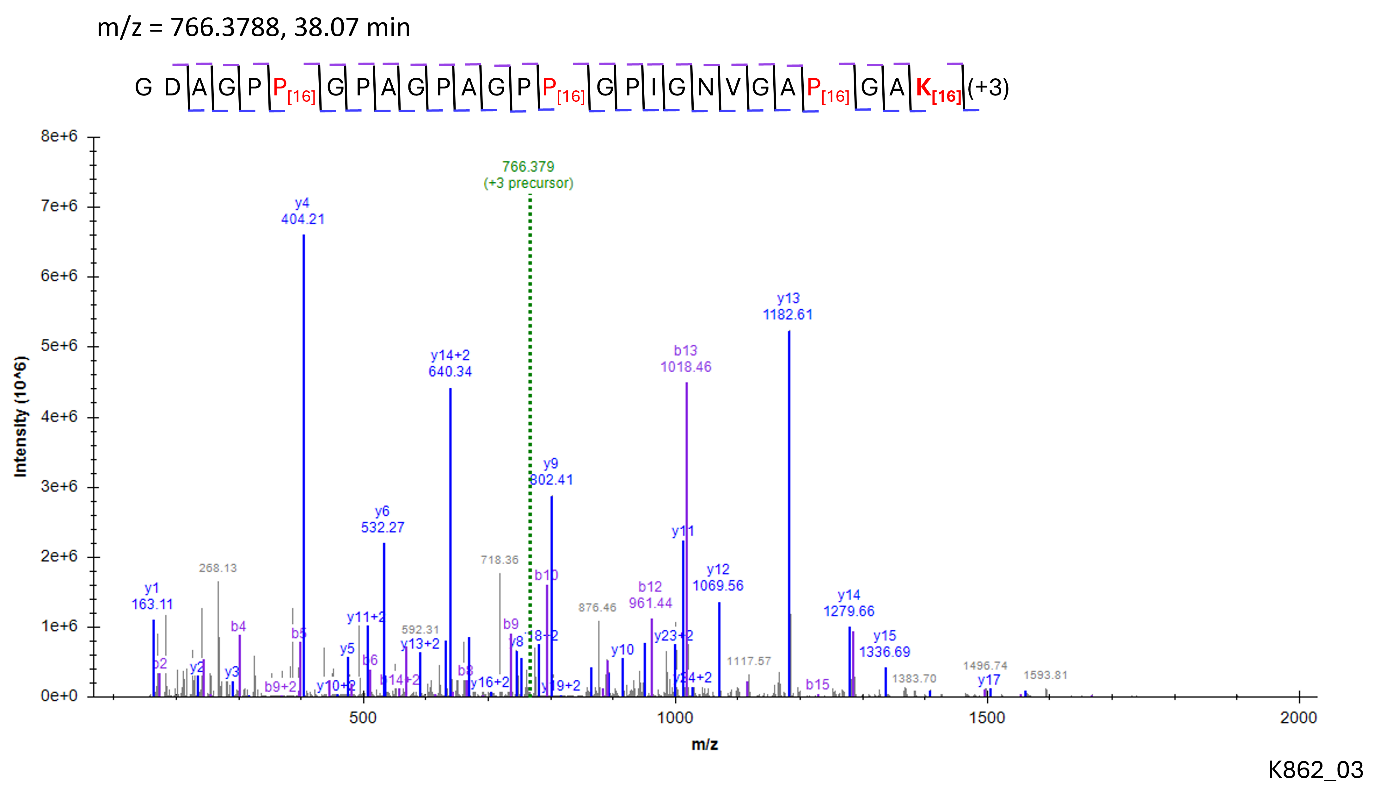

Site: K^934^

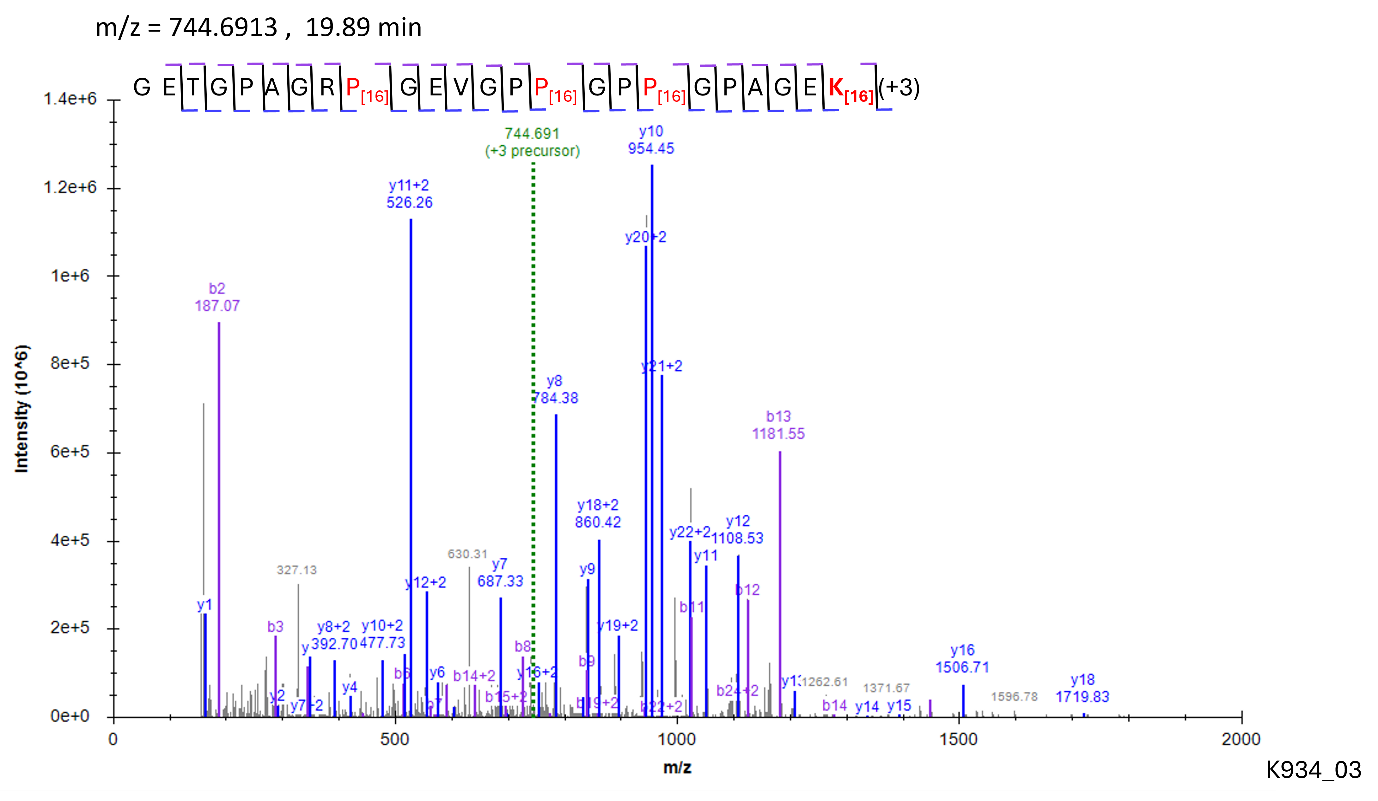

**Modification: G-HyK**

Site: K^286^

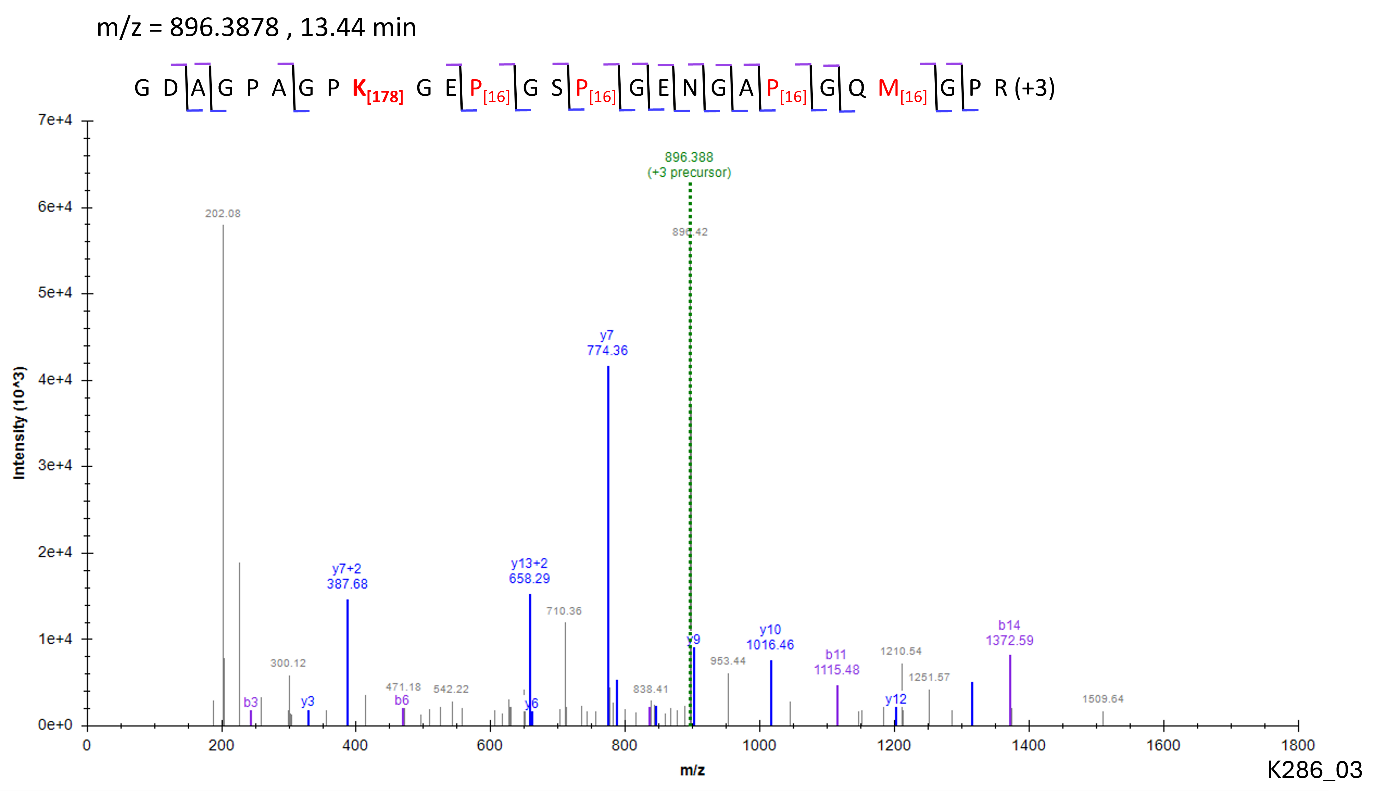

Figure – S30

Site: K^862^

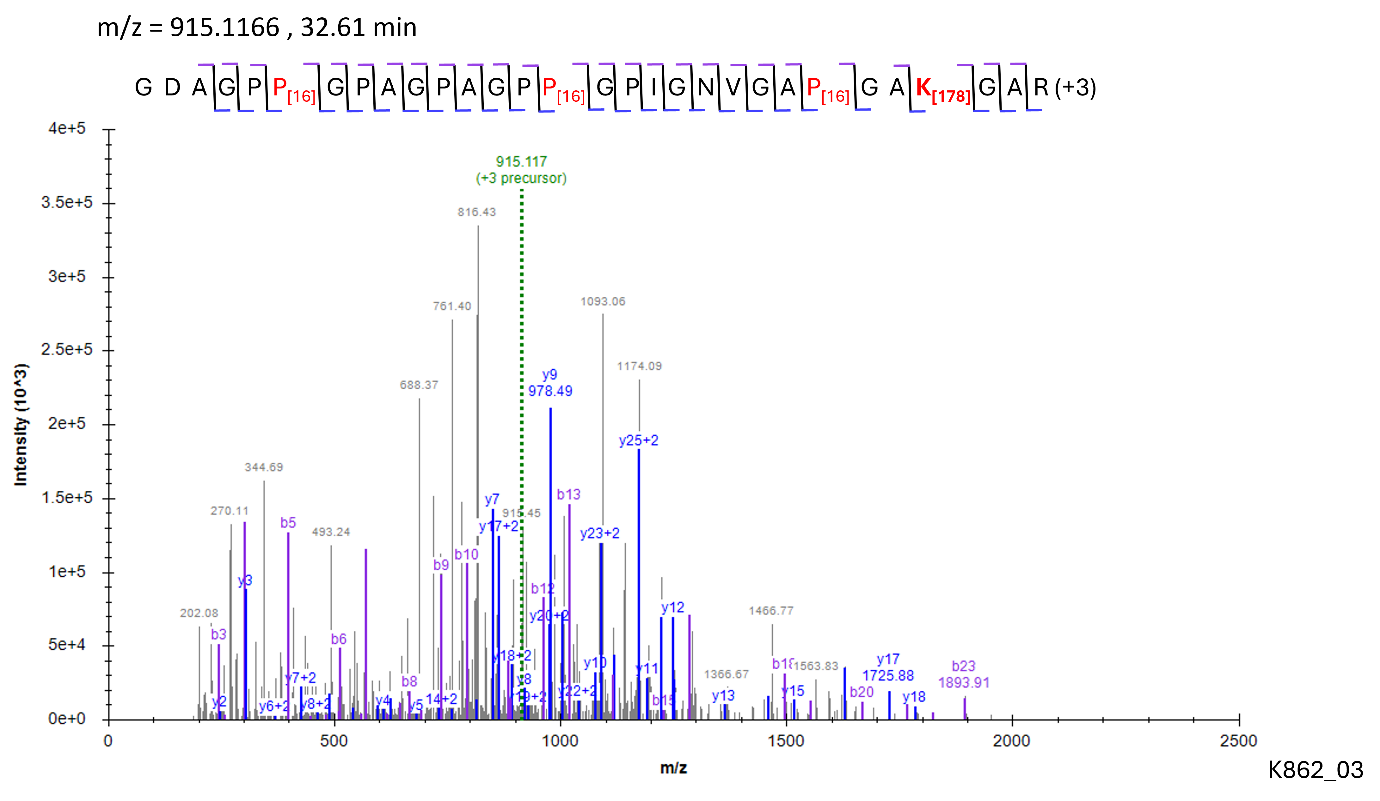

**Modification: GG-HyK**

Site: K^862^

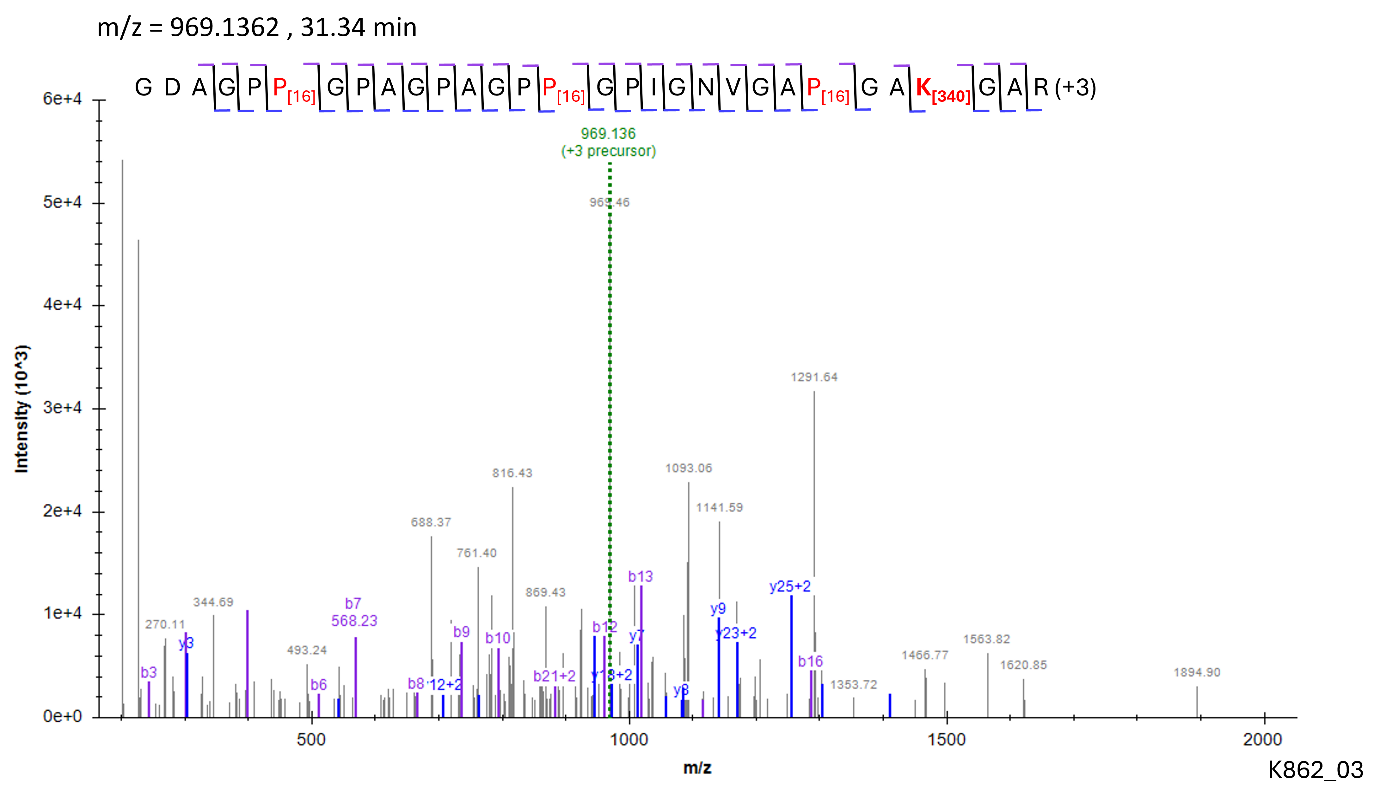

**PSMs for IF, COL1A1**

**Modification: 3-HyP**

Site: P^333^

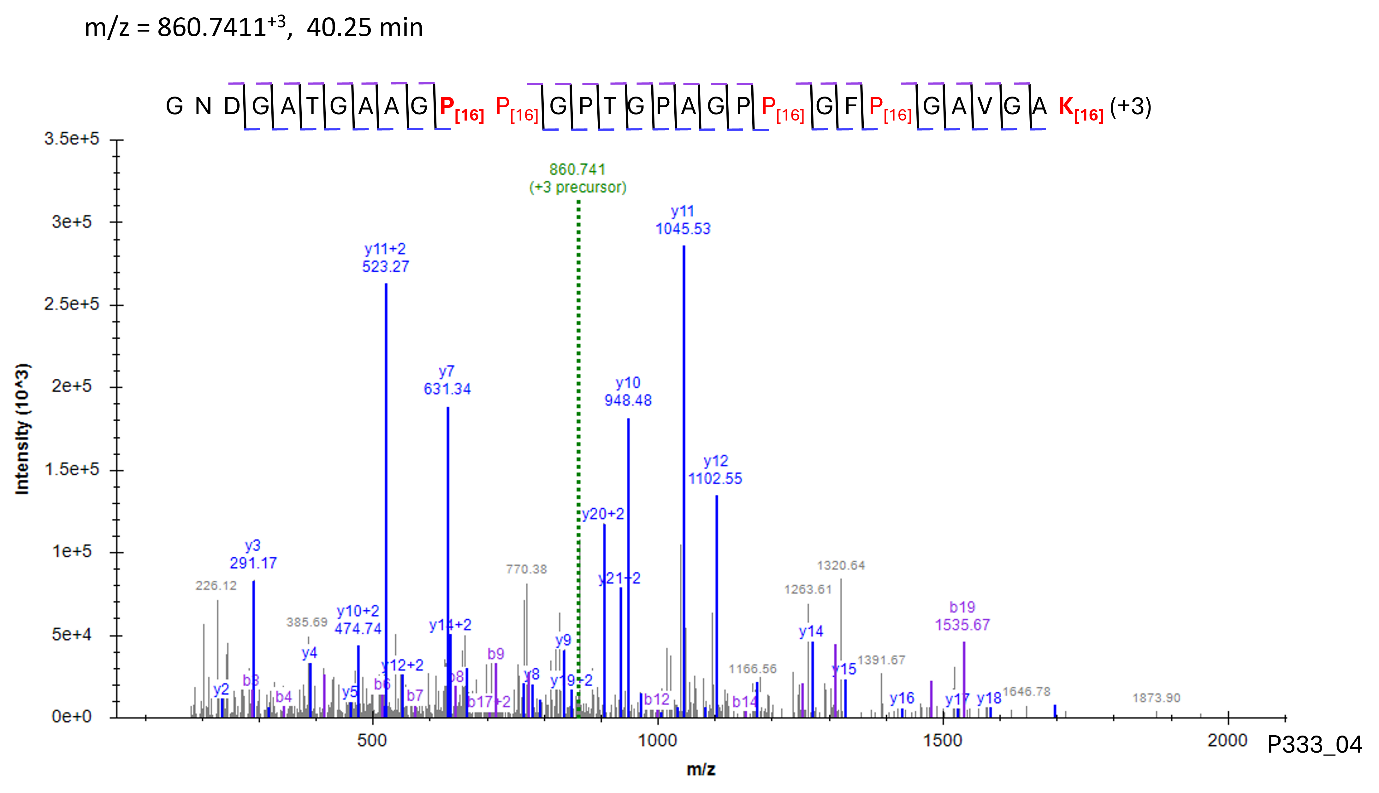

Site: P^375^

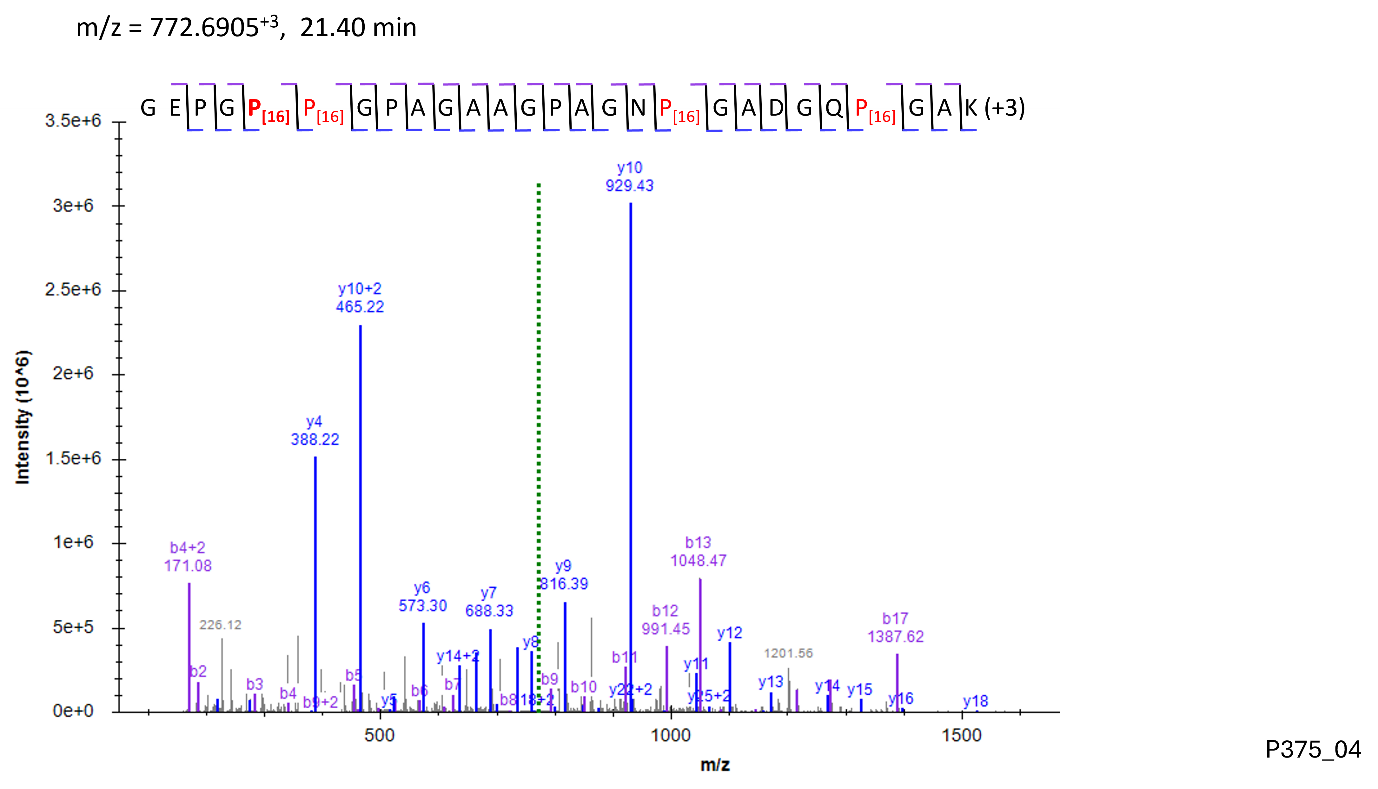

Site: P^426^

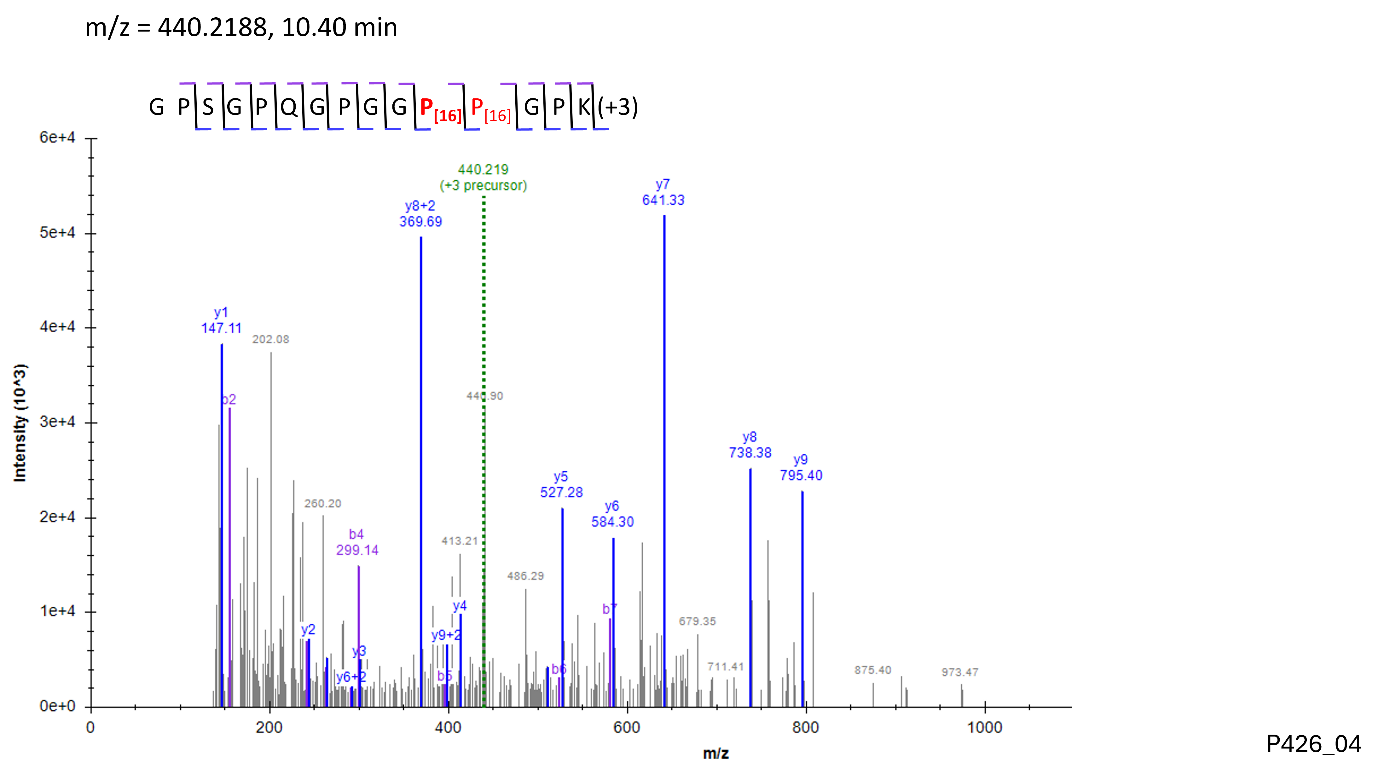

Site: P^459^

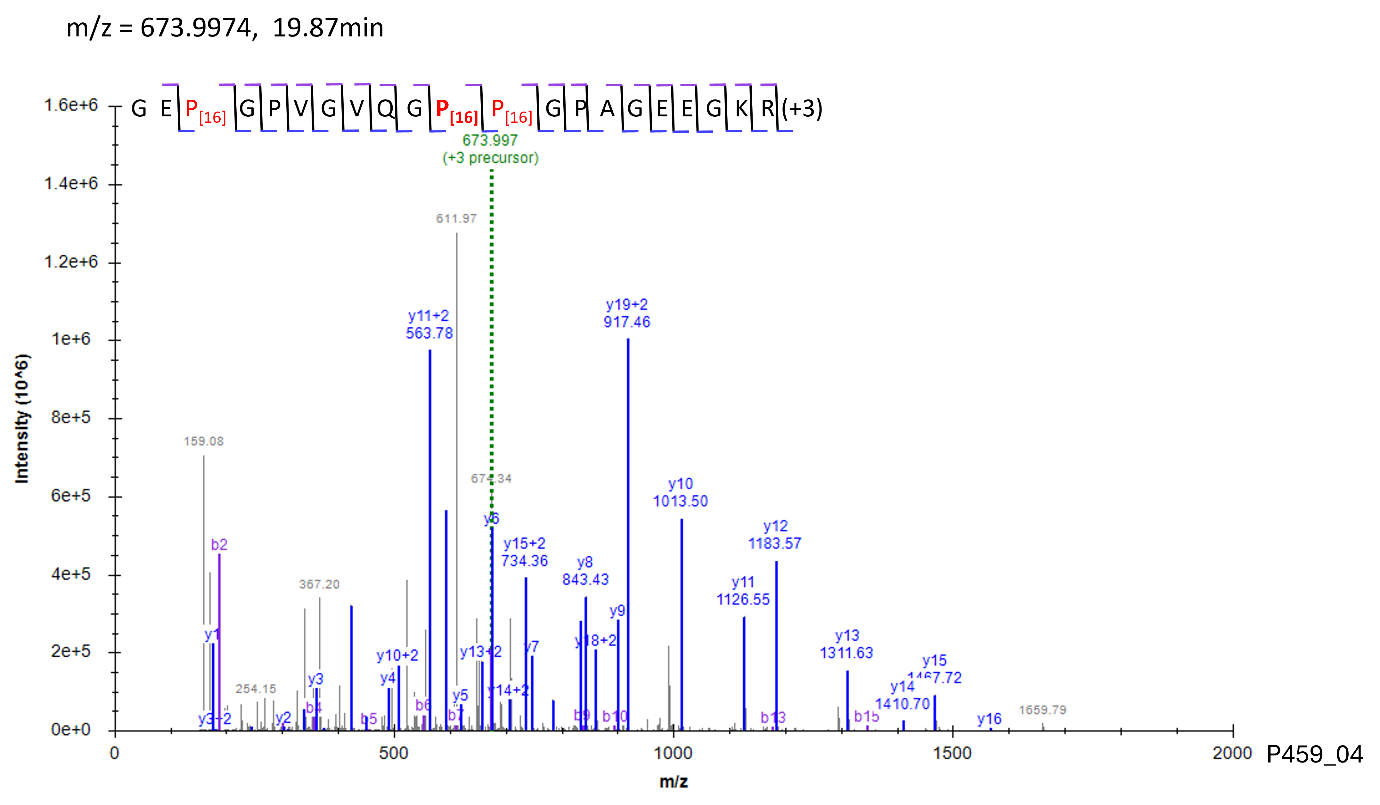

Site: P^555^, P^567^

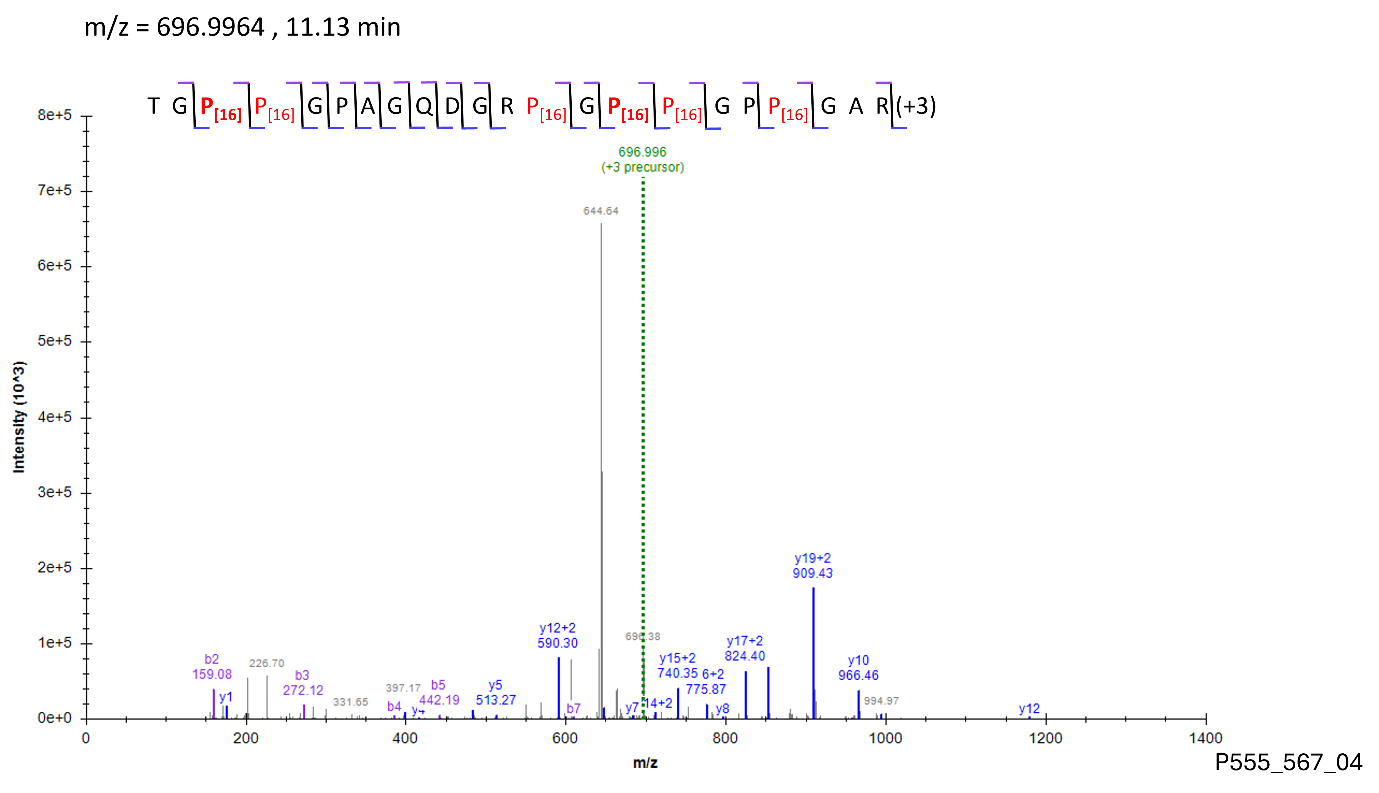

Site: P^570^

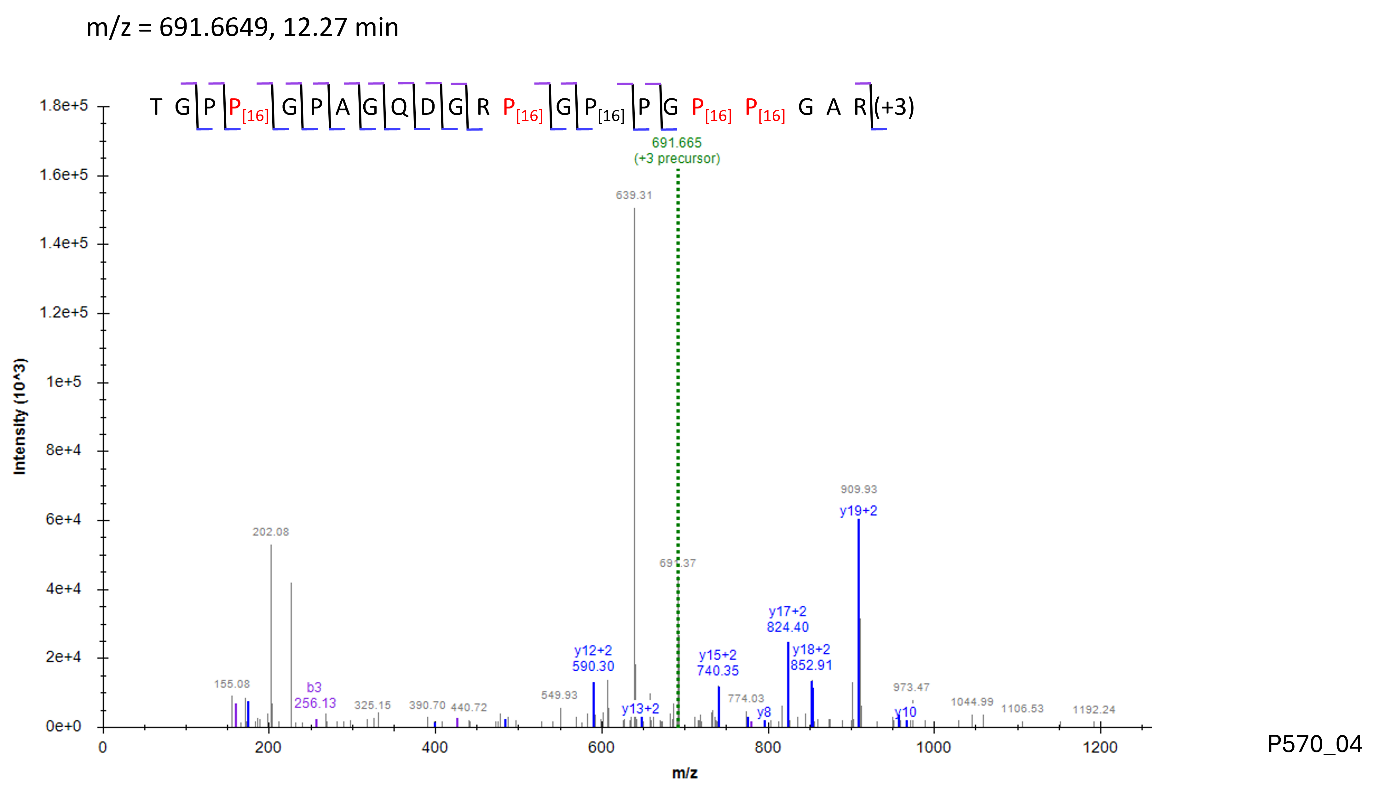

Site: P^603^

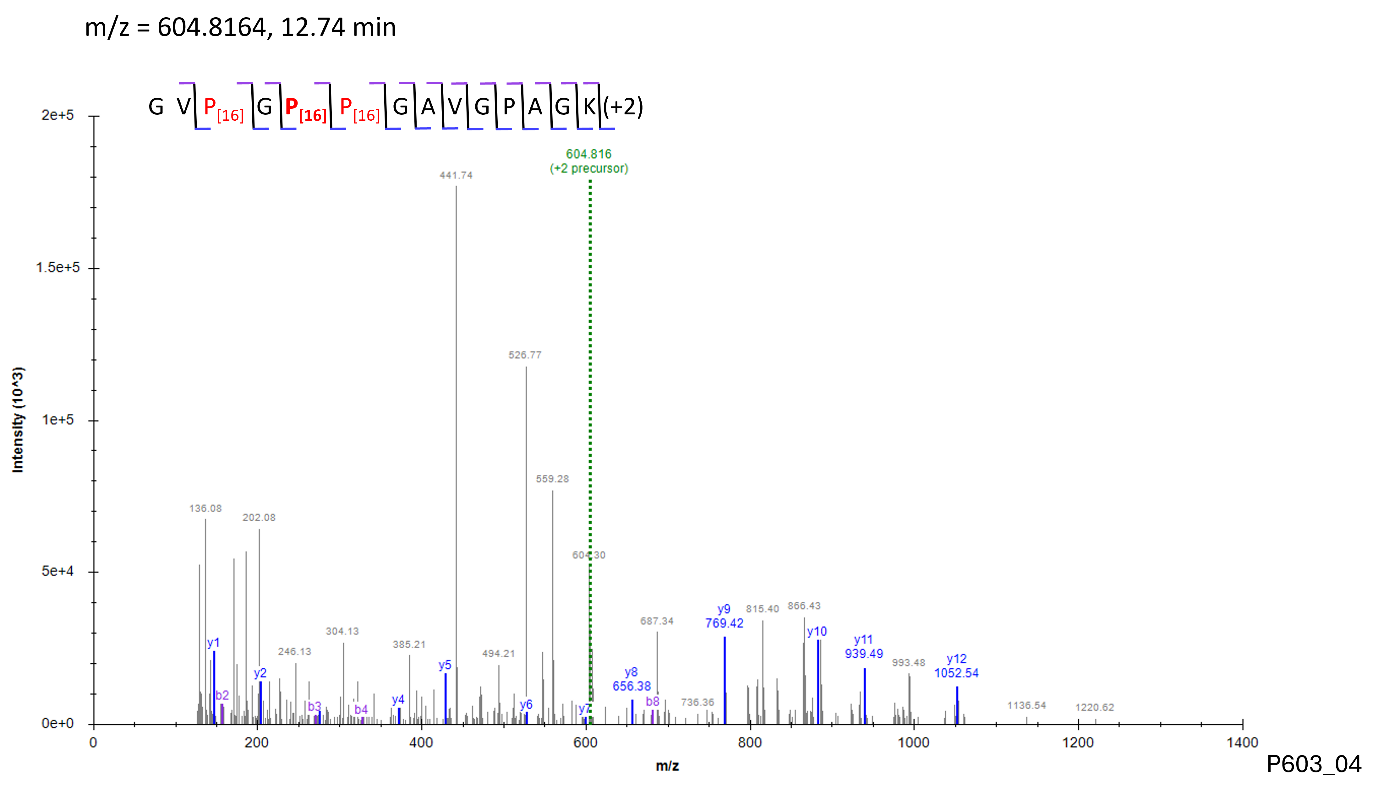

Site: P^690^

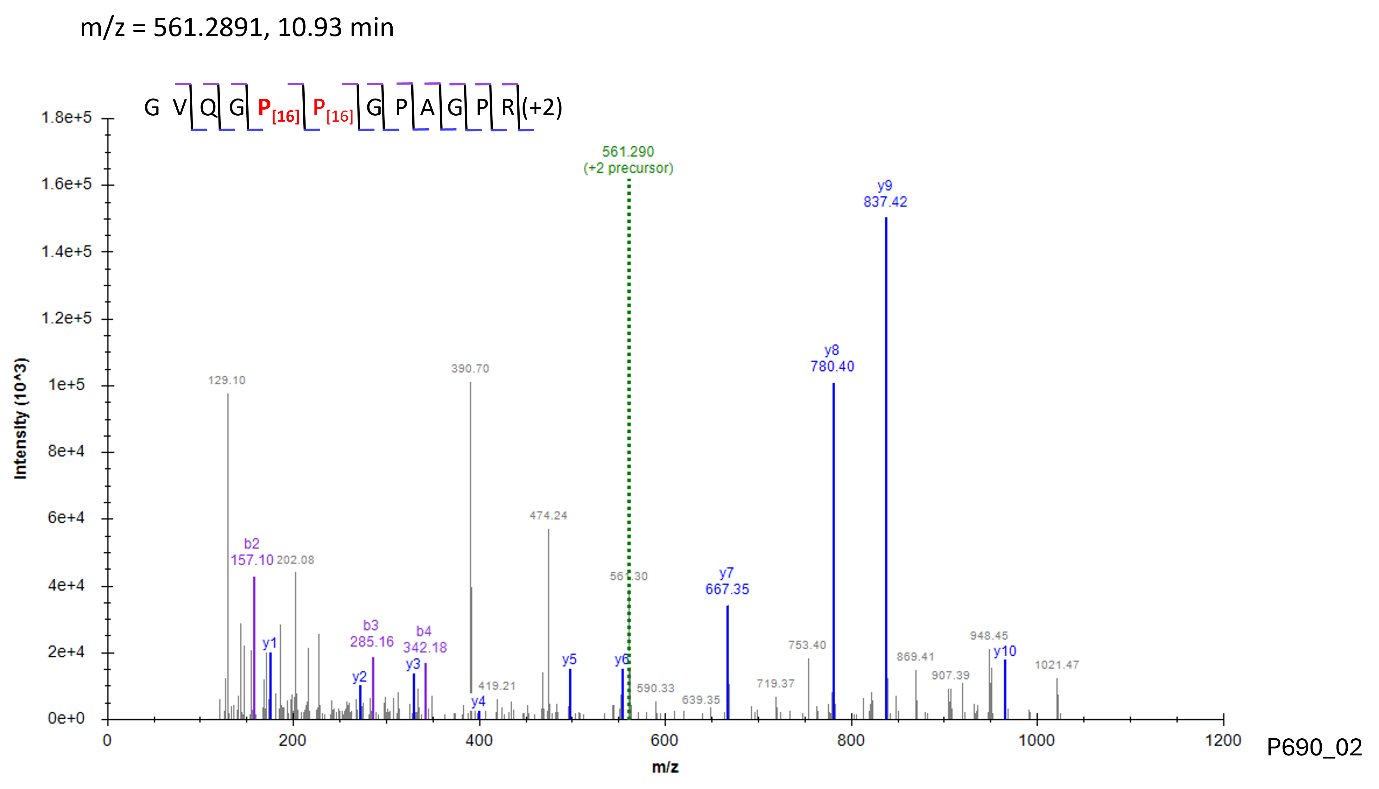

Site: P^771^

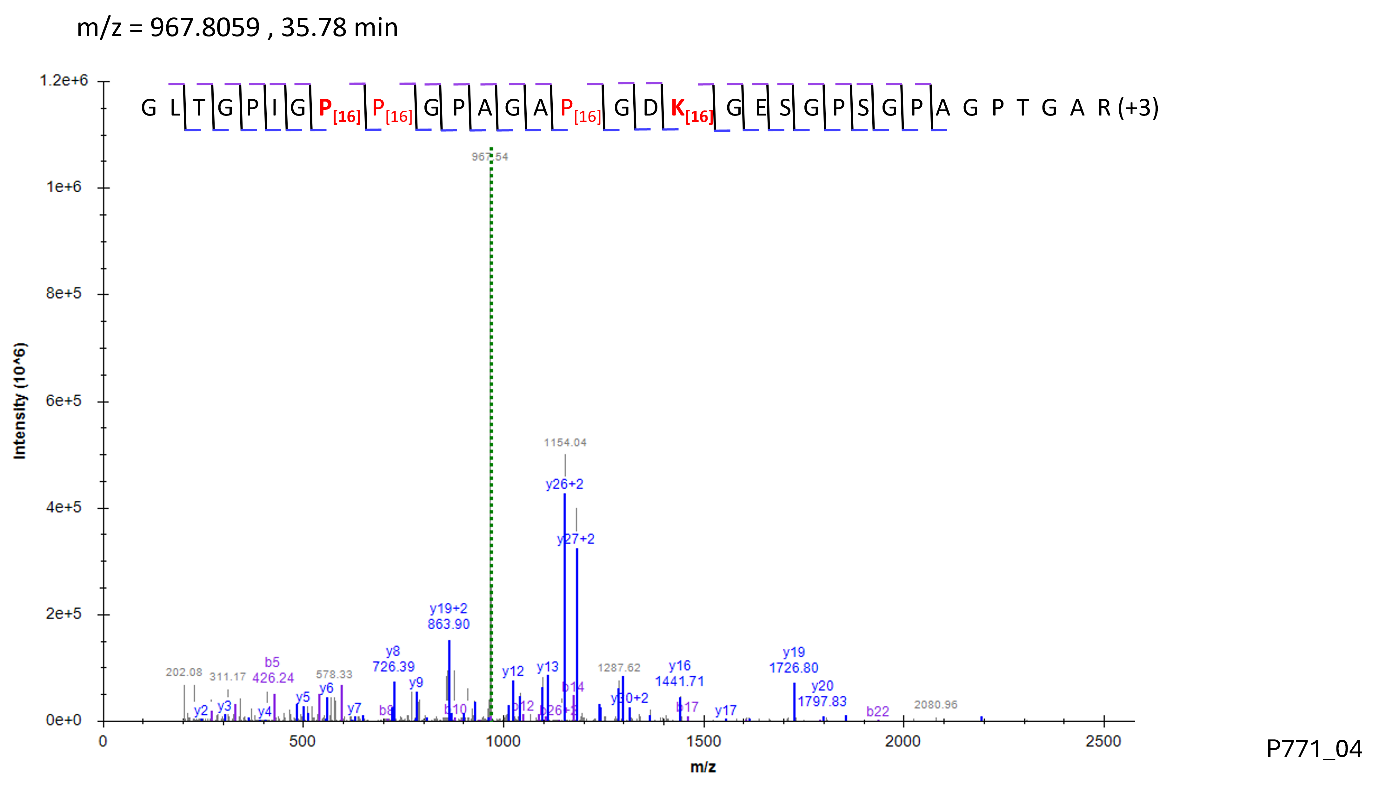

Site: P^807^

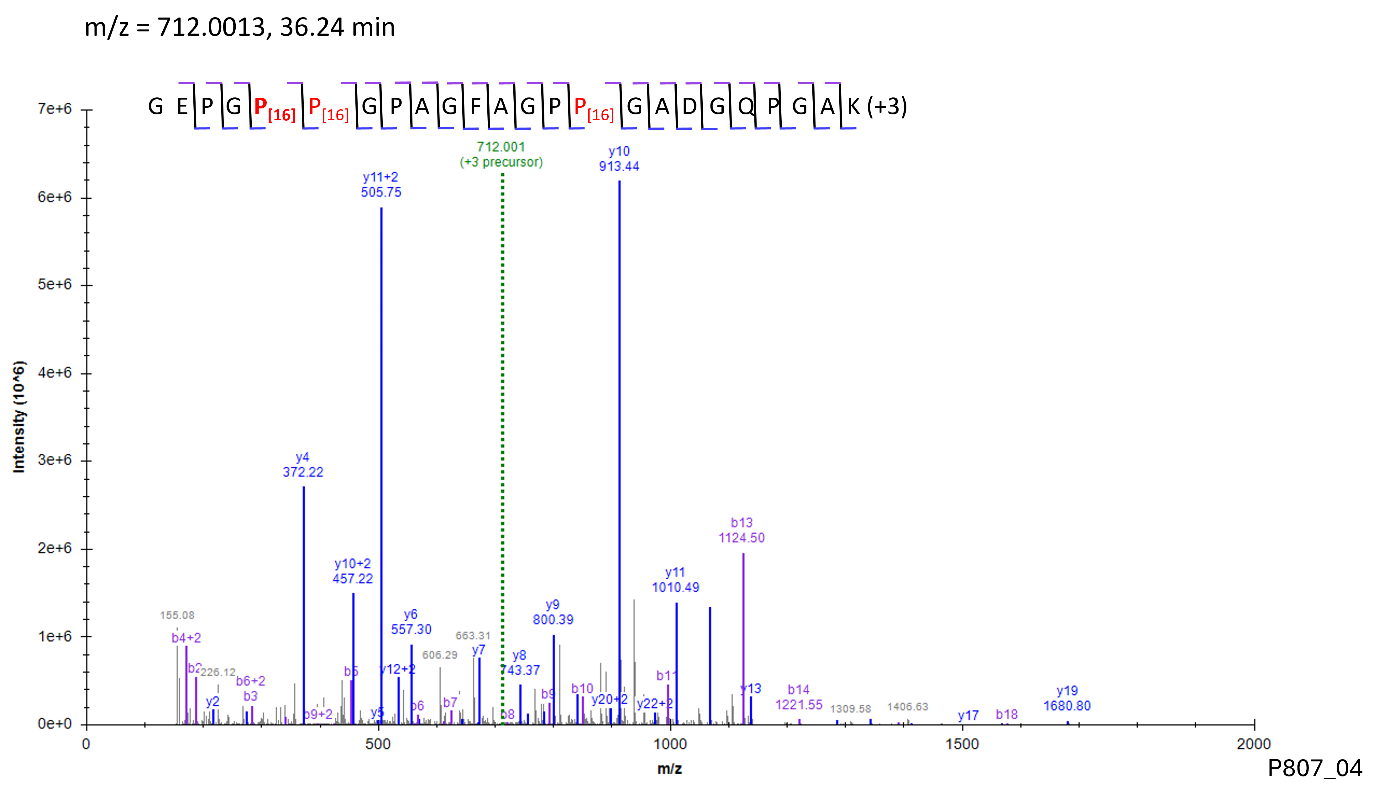

Site: P^816^

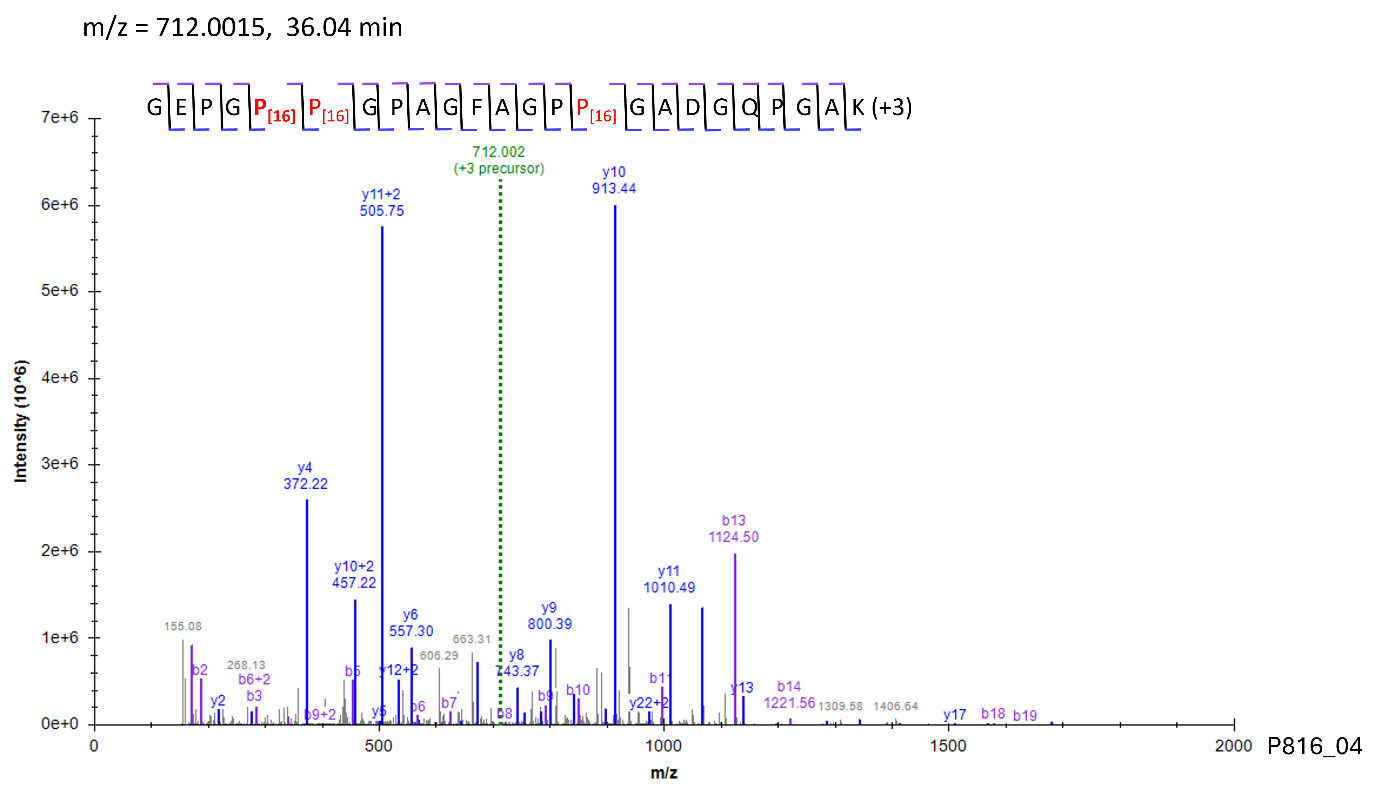

Site: P^840^

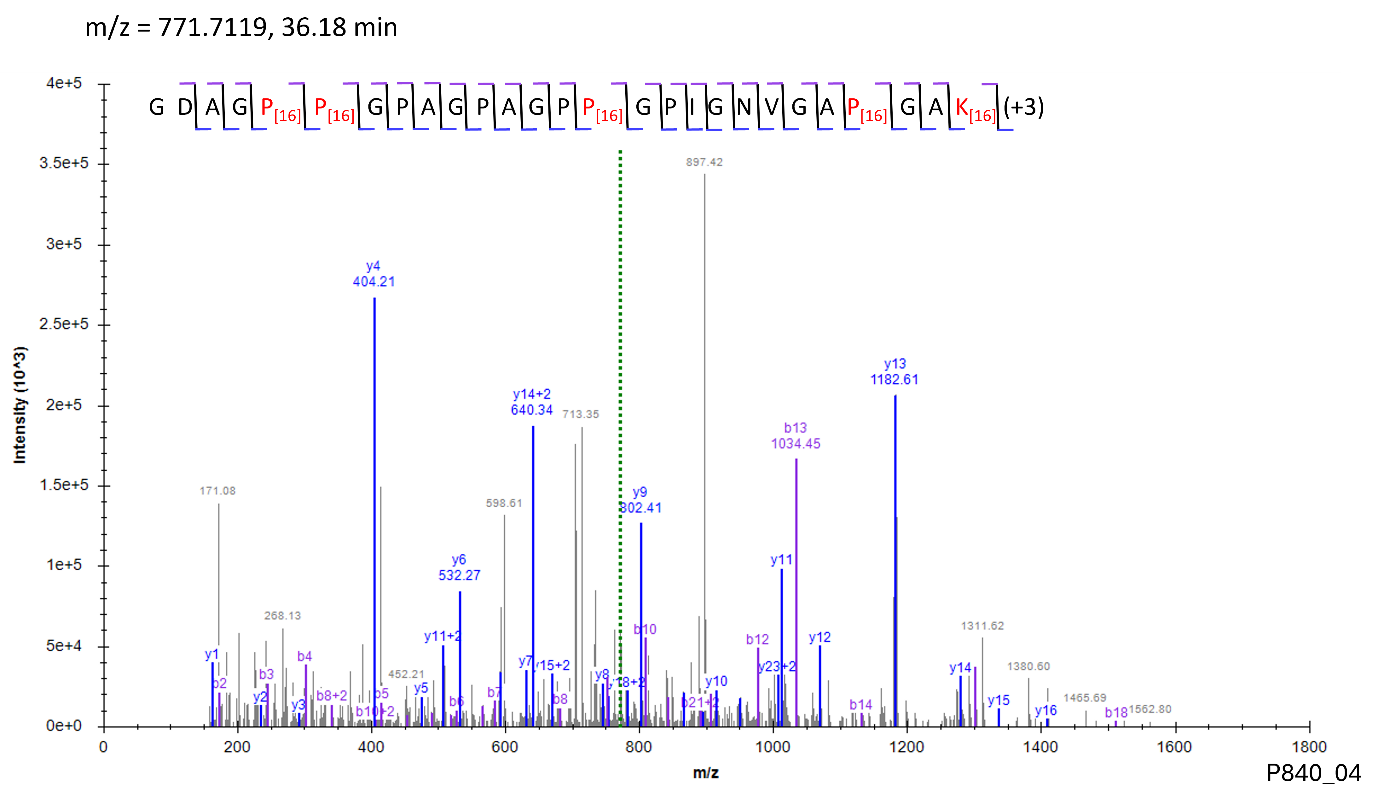

Site: P^870^

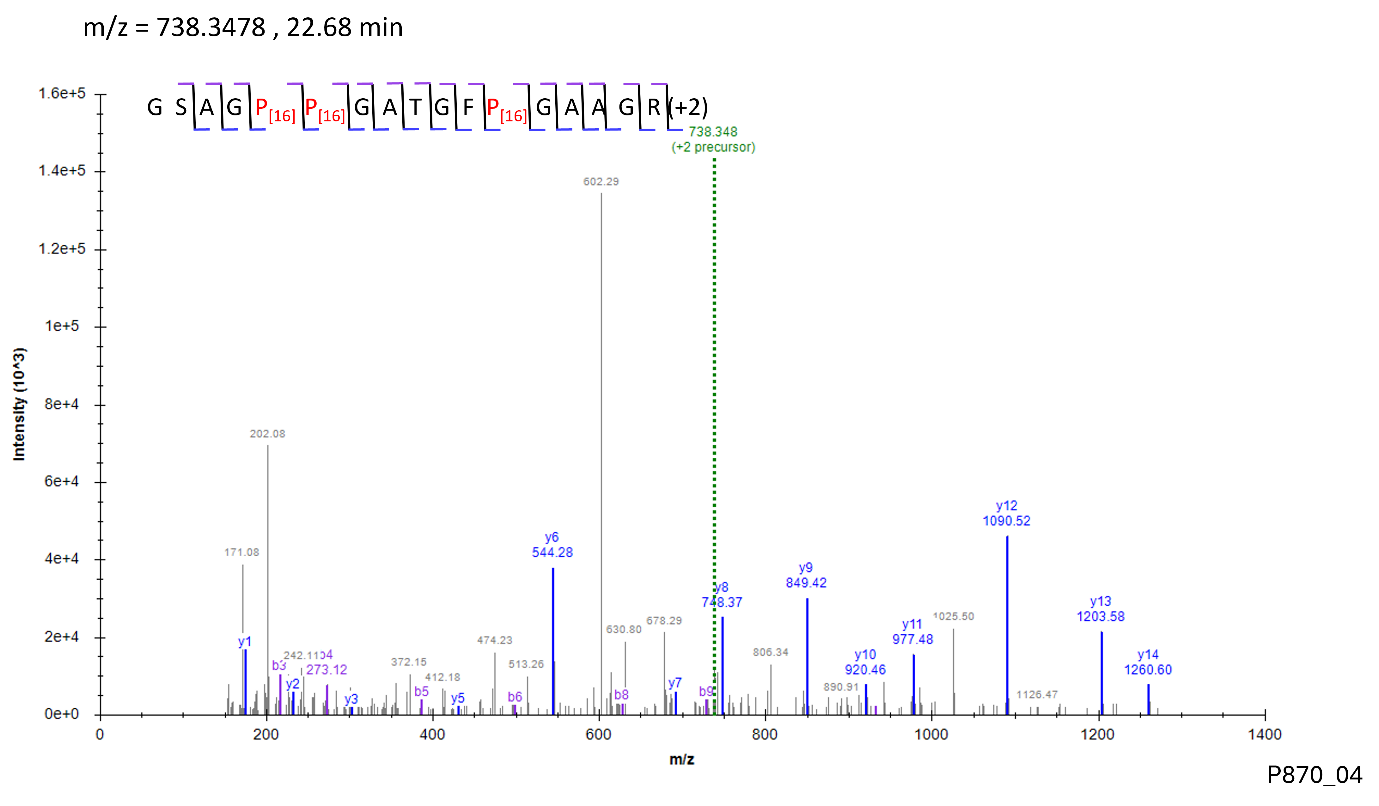

Site: P^897^

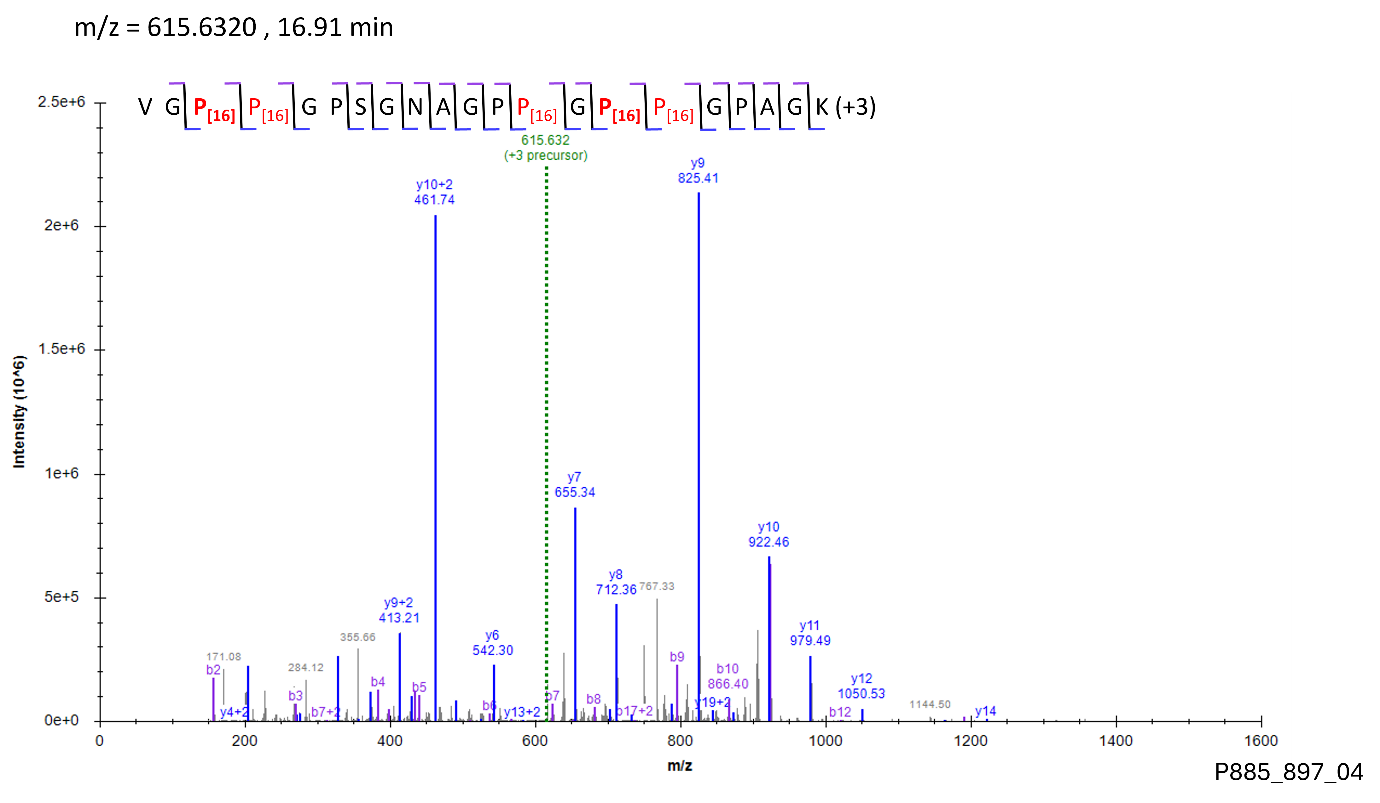

Site: P^894^

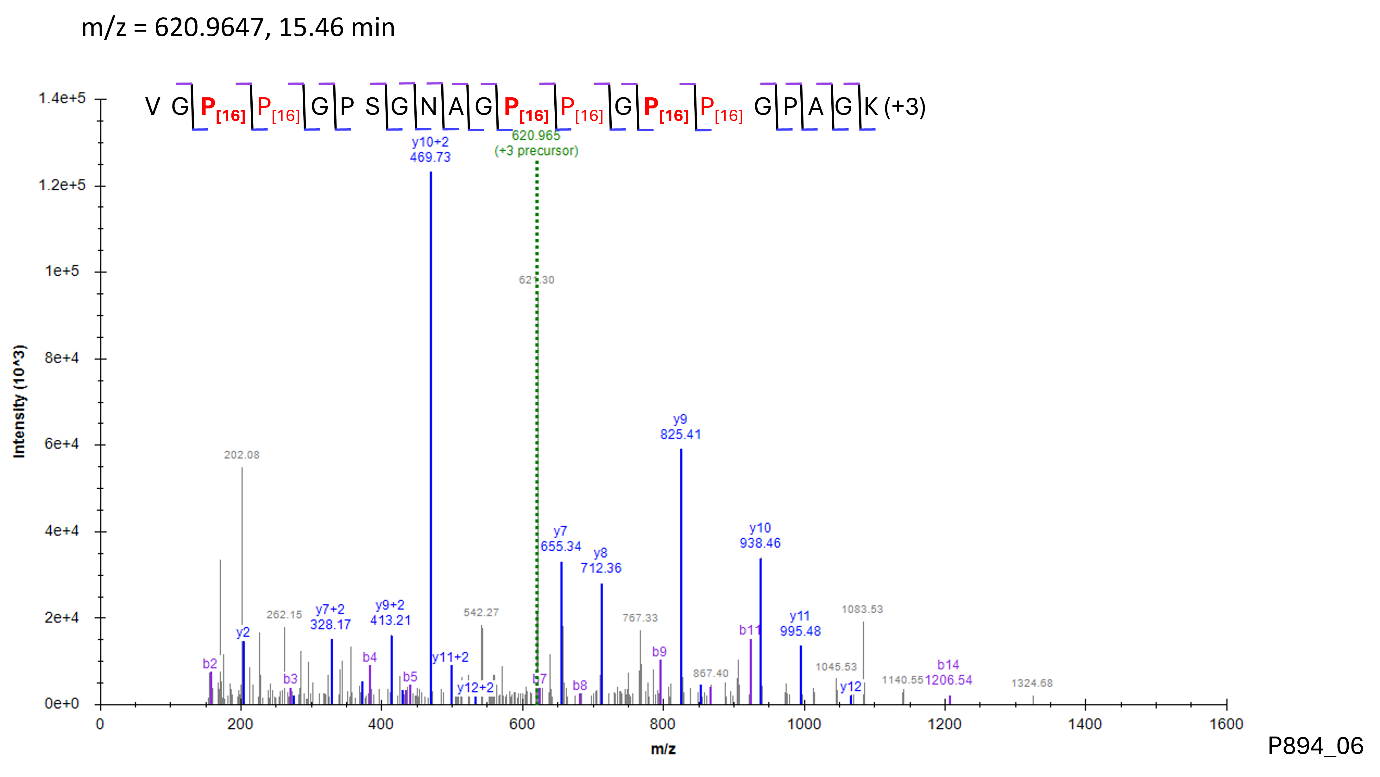

Site: P^927^

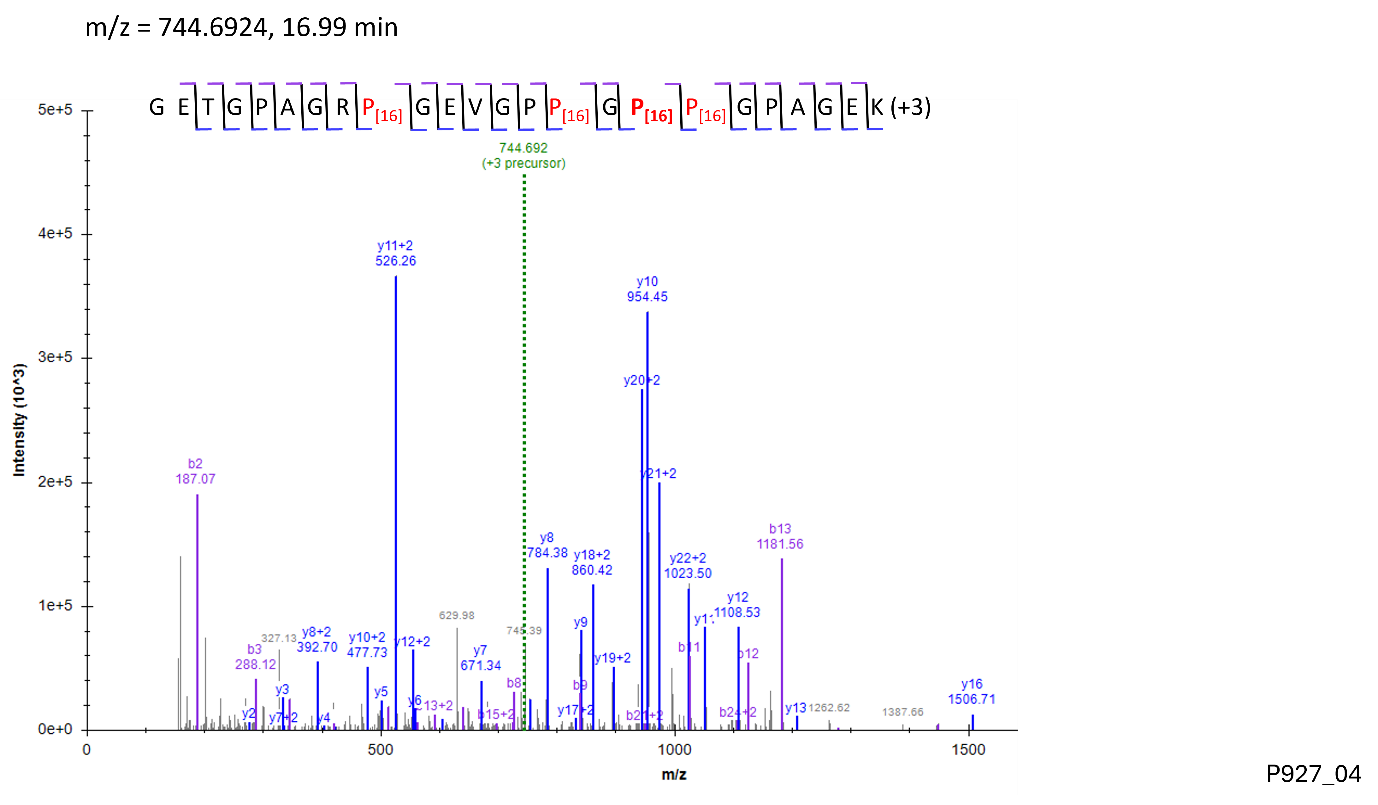

Site: P^996^

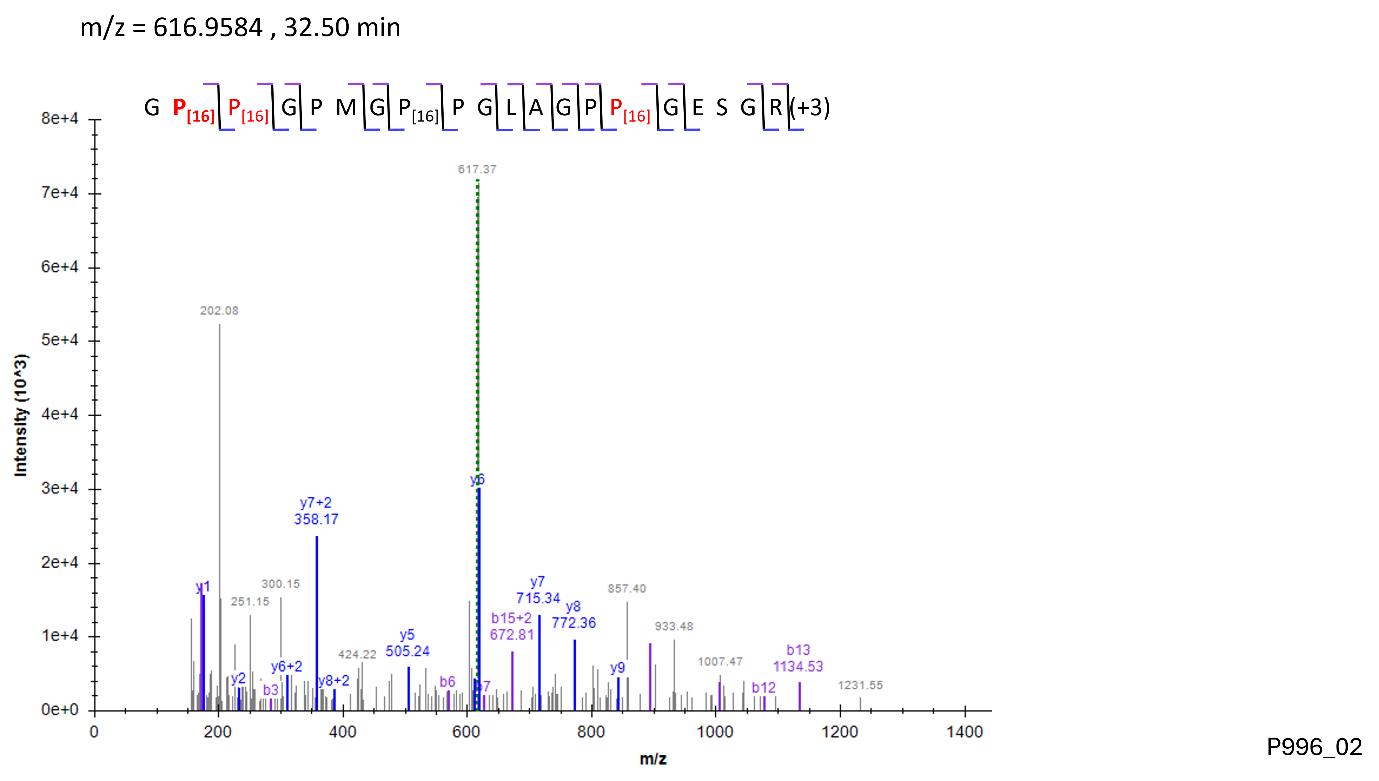

Site: P^1044^

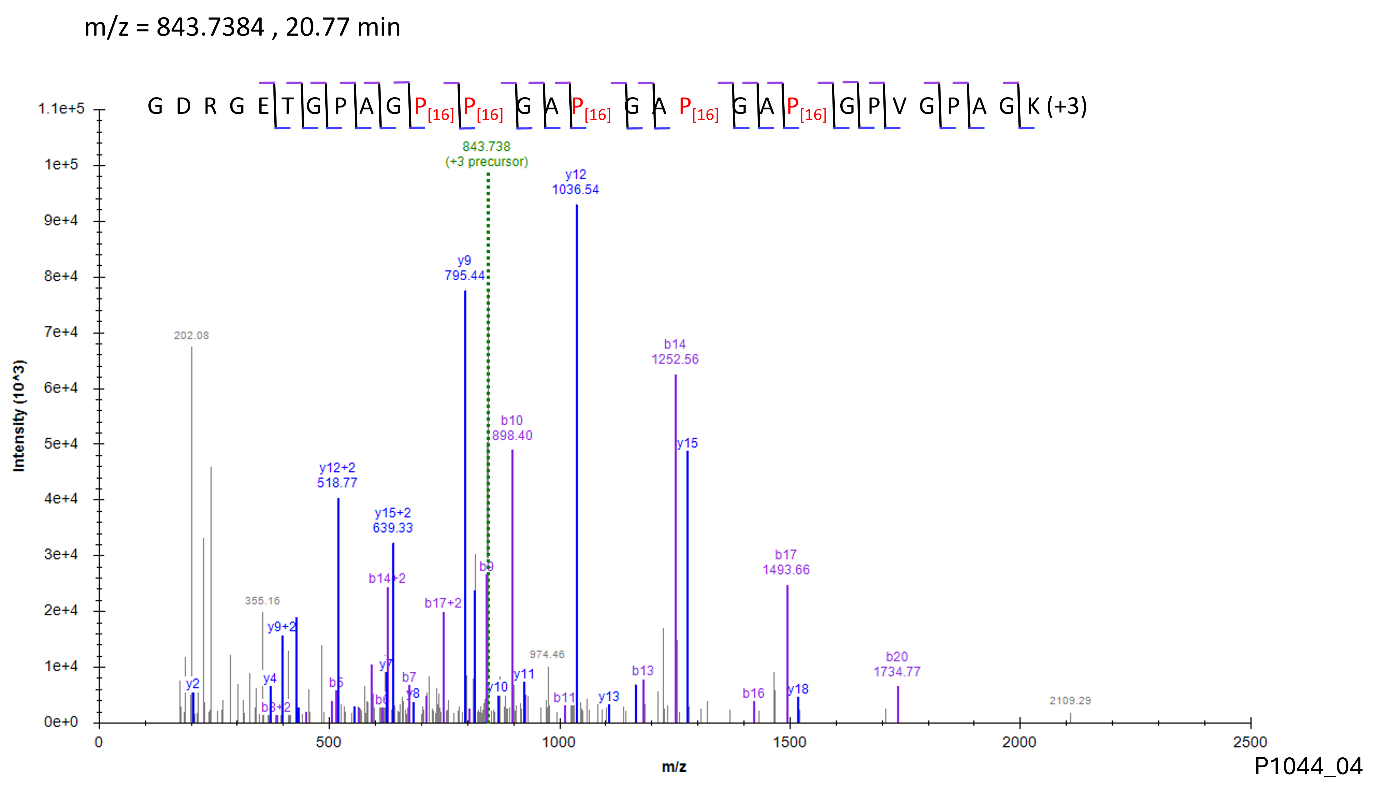

Site: P^1119^, P^1122^

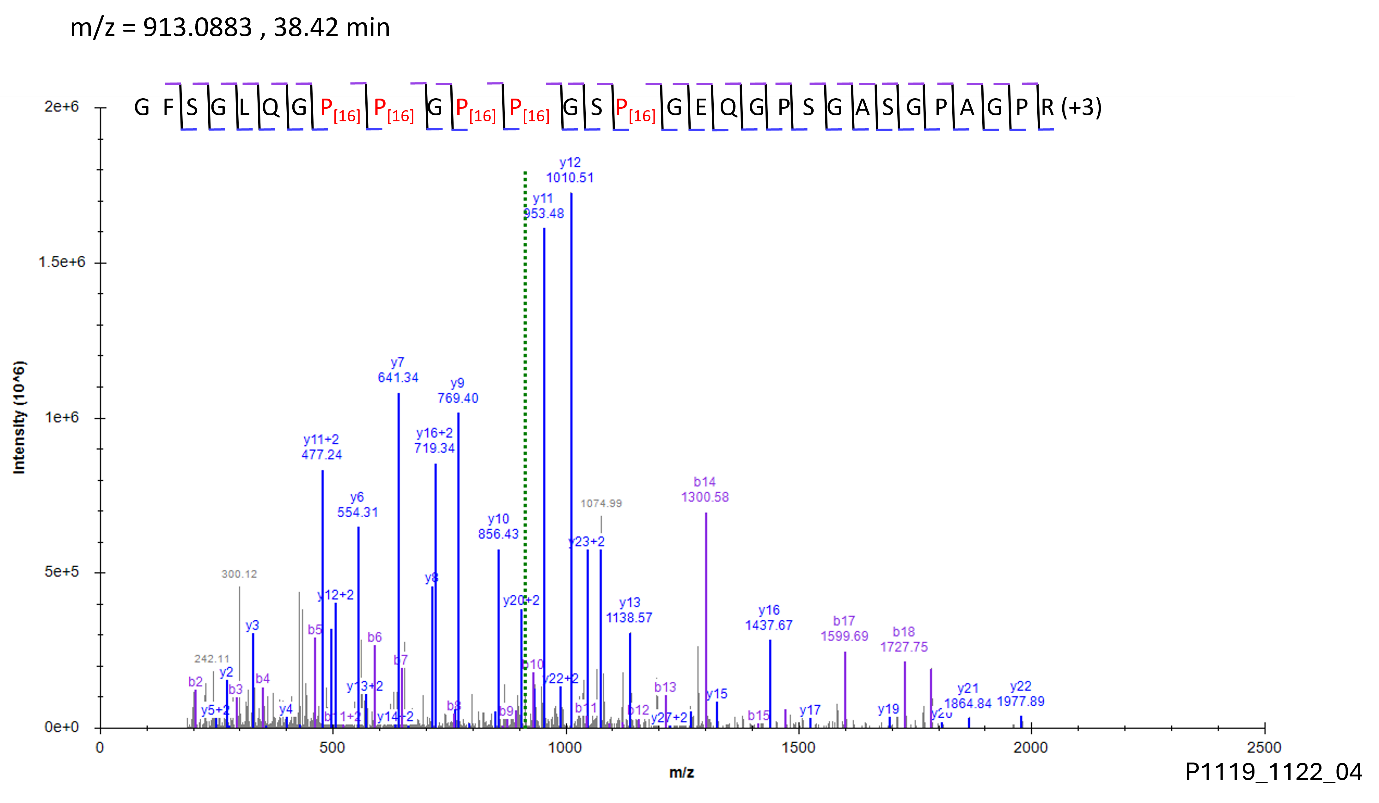

**Modification: HyK**

Site: K^277^

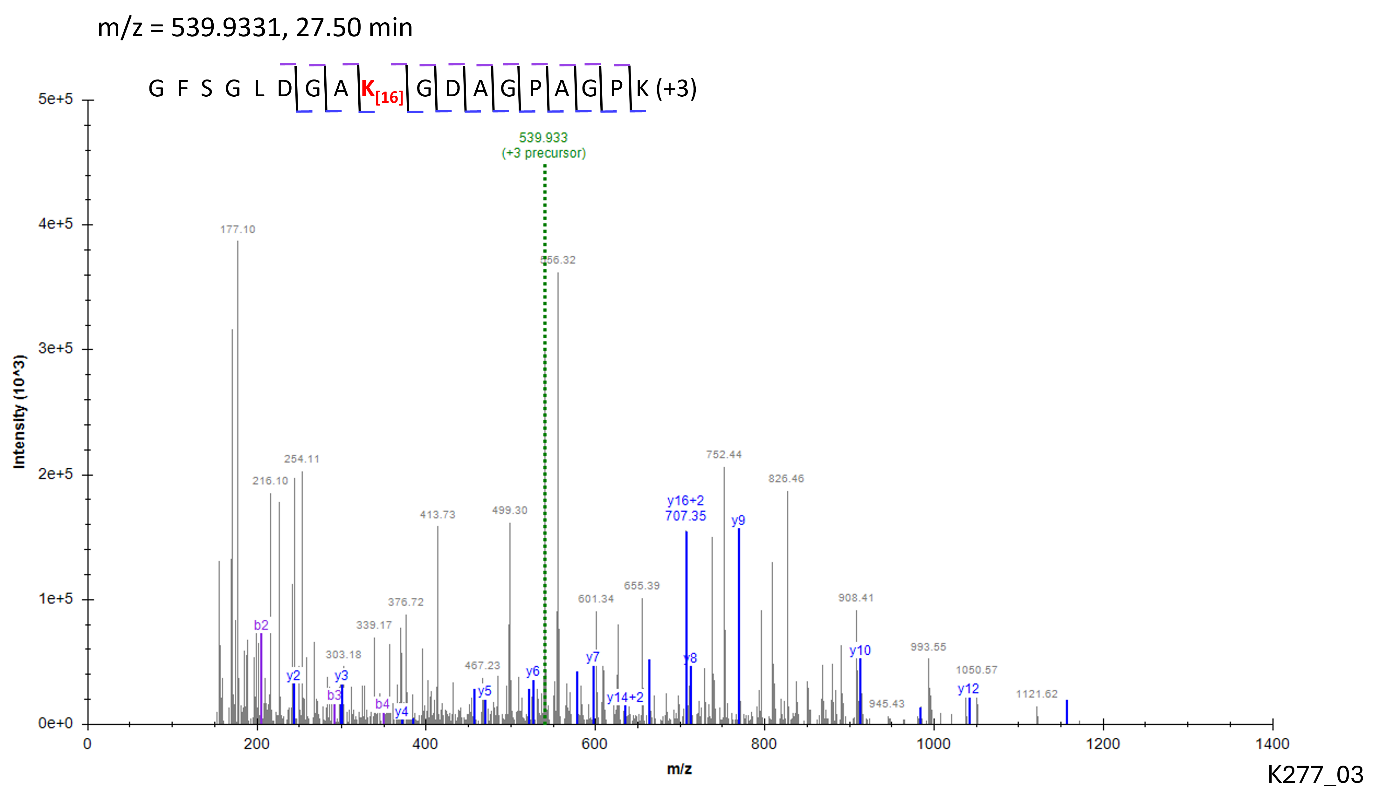

Site: K^286^

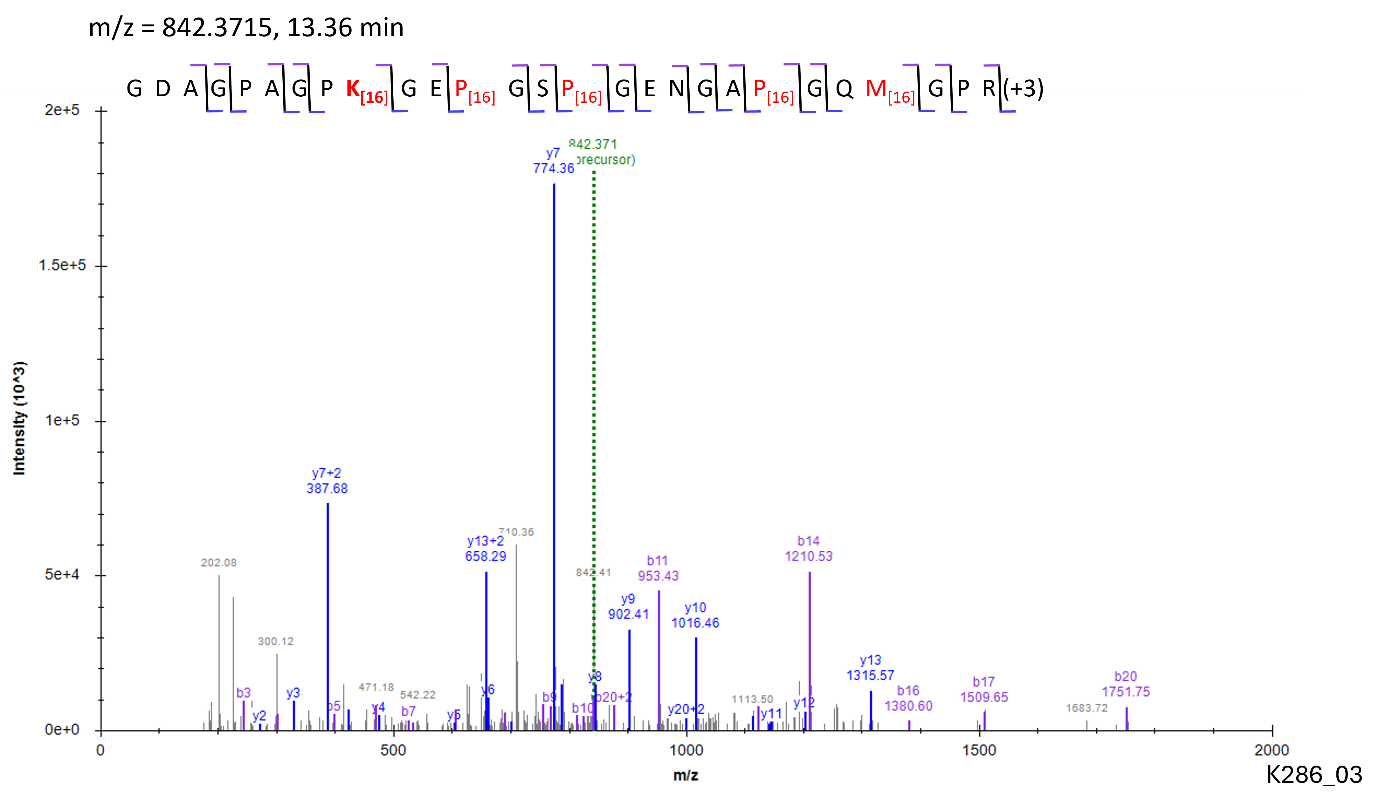

Site: K^397^

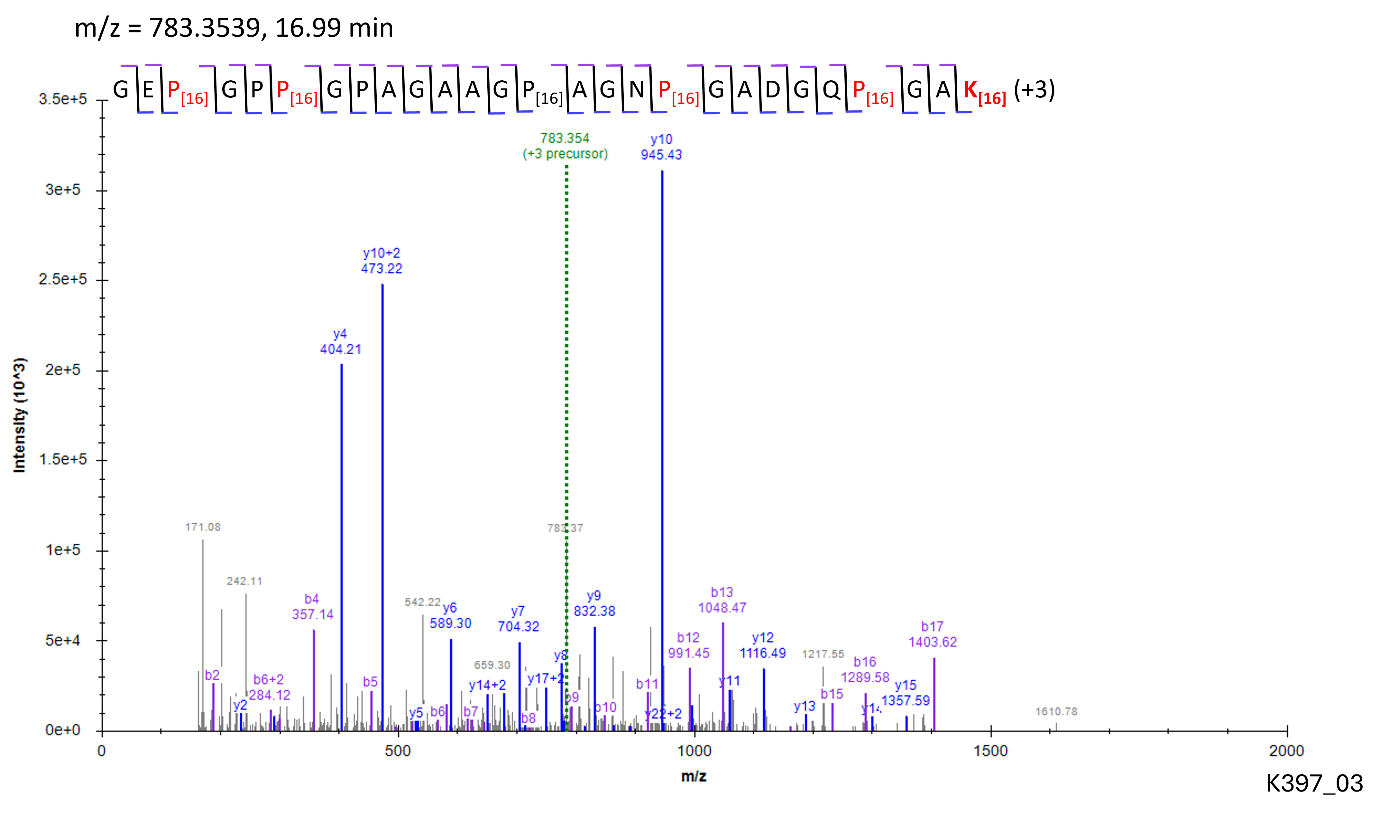

Site: K^448^

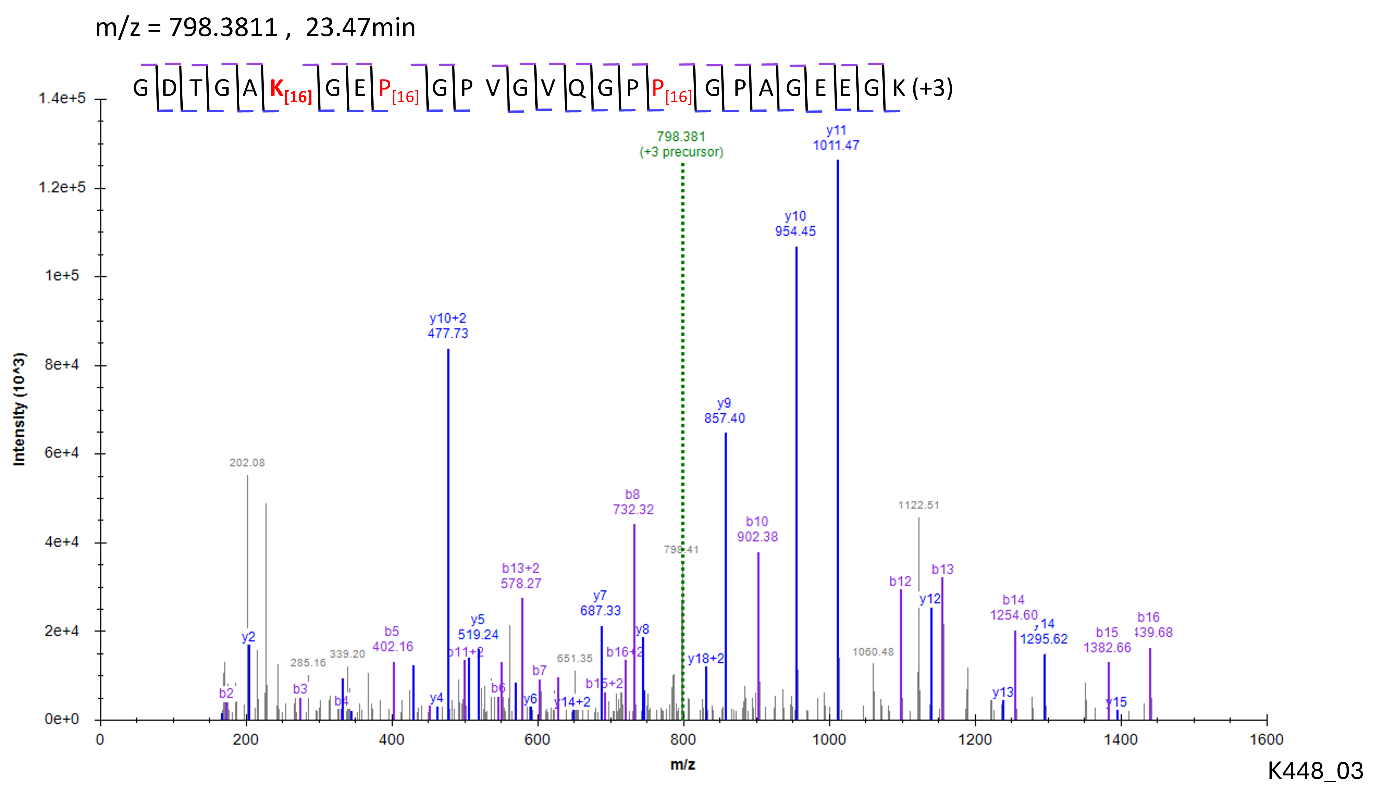

Site: K^538^

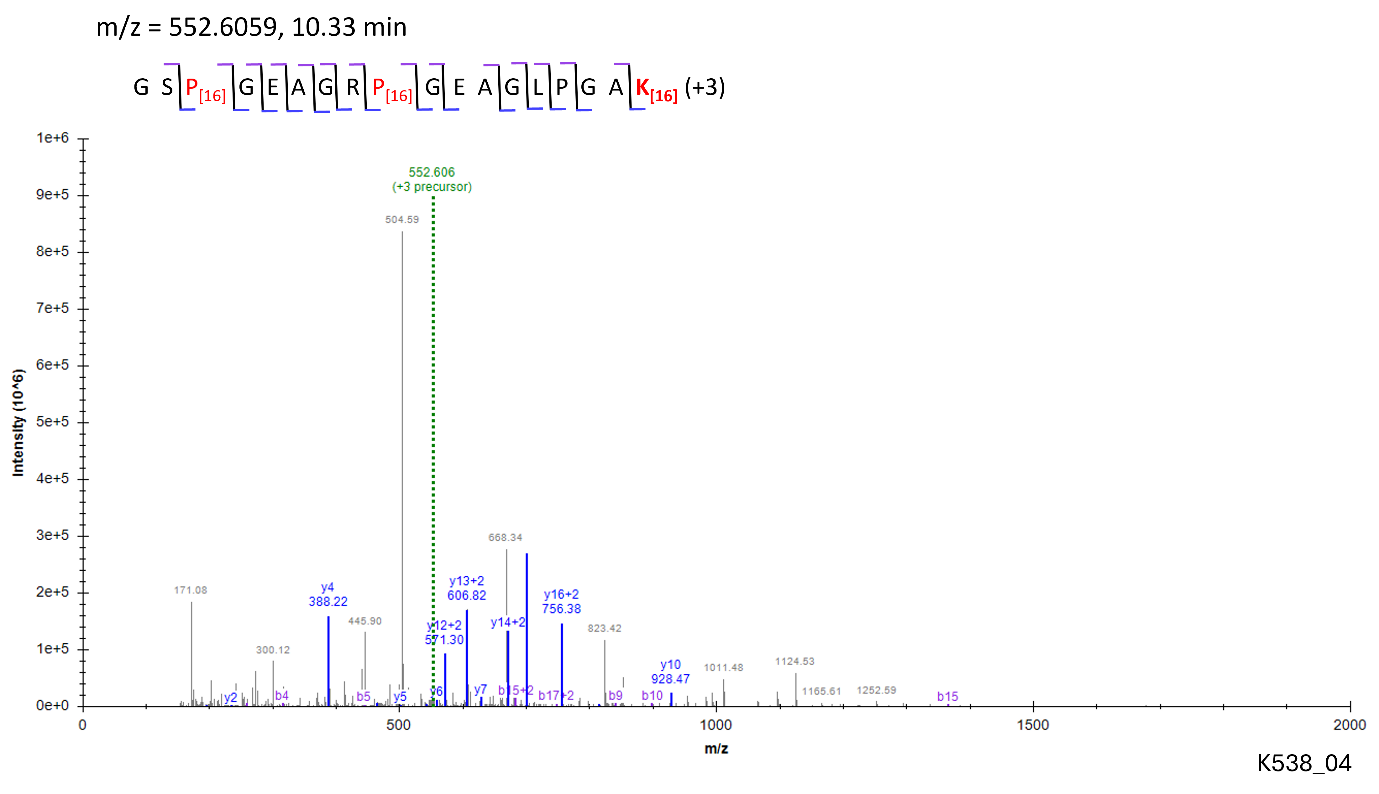

Site: K^586^

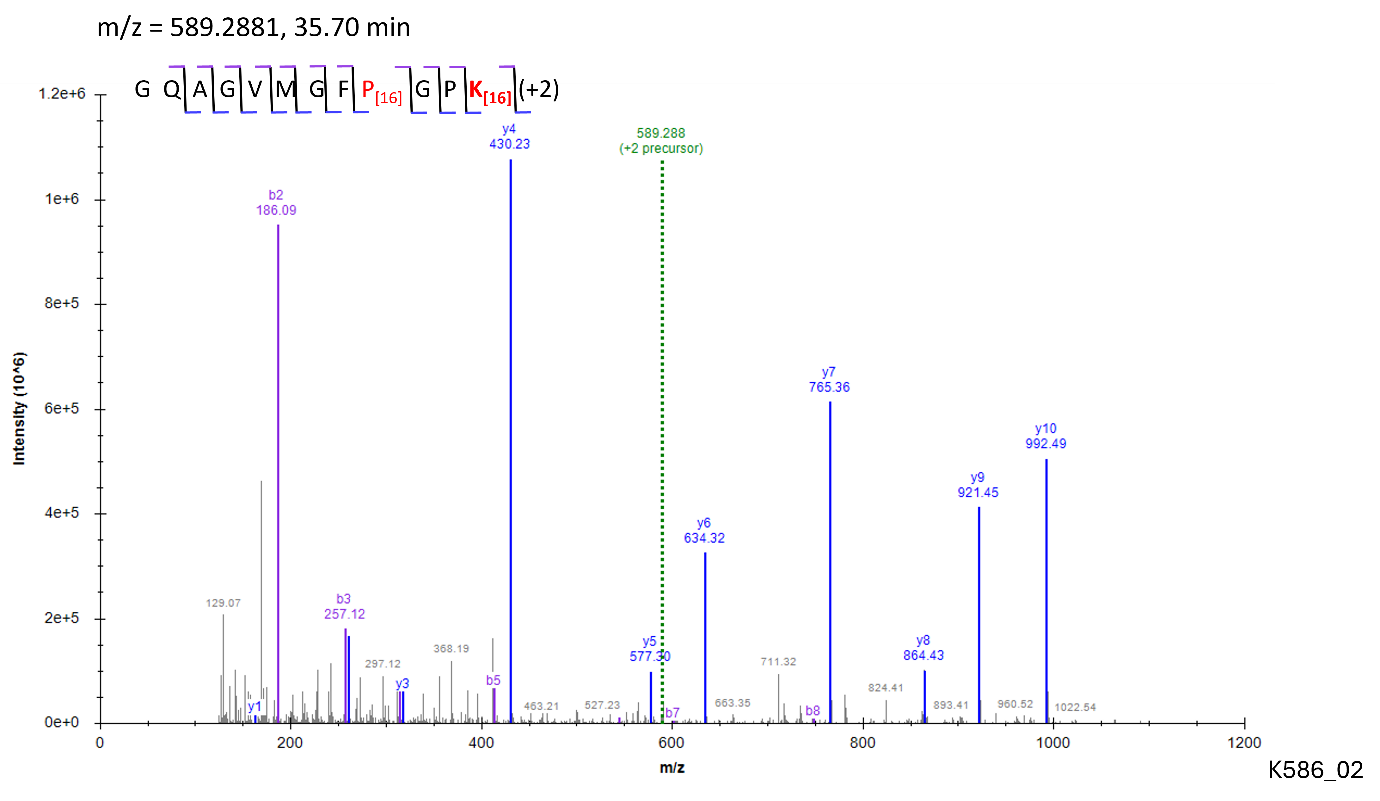

Site: K^781^

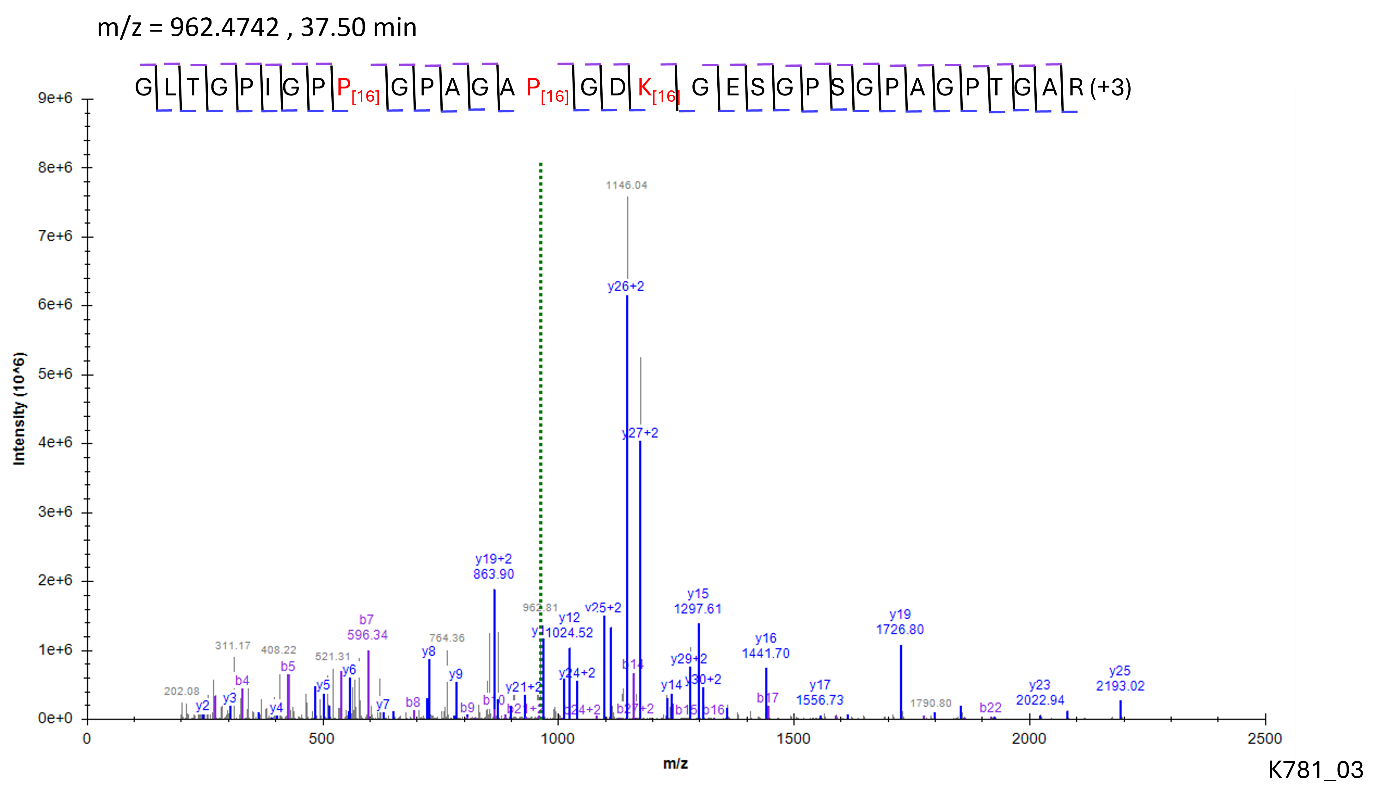

Site: K^826^

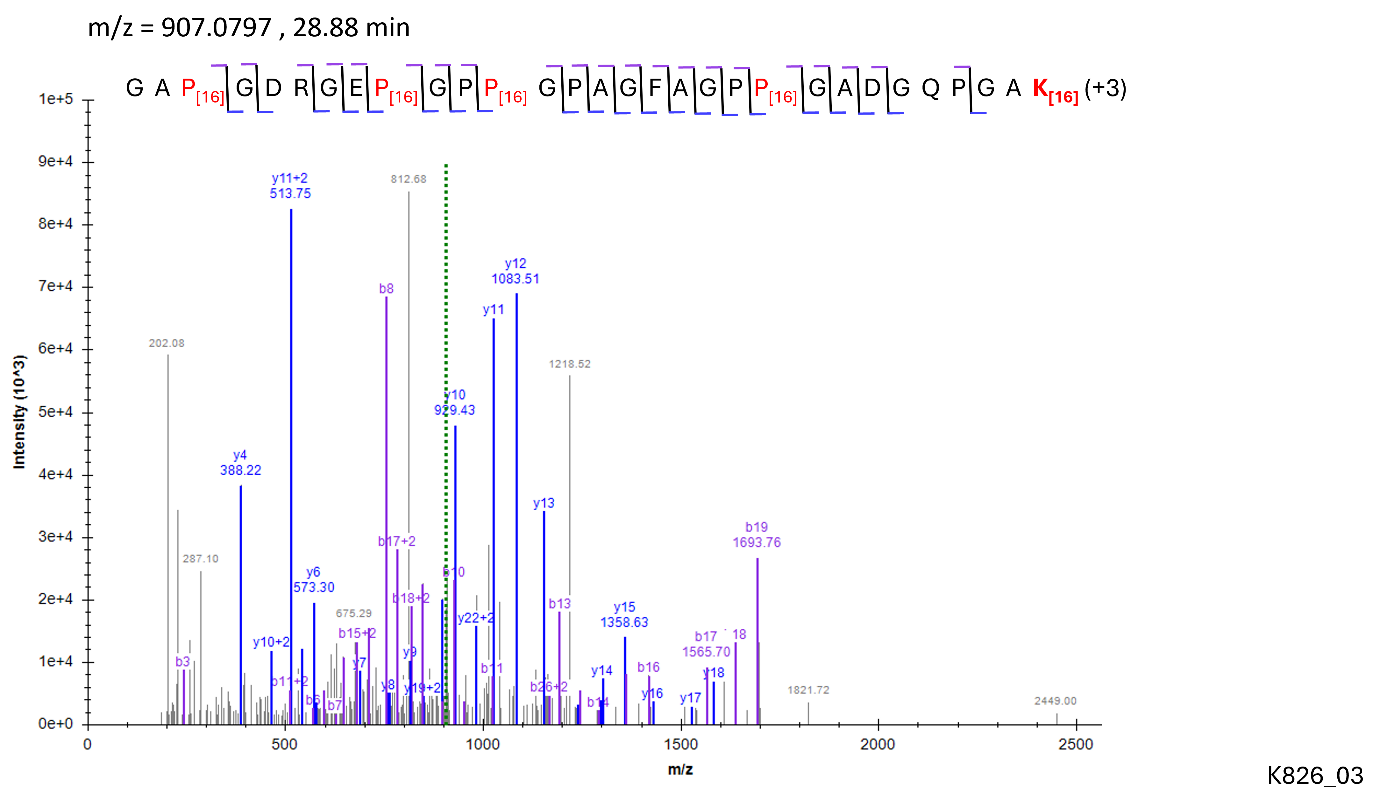

Site: K^862^

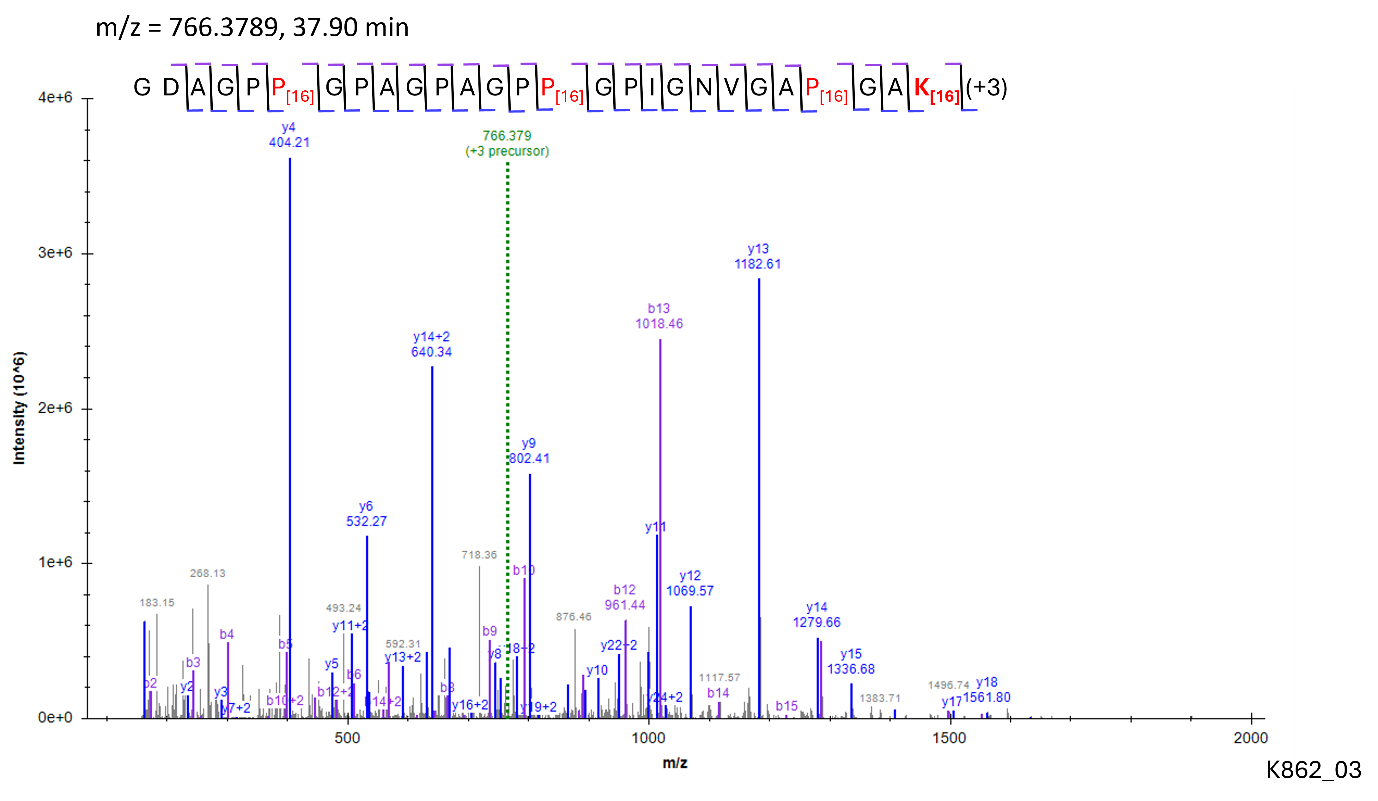

Site: K^934^

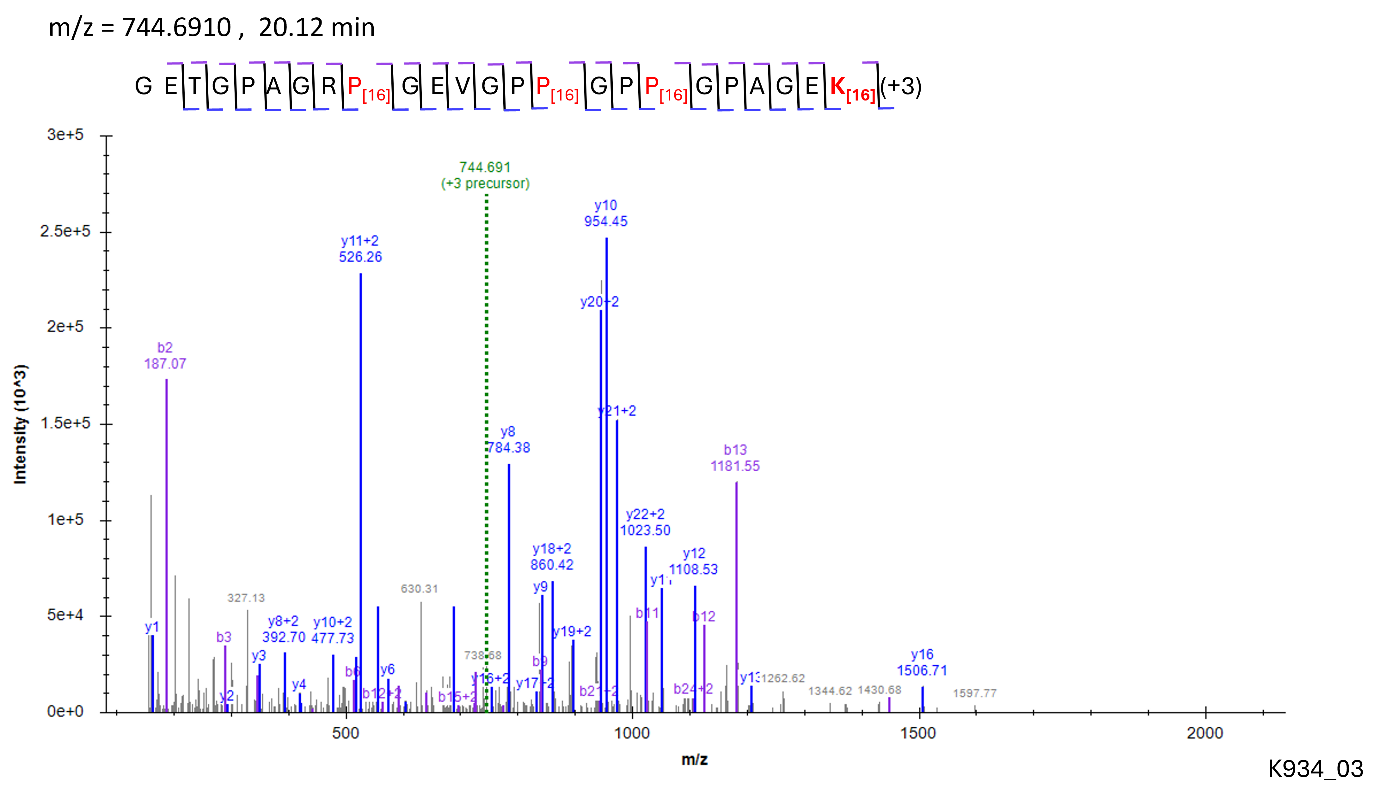

**Modification: G-HyK**

Site: K^448^

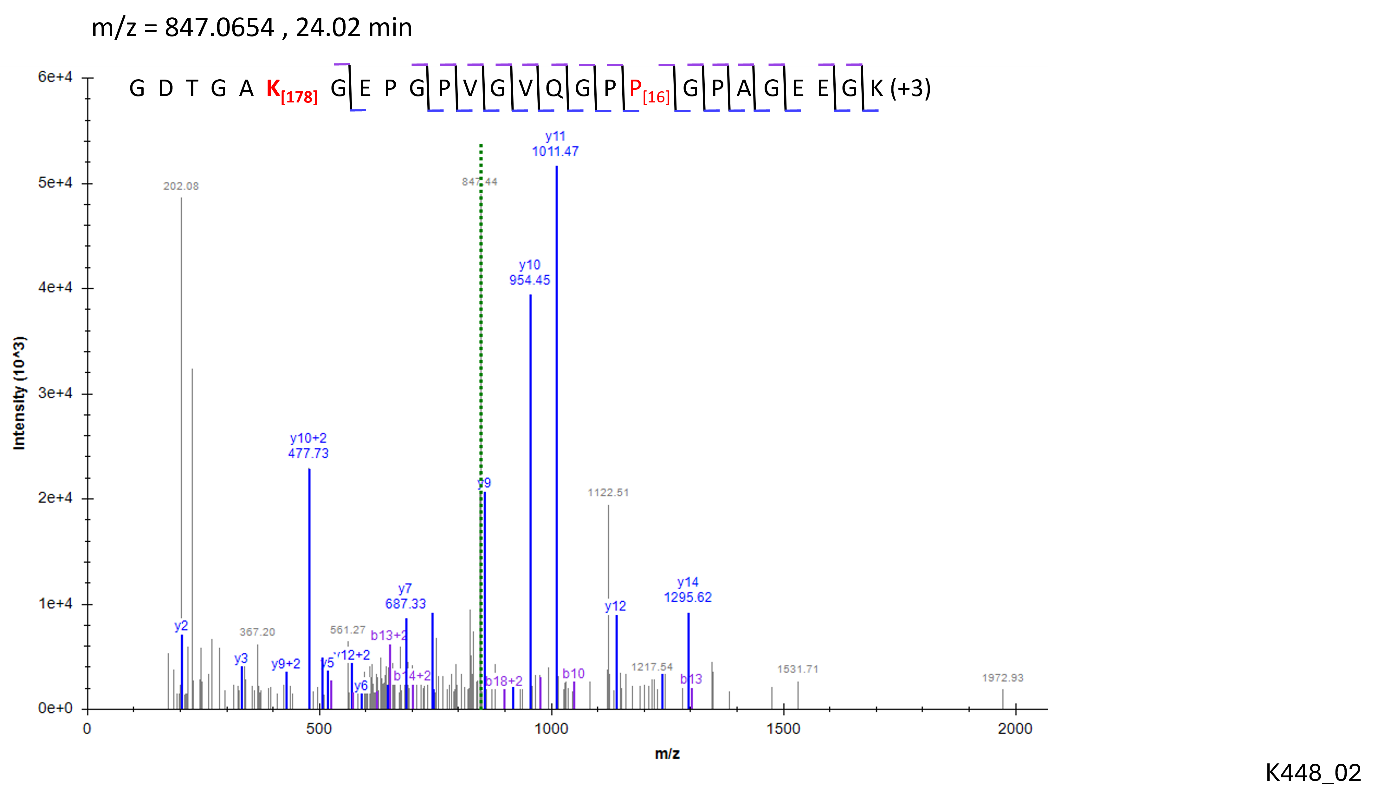

Site: K^862^

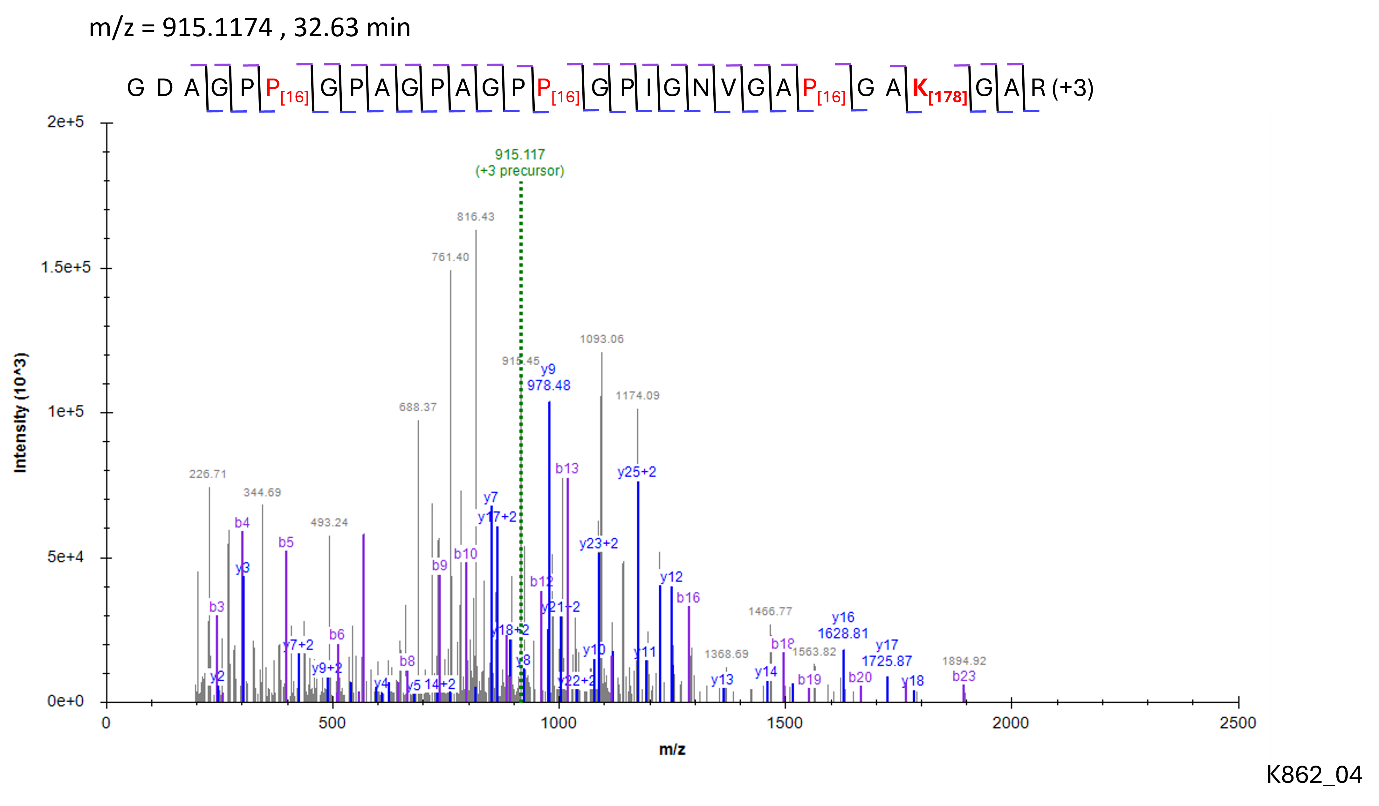

**Modification: GG-HyK**

Site: K^862^

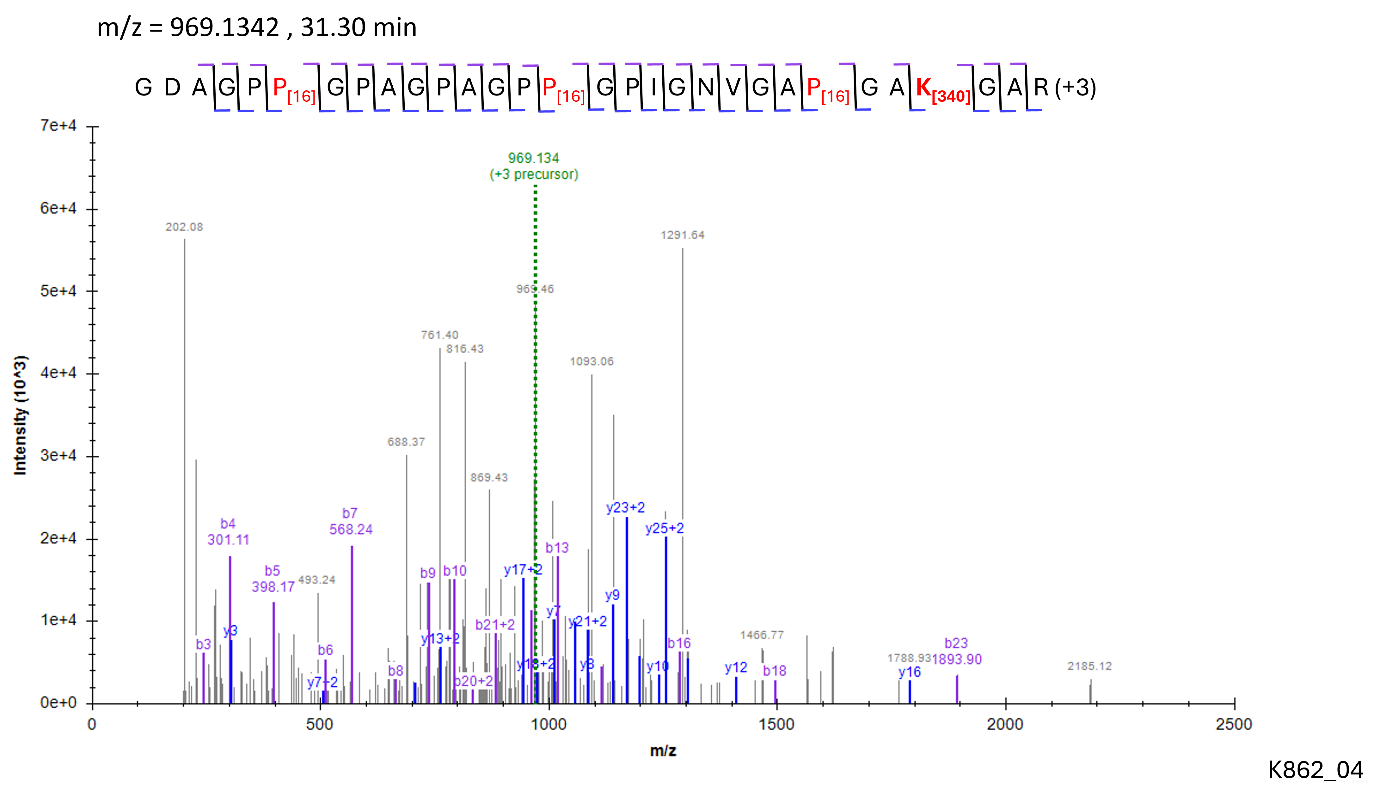
